# Genome wide analysis of gene dosage in 24,092 individuals shows that 10,000 genes modulate cognitive ability

**DOI:** 10.1101/2020.04.03.024554

**Authors:** Guillaume Huguet, Catherine Schramm, Elise Douard, Tamer Petra, Antoine Main, Pauline Monin, Jade England, Khadije Jizi, Thomas Renne, Myriam Poirier, Sabrina Nowak, Charles-Olivier Martin, Nadine Younis, Inga Sophia Knoth, Martineau Jean-Louis, Zohra Saci, Maude Auger, Frédérique Tihy, Géraldine Mathonnet, Catalina Maftei, France Léveillé, David Porteous, Gail Davies, Paul Redmond, Sarah E. Harris, W. David Hill, Emmanuelle Lemyre, Gunter Schumann, Thomas Bourgeron, Zdenka Pausova, Tomas Paus, Sherif Karama, Sarah Lippe, Ian J. Deary, Laura Almasy, Aurélie Labbe, David Glahn, Celia M.T. Greenwood, Sébastien Jacquemont

**Author notes:** **Corresponding authors:** Guillaume Huguet, Sainte Justine University Hospital, 3175 chemin de la Côte-Sainte-Catherine, Montréal, QC H3T 1C5, Sébastien Jacquemont, Sainte Justine University Hospital, 3175 chemin de la Côte-Sainte-Catherine, Montréal, QC H3T 1C5. Shared first authorship (Guillaume Huguet, Catherine Schramm). **Single sentence summary:** CNVs’ effect-sizes on intelligence are predicted using measures of 5 intolerance to haploinsufficiency and are distributed across half of the coding genes.

## Abstract

Genomic Copy Number Variants (CNVs) are routinely identified and reported back to patients with neuropsychiatric disorders, but their quantitative effects on essential traits such as cognitive ability are poorly documented. We have recently shown that the effect-size of deletions on cognitive ability can be statistically predicted using measures of intolerance to haploinsufficiency. However, the effect-sizes of duplications remain unknown. It is also unknown if the effect of multigenic CNVs are driven by a few genes intolerant to haploinsufficiency or distributed across tolerant genes as well.

Here, we identified all CNVs >50 kilobases in 24,092 individuals from unselected and autism cohorts with assessments of general intelligence. Statistical models used measures of intolerance to haploinsufficiency of genes included in CNVs to predict their effect-size on intelligence. Intolerant genes decrease general intelligence by 0.8 and 2.6 points of IQ when duplicated or deleted, respectively. Effect-sizes showed no heterogeneity across cohorts. Validation analyses demonstrated that models could predict CNV effect-sizes with 78% accuracy. Data on the inheritance of 27,766 CNVs showed that deletions and duplications with the same effect-size on intelligence occur *de novo* at the same frequency.

We estimated that around 10,000 intolerant and tolerant genes negatively affect intelligence when deleted, and less than 2% have large effect-sizes. Genes encompassed in CNVs were not enriched in any GOterms but gene regulation and brain expression were GOterms overrepresented in the intolerant subgroup. Such pervasive effects on cognition may be related to emergent properties of the genome not restricted to a limited number of biological pathways.

## Introduction

Copy Number Variants (CNVs) are deletions or duplications larger than 1000 base pairs. The contribution of CNVs to the etiology of intellectual disability (ID)[1–3], autism[4–6] and schizophrenia[6–8] is well established. The interpretation of CNVs in research and medical diagnostics remains essentially binary: benign or pathogenic (contributing to mental illness)*[9]*. The routine implementation of Chromosomal Micro-Arrays (CMAs) as a first-tier diagnostic test identifies “pathogenic” CNVs in 10 to 15 % of children with neurodevelopmental disorders (NDD)*[10]*. A binary interpretation is however of limited use because patients present a broad spectrum of cognitive symptoms ranging from severe ID to learning disabilities. The quantitative effects of CNVs are poorly documented even for important traits such as general intelligence. It may be available for the most frequently recurrent CNVs but data is often collected in patients ascertained in the clinic with a bias towards severely affected individuals, leading to potentially gross overestimation of effect size. Only two studies have been conducted in unselected populations [11, 12] showing reduced performance on cognitive test for 24 recurrent CNVs. However, recurrent CNVs only represent a very small fraction of the total amount of ultra-rare CNVs identified in the neurodevelopmental disorder clinic as well as in the general population.

Intelligence is a major trait assessed in the developmental pediatric and psychiatric clinic. There is a significant genetic correlation between intelligence and psychiatric disorders and cognitive impairments represent a major referral criterion to the NDD clinic. The heritability of general intelligence is estimated at around 50 to 80% *[13]*. The heritability of variants in linkage disequilibrium with common SNPs is estimated to be around 22.7%, with variants in poor linkage disequilibrium with SNPs, including rare CNVs, explaining 31.3% of the phenotypic variation in intelligence*[14]*. Two recent GWAS, have identified over 200 loci associated with intelligence and education*[15, 16]*, potentially implicating 1000 genes. The latter were largely non-overlapping with genes previously linked to ID*[15]*. Contrary to SNPs, there is no ambiguity in the molecular interpretation of a fully deleted or duplicated gene, which invariably decreases or increases transcription respectively. Therefore, CNVs represent a powerful tool to map the effect-sizes of genes (altered by gene dosage) on human traits.

We have previously proposed a framework to estimate and predict the effect-size on intelligence of CNVs. We showed that linear models*[17]* using the sum of the “probability of being loss-of-function intolerant” (pLI) scores*[18]* of all genes included in a deletion can predict their effect-size on intelligence quotient (IQ) with 75% accuracy. Our initial study was underpowered to measure the effect-size of duplications. It is also unknown if only a limited number of intolerant genes or a large proportion of genes within CNVs are driving effects on cognitive abilities. More broadly, the number of genes modulating general intelligence remains unknown. The pLI used in our earlier model, ranges from 0 to 1 but has a bimodal distribution and is essentially a categorical variable classifying genes as intolerant (>0.9) or tolerant (≤0.9) to protein-loss-of-function (pLoF) *[18]*. Continuous measures such as the LOEUF*[19]* (Loss-of-function Observed/Expected Upper bound Fraction) were recently introduced to reflect the full spectrum of intolerance to pLoF. LOEUF range from 0 to 2, and values below 0.35 are suggestive of intolerance.

Our present aims were 1) to test the robustness of effect-size estimates for CNVs across unselected and NDD populations, 2) to establish the effect-size on general intelligence of genomic duplications, 3) to investigate the quantitative relationship between effect-size on general intelligence and the frequency of *de novo* events, and 4) to estimate individual effect-sizes for all protein-coding genes that are intolerant as well as tolerant to pLoF.

We identified CNVs in 24,092 individuals from five general populations, two autism cohorts and one neurodevelopmental cohort. Measures of intolerance to pLoF were used as variables to estimate the effect of CNVs and individual genes on general intelligence. Validation procedures using cognitive data on CNVs from 47 published reports and the UKBB demonstrated a near 80% accuracy of model estimated. We implemented an online tool to help clinicians and researchers estimate the effect-size of any CNVs on general intelligence.

## Results

### 1) Deletions and duplications have a 3:1 effect-size ratio on general intelligence

We first sought to replicate our previous estimates for the effect-size of deletions on general intelligence computed using pLI *[17]*. We performed a meta-analysis on 20,151 individuals from 5 unselected populations (Table 1, Supplementary Fig. 1) showing that the deletion of one point of pLI decreases NVIQ or g-factor by 0.18 z-score (95% CI: −0.23 to −0.14, equivalent to 2.7 points of NVIQ, Fig. 1a, Supplementary Table 1). For duplications, we performed a meta-analysis using the same unselected populations. It shows that duplicating one point of pLI decreases NVIQ or g-factor by 0.04 z-score (95% CI: −0.09 to −0.01), which is equivalent to 0.75 points of IQ. Of notes, our previous study *[17]* was unable to estimate effect-sizes of duplications on general intelligence, likely due to sample size. There was no heterogeneity across cohorts. Sensitivity analyses showed that methods used for cognitive assessments did not influence these results (Fig. 1, Supplementary Table 2).

**Table 1.** Cohort descriptions. Cohorts include 24,092 individuals, including 14,874 unrelated individuals. SSC and CaG cohorts were broken down into sub-samples based on array technology (Supplementary methods). †63 and ‡ 12 cognitive measures were respectively used to compute the g-factor in SYS children and parents (Supplementary methods). NDD: neurodevelopmental disorders, SYS: Saguenay Youth Study, CaG: CARTaGEN, LBC: Lothian Birth Cohort, SSC: Simons Simplex Collection; n=number of individuals remaining for analysis after quality control. The mean and Standard Deviation (SD) for g-factor slightly deviate from 0 and 1 in some cohorts since they were computed on all available data (before the exclusion of some individuals for poor quality array) and summarized here only for individuals included in the analyses. *All individuals from LBC1936 were assessed at 70 years old explaining the absence of SD computation. **The NDD cohort was used only in the replication analysis and was not included in meta- or mega-analyses. *** We displayed the Z-scores of IQ, because IQ was preferred to g-factor for all analyses, even if results were similar (Supplementary Table 1 and 3).

| Ascertainment | Cohort | Array type | n= | Females, n (%) | Age in years Mean (SD) | Type of intelligence measures | Z-scored intelligence measure Mean (SD) |
| --- | --- | --- | --- | --- | --- | --- | --- |
| Unselected (n=20,151) | IMAGEN [36] | 610Kq; 660Wq | 1,744 | 891 (51%) | 14.4 (0.4) | WISC-IV (and g-factor, similarities score, vocabulary score, block design score, matrix reasoning score) | 0.44 (0.98) *** |
|  | SYS children[37] | 610Kq; HOE-12V | 967 | 505 (52%) | 15.0 (1.8) | WISC-III (and g-factor using 63 cognitive measures†) | 0.30 (0.87) *** |
|  | SYS parents[37] | HOE-12V | 602 | 321 (53%) | 49.5 (4.9) | g-factor, 12 cognitive measures‡ | 0 (1) |
|  | LBC1936[38] | 610Kq | 504 | 247 (49%) | 70.0 (-)* | Moray House Test (and g-factor) | 0.05 (0.96) *** |
|  | CaG-GSA[39] | GSA | 2,074 | 1,094 (53%) | 54.3 (7.6) | g-factor, Reasoning, Memory, Reaction time | -0.02 (1.03) |
|  | CaG-Omni2.5[39] | Omni2.5 | 515 | 281 (55%) | 52.4 (8.6) |  | -0.10 (1.02) |
|  | CaG (all)[39] | GSA; Omni2.5 | 2,589 | 1,375 (53%) | 53.9 (7.8) |  | -0.03 (1.03) |
|  | G-Scot[40] | 610Kq | 13,745 | 8,101 (59%) | 46.7 (15.0) | g-factor, Logical Memory, Digit Symbol, Verbal fluency, Mill Hill Vocabulary | 0.00 (0.99) |
| Autism (n=3,941) | SSC-1Mv1[41] | 1Mv1 | 332 | 44 (13%) | 9.5 (3.2) | WISC-IV n=19; DAS-II E-Y n=96; DAS-II S-A n=179; Mullen n=12; WASI-I n=26 | -0.55 (1.59) |
|  | SSC-1Mv3[41] | 1Mv3 | 1,182 | 157 (13%) | 8.8 (3.5) | WISC-IV n=16; DAS-II E-Y n=531; DAS-II S-A n=539; Mullen n=77; WASI-I n=19 | -0.98 (1.66) |
|  | SSC-Omni2.5[41] | Omni2.5 | 1,048 | 140 (13%) | 9.2 (3.7) | WISC-IV n=10; DAS-II E-Y n=403; DAS-II S-A n=494; Mullen n=124; WASI-I n=17 | -1.25 (1.87) |
|  | SSC (all)[41] | 1Mv1; 1Mv3; Omni2.5 | 2,562 | 341 (13%) | 9.03 (3.6) | WISC-IV n=45; DAS-II E-Y n=1,030; DAS-II S-A n=1,212; Mullen n=213; WASI-I n=62 | -1.03 (1.75) |
|  | MSSNG[30] | WGS | 1,379 | 275 (20%) | 9.2 (4.4) | WISC-IV n=46; WASI-II n=338; Leiter n=372; Raven n=214; Stanford Binet n=281; WPPSI n=128 | -0.47 (1.58) |
| NDD** (n=551) | Ste-Justine-probands | Agilent 180 K array | 132 | 52 (39%) | 7.23 (5.46) | WISC-V n=36; WASI-II n=8; WPPSI-IV n=38; Leiter-R n=18; Mullen n=32 | -1.31 (1.02) |
|  | Ste-Justine-siblings |  | 87 | 44 (50%) | 7.75 (5.72) | WISC-V n=28; WASI-II n=13; WPPSI-IV n=31; Leiter-R n=3; Mullen n=12 | -0.29 (0.98) |
|  | Ste-Justine-parents |  | 310 | 180 (58%) | 37.80 (7.13) | WASI-II | -0.10 (1.16) |
|  | Ste-Justine-other members |  | 22 | 12 (54%) | 43 (21.27) | WASI-II | -0.04 (1.32) |

**Fig. 1.**
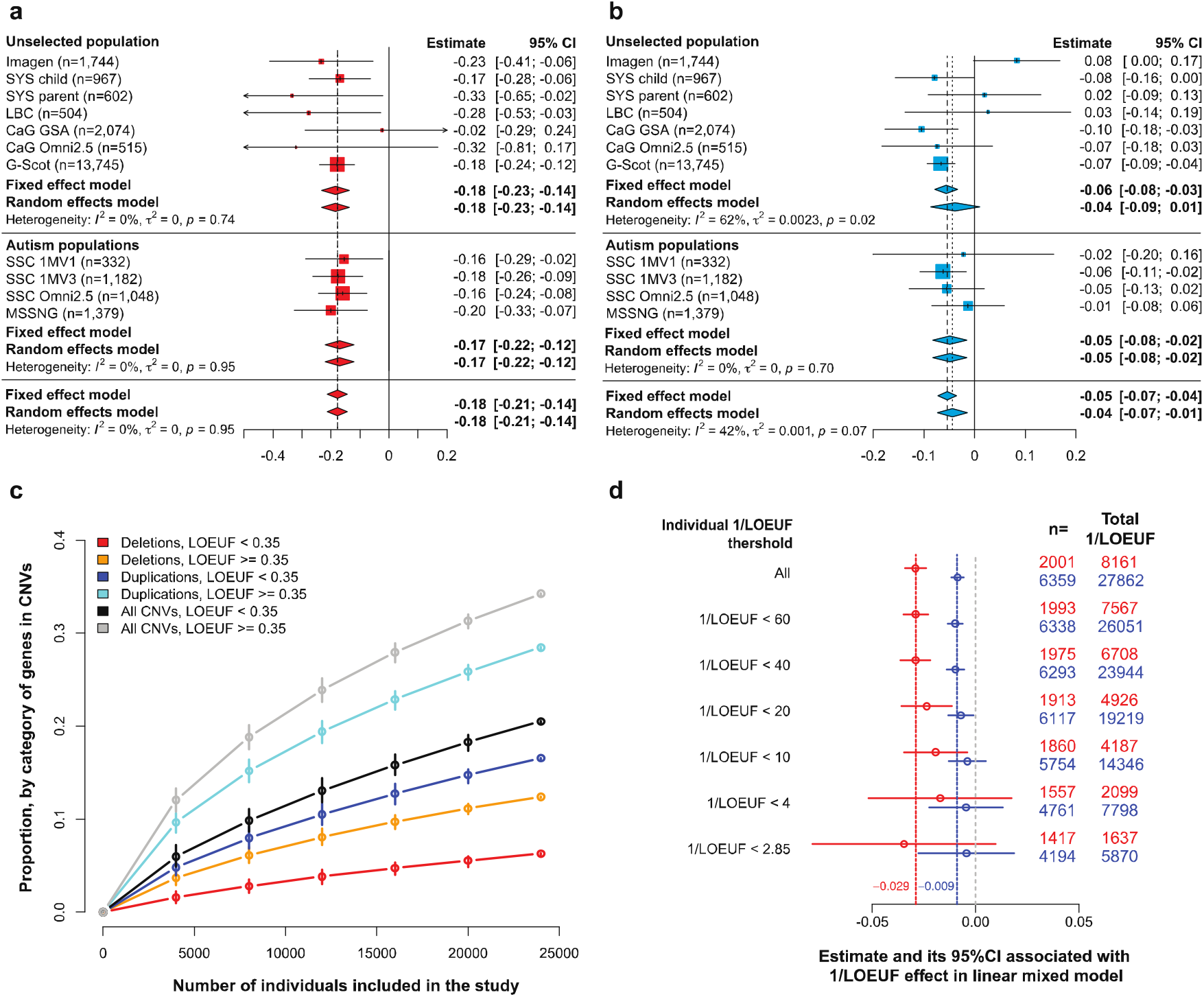
Effect of intolerant score on general intelligence measured for deletions and duplications. Meta-analysis estimating the effect of deletions **a.** and duplications **b.,** measured by sum of pLI, on general intelligence (Table S1). X-axis values represent z-scores of general intelligence. Deleting one point of pLI decreases the general intelligence by 0.18 z-scores (2.7 points of IQ). Duplicating one point of pLI decreases the general intelligence by 0.05 z-scores (0.75 points of IQ). The squares represent the effect-size computed for each sample. Their size negatively correlated to variance. Diamonds represent the summary effect across cohorts. Their lengths correspond to the 95% confidence intervals of the mean effect-size. **c.** Estimated proportion of the coding genome within each category defined by LOEUF, encompassed in CNVs present in the mega-analysis according to sample size (randomly selected within the mega-analysis). We observed N_CNVs gene_=6,315 with N_Del. gene_=2,282 and N_Dup. gene_=5,223). **d.** Estimated effect of 1/LOEUF on general intelligence after removing individuals with a sum of 1/LOEUF larger than 60, 40, 20, 10, 4 and 2.85 (2.85 corresponds to 1/0.35, the cut-off for intolerance to pLoF gnomAD). n: number of individuals with a total sum of 1/LOEUF > 0.

### 2) The effect-size of CNVs on general intelligence is not influenced by ascertainment

Since genomic variants with large effects on general intelligence are thought to be removed from the general population as a result of negative selective pressure, this may have led to an underestimation of the effect-size of CNVs in unselected populations. To examine this possibility, we analyzed 3,941 individuals (Table 1, Supplementary Fig. 1) from two autism cohorts, which include individuals with ID and *de novo* CNVs. Effect-sizes of pLI on general intelligence were the same than those observed in unselected populations for deletions and duplications and we did not observe any heterogeneity across cohorts (Fig. 1, Supplementary Table 1). Finally, we asked if effect-sizes of pLI were the same in large CNVs rarely observed in the general population or in autism cohorts. We tested 226 CNV carriers and 325 intrafamilial controls from 132 families ascertained in the clinic (Table 1). Effect-sizes of pLI on IQ were very similar with a decrease of 0.147 z-score, 95% CI: −0.18 to −0.11 (*P*= 1.1×10^−15^) in deletions and 0.069 z-score, 95% CI: −0.1 to −0.04 (*P*=8.7×10^−6^) in duplications (Supplementary Table 3).

### 3) Mega-analysis suggests additive effects of constraint scores on general intelligence

We pooled samples after adjusting for variables including cognitive test and cohorts to perform a mega-analysis of 24,092 individuals carrying 13,001 deletions and 15,856 duplications encompassing 36% of the coding genome (Fig. 1b, Supplementary Fig. 2a). The effect-size of pLI was unchanged, decreasing general intelligence by 0.175 z-score (SE=0.016, *P*=1.25×10^−28^) and 0.054 z-score (SE=0.009, *P*=1.90×10^−9^) for deletions and duplications, respectively (Supplementary Table 4). The partial R^2^ shows that deletions and duplications measured by pLI explain respectively 0.5% and 0.1% of the total variance of intelligence in the complete dataset; in line with the fact that large effect-size CNVs are rare in the general population.

Among 11 variables, the 2 main constraint scores (pLI and 1/LOEUF) best explained (based on AIC) the variance of general intelligence (Supplementary Table 4). For the remainder of the study, we transitioned to using LOEUF because it is a continuous variable (the pLI is essentially binary) and is now recommended as the primary constraint score by gnomAD. Analyses using pLI are presented in supplemental results.

There was no interaction between constraint scores and age or sex (Supplementary Table 5 to 8). Non-linear models did not improve model fit (Supplementary Table 9 to 10), suggesting an additive effect of constraint scores.

### 4) The effect-size of 1/LOEUF on intelligence is the same in recurrent neuropsychiatric CNVs and non-recurrent CNVs

We show that removing 608 individuals carrying any of the 121 recurrent CNV previously associated with neuropsychiatric conditions*[17]* does not influence the effect-size of 1/LOEUF on general intelligence (Supplementary Table 11). It has been posited that the deleteriousness of large psychiatric CNVs may be due to interactions between genes encompassed in CNVs. We therefore asked if the effect-size of 1/LOEUF is the same for CNVs encompassing small and large numbers of genes. We recomputed the linear model 6 times after incrementally excluding individuals with a total sum of 1/LOEUF ≥60, 40, 20, 10, 4 and 2.85 for deletions and duplications separately. Effect-sizes remain similar whether deletions encompass >10 or >60 points of 1/LOEUF (Fig. 1d, Supplementary Fig. 2b).

### 5) Gene dosage of 1% of coding genes shows extreme effect-size on general intelligence

Our ability to estimate large effect sizes is likely hampered by the explanatory variable (1/LOEUF) used in the model because there is only a 60-fold difference between the smallest and largest value. To improve model accuracy for large effect-size genes, we used a list of 256 ID-genes*[2, 20]*, previously identified with an excess of *de novo* mutations in NDD cohorts. We identified 126 CNVs encompassing at least one ID-gene (Fig. 2).

**Fig. 2.**
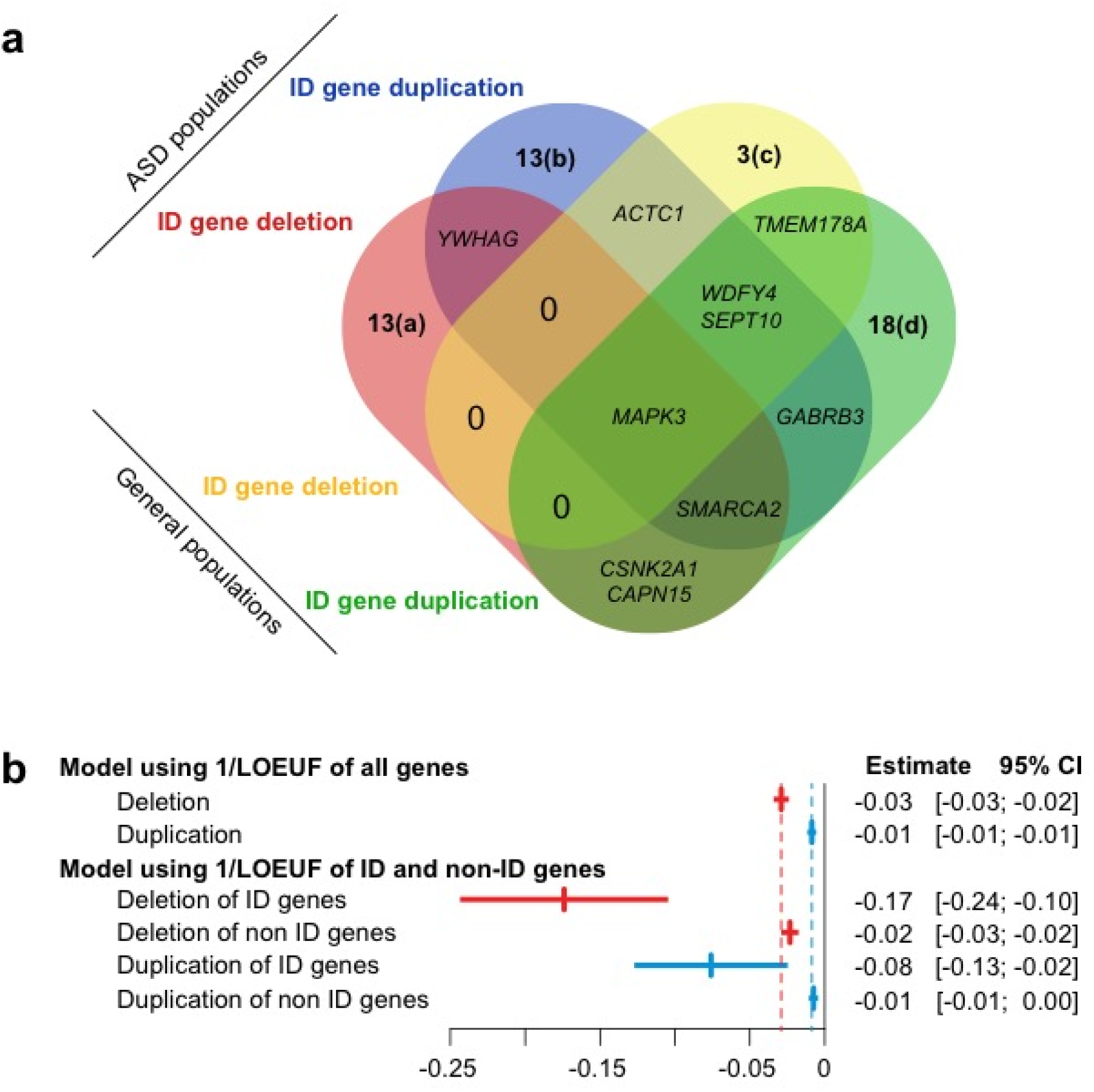
Effect-size of intellectual disability (ID) genes on general intelligence. **a.** Venn diagram of ID genes in ASD and in general population cohorts. We identified 66 CNVs encompassing at least one ID-gene in ASD cohorts (31 deletions and 35 duplications) and 60 in the general population (13 deletions and 47 duplications) (Supplementary methods). Genes were previously defined as harboring an excess of *de novo* loss of function (bold) or missense mutations in neurodevelopmental cohorts: (a) *DYNC1H1, **PHF21A, SHANK3, TRA2B, FOXP1, SETD5,** NR4A2, **TCF7L2, SOX5, POU3F3, ARID1B, EBF3, HNRNPU***; (b) ***SET, ZBTB18, DLG4, CHAMP1, CNOT3**, U2AF2, HIST1H2AC, DNM1, **RAI1, CREBBP, HIST1H1E, ASXL1**, CABP7;* (c) *PRPF18, PPP2R1A, EEF1A2; (d) TRAF7, DEAF1, STC1, **MYT1L, BRPF1**, CBL, **SPAST, WDR87, NFE2L3, STARD9, TCF20, KMT2C, FAM200B, KDM5B, CHD2**, BTF3, ITPR1, HMGXB3*. **b.** Effect-size of 1/LOEUF on general intelligence estimated in a model using two explanatory variables (sum of 1/LOEUF of deleted and duplicated genes) or 4 explanatory variables (sum of 1/LOEUF of ID genes and non-ID genes for deletions and duplication).

We recomputed the model by integrating 4 explanatory variables: the sum of 1/LOEUF for ID and non-ID-genes encompassed in deletions and duplications. The effect-size on intelligence of 1/LOEUF for ID-genes was 7 to 11-fold higher than the effect-size of non-ID genes which remained unchanged (Supplementary Table 12, 13 and Fig. 3). The mean effect of ID-genes intolerant to pLoF (LOEUF<0.35) was a decrease of 20 points of IQ for deletions and 9 points for duplications (Supplementary Table 13).

**Fig. 3.**
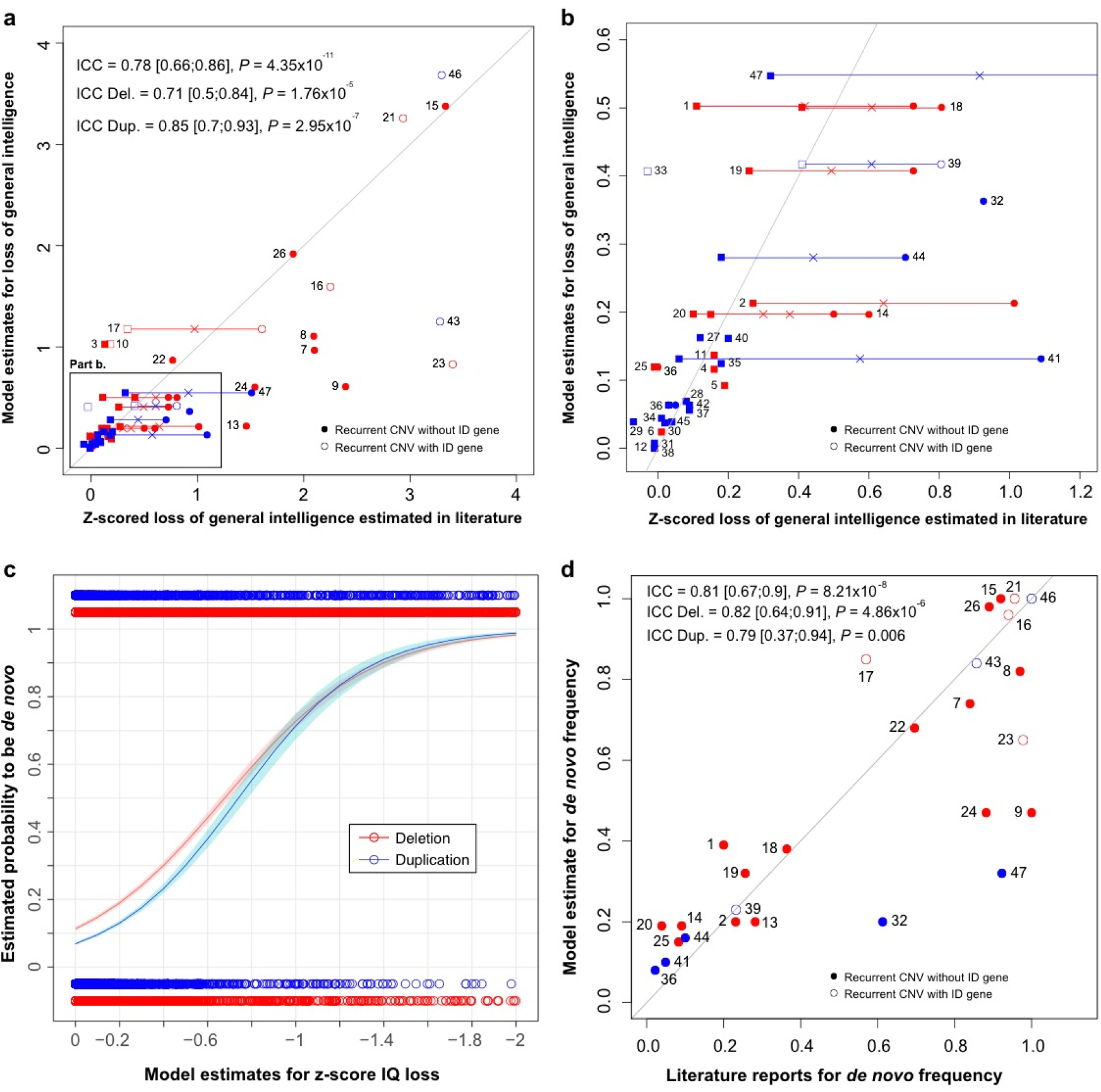
Concordance between model predictions and published observations for CNV effects on general intelligence and for de novo frequency. **a.** and **b.** Concordance between model estimates (with 1/LOEUF and ID-genes) and literature of clinical data and UKBB reports for general intelligence loss observed in respectively 27 and 33 recurrent CNVs for a total of ascertained carriers of 47 recurrent CNVs (Supplementary Table 15). X- and Y-values: effect size of CNVs on z-scored general intelligence. **b.** Zoom of the rectangle drawn in the lower left section of panel **a**. We represented values from clinical data by a circle and those from UKBB data by a square. The cross represents the mean value of z-scored IQ loss for the 13 recurrent CNVs observed both in literature and in UKBB. **c.** and **d.** The model uses 2 explanatory variables (1/LOEUF of non-ID-genes and ID-genes). **c.**Probability of *de novo* estimated by our *de novo* model (Y-axis) according to the loss of IQ estimated by a model using 1/LOEUF for ID and non-ID genes as two explanatory variables (X-axis). The *de novo* model was fitted on 13,114 deletions (red) and 13,323 duplications (blue) with available inheritance information observed in DECIPHER, CHU Sainte-Justine, SSC, MSSNG, SYS and G-Scot. **d.** Concordance between *de novo* frequency observed in DECIPHER (X-axis) and the probability of being *de novo* estimated by models when excluding recurrent CNVs of the training dataset (Y- axis) 1/LOEUF for ID and non-ID genes as an explanatory variable for 27 recurrent CNVs. The first bisector represents the perfect concordance. Deletions are in red and duplications in blue. Empty circles or square are CNVs encompassing ID-genes. ICC indicates intraclass correlation coefficient (3, 1). Each point represents a recurrent CNV: (1) TAR Deletion; (2) 1q21.1 Deletion; (3) 2q11.2 Deletion; (4) 2q13 Deletion; (5) *NRXN1* Deletion; (6) 2q13 (*NPHP1*) Deletion; (7) 3q29 (*DLG1*) Deletion; (8) 7q11.23 (William-Beuren) Deletion; (9) 8p23.1 Deletion; (10) 10q11.21q11.23 Deletion; (11) 13q12.12 Deletion; (12) 13q12 (*CRYL1*) Deletion; (13) 15q13.3 (BP4-BP5) Deletion; (14) 15q11.2 Deletion; (15) 16p11.2-p12.2 Deletion; (16) 16p13.3 ATR-16 syndrome Deletion; (17) 16p11.2 Deletion; (18) 16p11.2 distal Deletion; (19) 16p13.11 Deletion; (20) 16p12.1 Deletion; (21) 17p11.2 (Smith-Magenis) Deletion; (22) 17q12 Deletion; (23) 17q21.31 Deletion; (24) NF1-microdeletion syndrome Deletion; (25) 17p12 (*HNPP*) Deletion; (26) 22q11.2 Deletion; (27) TAR Duplication; (28) 1q21.1 Duplication; (29) 2q21.1 Duplication; (30) 2q13 Duplication; (31) 2q13 (*NPHP1*) Duplication; (32) 7q11.23 Duplication; (33) 10q11.21q11.23 Duplication; (34) 13q12.12 Duplication; (35) 15q11q13 (BP3-BP4) Duplication; (36) 15q11.2 Duplication; (37) 15q13.3 Duplication; (38) 15q13.3 (*CHRNA7*) Duplication; (39) 16p11.2 Duplication; (40) 16p11.2 distal Duplication; (41) 16p13.11 Duplication; (42) 16p12.1 Duplication; (43) 17p11.2 Duplication; (44) 17q12 (*HNF1B*) Duplication; (45) 17p12 (*CMT1A*) Duplication; (46) Trisomic 21 Duplication; (47) 22q11.2 Duplication.

### 6) Model explains nearly 80% of the effect-size of CNVs

As a validation procedure, we compared model estimates to published observations for 47 recurrent CNVs reported in clinical series and in the UKBB^17^ (Supplementary Table 14 and 15). When cognitive data was available from both clinical and the UKBB (n=13), we used the mean of both effect-sizes. Concordance between model estimates and previously published measures was 0.78 for all CNVs (95% CI, 0.66-0.86, *P*= 4.3×10^−11^, Fig. 4). Accuracy was similar for deletions (ICC=0.71 [0.5;0.84], *P*= 1.8×10^−5^) and duplications (ICC=0.85 [0.7;0.93], *P*= 3×10^−7^) as well as for small and large CNVs including trisomy 21 (Fig. 3a and 3b, Supplementary Fig. 5).

**Fig. 4.**
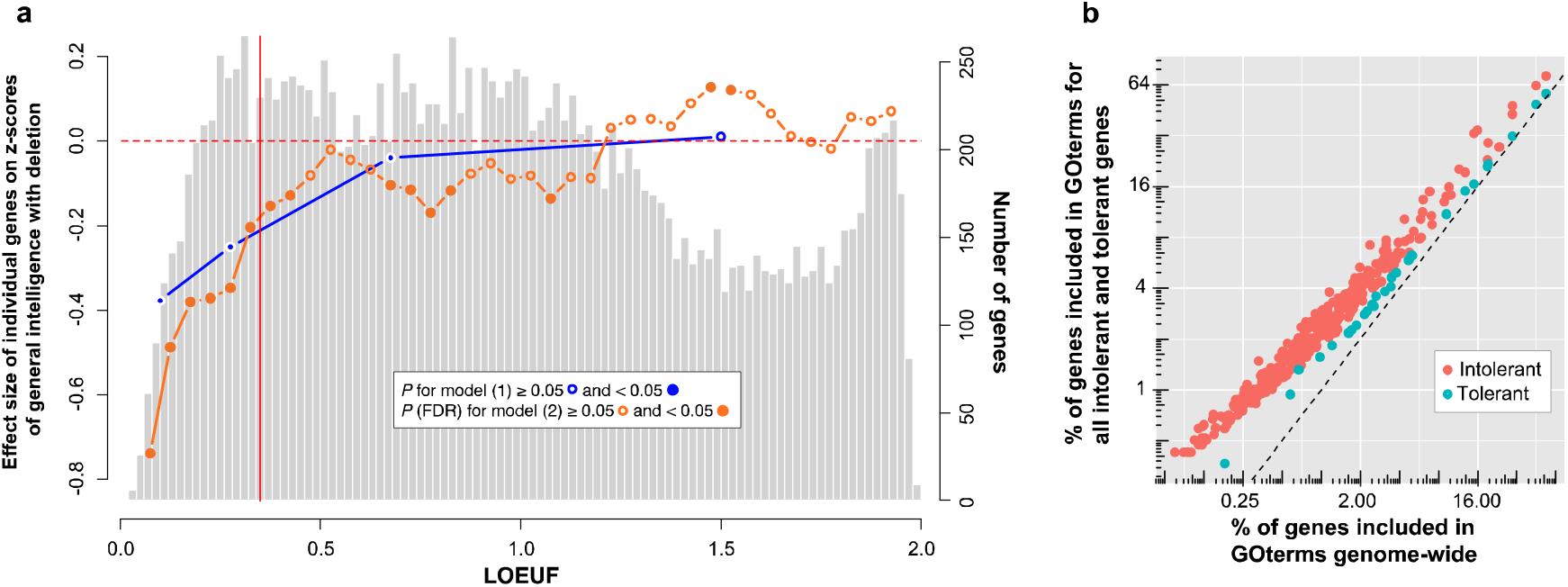
Effect-size on general intelligence of individual genes encompassed in CNVs and their GOterms enrichment. The light grey histogram represents the distribution of LOEUF values for 18,451 autosomal genes. The blue line represents the estimates for a gene in each of the 4 categories of LOEUF included in the model (Supplementary methods): highly intolerant genes (LOEUF <0.2, n=980), moderately intolerant genes (0.2≤LOEUF<0.35 n=1,762), tolerant genes (0.35≤LOEUF<1, n=7,442) and genes highly tolerant to pLoF (LOEUF≥1, n=8,267). The orange line represents the estimated effect-size of 37 categories of genes based on their LOEUF values (sliding windows=0.15) in the model (Supplementary methods). Genes with a LOEUF below 0.35 (vertical red line) are considered to be intolerant to pLoF by gnomAD. Left Y-axis values: z-scored general intelligence (1 z-score is equivalent to 15 points of IQ) for deletion. Right Y-axis values: number of genes represented in the histogram. For **Fig. 6b** each point represents a GOterm for which enrichment was observed for all intolerant (n=2,742) or tolerant genes (n=7,442) (**b.**), for all intolerant (n=609) or tolerant genes (n=2,251) encompassed in CNVs (C) when compared to the whole coding genome (Bonferroni). **b.** X-axis: % of genes included in the GOterm genome-wide; Y-axis: % of genes included in the GOterm for all intolerant (0<LOEUF<0.35) and tolerant genes (0.35≤LOEUF<1).

### 7) CNVs with the same impact on intelligence have the same*de novo* frequency

Because measures of intolerance to haploinsufficiency explain equally well the effect-sizes of deletions and duplications on intelligence, we investigated the relationship between effects on intelligence and *de novo* frequency for deletions and duplications. We established inheritance for 26,437 CNVs in 6 cohorts (Supplementary Table 16). There was a strong relationship between effects on general intelligence estimated by the model and the frequency of *de novo* observations for deletions (*P*=1.9×10^−65^) and duplications (*P*=4.6×10^−24^, Fig. 3c).

Deletions and duplications with the same impact on general intelligence show similar *de novo* frequency CNVs (Fig. 3c).

The concordance between the probability of occurring *de novo* estimated by the model (after removing recurrent CNVs) and *de novo* frequency reported in the DECIPHER database on 31 recurrent CNVs was 0.81 ([0.67-0.9]; *P*=8.2×10^−8^) (Fig. 3d, Supplementary Table 17 and Fig. 6).

### 8) Estimating effect-sizes of individual genes using LOEUF

Since we were underpowered to perform a gene-based GWAS, we first divided all genes in 4 categories: highly intolerant genes (LOEUF<0.2; n=980), moderately intolerant genes (0.2≤LOEUF<0.35 n=1,762), tolerant genes (0.35≤LOEUF<1; n=7,442) and highly tolerant genes (LOEUF≥1; n=8,267). This dichotomization of LOEUF values also allowed to test whether the previous linear models were driven by subgroups of genes. The sum of genes in each category was used as four explanatory variables to explain general intelligence in the same linear model. For deletions, highly, moderately intolerant and tolerant genes showed negative effects on general intelligence (Fig. 4a, Supplementary Table 18). For duplications only moderately intolerant genes showed negative effects (Supplementary Fig. 7 and Table 18). We were underpowered to further subdivide these LOEUF categories, so we tested 38 overlapping LOEUF categories in 38 linear models. Each model used 2 explanatory variables: number of genes within and outside the LOEUF category (size = 0.15 LOEUF). For haploinsufficiency, negative effects on general intelligence were observed for genes within 13 categories across intolerant and tolerant LOEUF values. For duplications, only 2 categories had negative effects (Fig. 4a, Supplementary Fig.7 and Table 19).

### 9) Most biological functions affect cognition

The 6,114 different genes encompassed in the CNVs of our dataset did not show any GOterm enrichment except for olfactory related terms (Supplementary Tables 20). We asked if intolerant (LOEUF<0.35) and tolerant genes (0.35<LOEUF<1), which negatively affect IQ in the analysis above were enriched in GOterms. All intolerant and tolerant genes genome-wide, were enriched in 365 and 30 GOterms respectively (Fig. 4b, Supplementary Tables 21, 22). The largest group of GOterms enriched in intolerant genes represented gene regulation (RNA polymerase II transcription factor activity, chromatin organization; Fig. S11), cell death regulation and neuronal function (dendrite and synapse). Among 23 tissues overrepresented in intolerant genes, adult brain and epithelium showed the strongest enrichment (Supplementary Table 21). Top enriched pathways included those in cancer, focal adhesion, Wnt signaling and MAPK (Supplementary Table 21). For tolerant genes, milder enrichments included translation (tRNA) and cytoskeletal structure. Among the 7 significant tissues adult brain showed the strongest enrichment (Fig. 4b, Supplementary Table 22 and Fig. 12). The 2,862 intolerant and tolerant genes encompassed in the CNVs of our dataset showed the same GOterm distribution observed above for the full intolerant and tolerant coding genome. Genes encompassed in CNVs were therefore represented well all molecular functions observed for each LOEUF group at the genome-wide level (Supplementary Table 23).

## DISCUSSION

Deletions and duplications have effect-sizes on cognitive ability that are robust across cohorts, clinical diagnoses, and general intelligence assessments. The effect-size ratio on cognitive ability of deletions to duplications is 3:1. The linear sum of pLI or 1/LOEUF predicted the effect-size on intelligence of deletions and duplications with equal accuracy (78%). Using categories of LOEUF values, we provide the first estimates for the individual effect-sizes of protein-coding genes, suggesting that half of the coding genome affects intelligence. The 2,862 genes encompassed in CNVs of our dataset show the same GOterm distribution observed in the intolerant and tolerant coding genome.

### Model validation and ascertainment biases

Models show 78% concordance with effect-size of CNVs on IQ from previous literature reports. Estimates are discordant for several CNVs, which may be due to either 1) unidentified large effect-size genes with unreliable LOEUF measures due to the small size of the protein coding region, and 2) ascertainment bias. However, biases from clinically referred individuals can be adjusted for using intrafamilial controls *[21, 22]*. This is confirmed by effect-sizes using the Ste Justine family genetic cohort. Also, our results suggest that the effect-size of pathogenic CNVs are underestimated in the UKBB*[21]* while those of small CNVs are largely overestimated in clinical series. The maximum effect size measured in UKBB was only 0.4 z-score including pathogenic CNVs such as 16p11.2, 2q11.2 deletions and 10q11.21-q11.23 deletion containing an ID-gene (*WDFY4*). On the other hand, the effect size of variants such as the 16p13.11 duplications and 1q21.1 CNVs are likely overestimated in clinical series*[23]*. Therefore, statistical models using a variety of disease and unselected cohorts are likely to provide the most accurate estimates. Surprisingly, an autism diagnosis is not associated with a different impact of CNVs on cognitive ability. A recent study characterizes this finding showing that CNVs similarly decrease IQ in autism and in unselected populations but are nevertheless more frequent in autism than in controls with same intelligence[24].

### Individual effect-sizes of genes, and go their GOterm enrichments

Our study is based on CNVs encompassing intolerant and tolerant genes with the same GOterm distribution observed in those LOEUF categories genome-wide. Only one percent of coding genes with the highest intolerance to pLoF has large effects on cognitive ability (20 and 9 IQ points for deletions and duplications of ID genes). The rest of the intolerant genes (15% of coding genes) have moderate to mild effect-sizes. The group of all intolerant genes is enriched in many GOterms including brain expression and gene regulation as previously reported for this group*[2, 25]*. Genes considered tolerant to pLoF (0.35<LOEUF<1; 40% of coding genes) impact intelligence with small effect-size and are only mildly enriched in GOterms. This is reminiscent of GWAS results for schizophrenia showing that most GOterms contribute to it’s heritability *[26]*.

### Potential clinical application

Models developed in this study provide a translation of gnomAD constraint scores into cognitive effect-sizes. Model outputs are implemented in a prediction tool (https://cnvprediction.urca.ca/), which is designed to estimate the population-average effect-size of any given CNV on general intelligence, not the cognitive ability of the individual who carries the CNV. If the cognitive deficits of an individual are concordant with the effect-size of the CNV they carry, one may conclude that the CNV contributes substantially to those deficits. When discordant (ie. The observed IQ drop is ≥15 points (1SD) larger than the model estimate), the clinician may conclude that a substantial proportion of the contribution lies in additional factors which should be investigated, such as additional genetic variants and perinatal adverse events (e.g. neonatal hypoxic ischemic injury, seizure disorders etc). If IQ cannot be reliably measured (ie. ≤ 4 years or in the case of severe behavioral disorders), the cognitive impact of the CNV predicted by the model may allow to anticipate the need for potential interventions. Overall, the output of this tool can help interpret CNVs in the clinic, but estimates should be interpreted with caution. The model can provide an estimate for the effect size on intelligence of individual genes when deleted. Therefore, one may use this information to estimate the effect size on intelligence of any SNV resulting in a loss of function. However, larger datasets are required to refine the estimates for individual gene.

### The relationship between genetic fitness and cognitive abilities

The reasons underlying the tight relationship between general intelligence and epidemiological measures of intolerance to pLoF, is unclear. This relationship is further highlighted by the fact that deletions and duplications with the similar impact on intelligence occur *de novo* with similar frequencies. Behavioral interpretations are intuitive for severe ID but do not apply for CNVs with much milder effects. In other words, individuals with moderate or severe ID have limited offspring due to behavioral deficits but it is unclear how small changes in intelligence may lead to behavioral issues resulting in decreased fitness. Our results also suggest that genes considered as “tolerant” with LOEUF <1 affect cognitive abilities and are likely under “mild constraint”. Larger samples are required to better characterize the effect of this broad category of “mildly intolerant” genes on cognitive ability.

### Limitations

The model relies on constraint scores (LOEUF or pLI), which are epidemiological measures of genetic fitness in human populations, without any consideration of gene function*[18, 19]*. It is likely that some genes decrease fitness (eg. genes involved in fertility) without affecting general intelligence. Further studies combining intolerance scores with functional categories are required to investigate this question. While LOEUF was designed to measure intolerance to loss of function, we used it to assess both deletions and duplications. However, our results and a recent report suggest that it also measures the intolerance to increased gene expression *[27]*. Noise in the model may be related to unreliable constraint scores computed for small genes with a limited number of pLoF variants observed in the gnomAD database. Bias in the model may be introduced by ID genes observed in our dataset. Indeed, they may reflect a less severe subgroup and model outputs should be interpreted with caution when CNVs encompass ID-genes. Another potential bias is related to the fact that models were trained on CNVs encompassing 36% of the coding genome. Projections suggest that 500K individuals from an unselected population would cover 78% (Fig. S8).

Finally, all models imply additive effects and massive datasets would be required to test for gene-gene and gene-environment interactions. However, the fact that very large CNVs (such as trisomy 21) are accurately estimated by the model suggests that genetic interactions within large genomic segments or even chromosomes cannot be readily observed. There is long standing discordance between observations made at the microscopic and macroscopic level. Indeed, molecular studies provide unequivocal evidence that gene-gene interactions are common but quantitative genetic theory suggests that contributions from non-additive effects to phenotypic variation in the population are small. Reconciling these two observations, polygenic models assume that interactions are the rule rather than the exception. Interactions are, in fact, accounted for in the additive models[28]. For example, LOEUF values are correlated with the number of protein-protein interactions[19] and our results also show that the intolerant genes are enriched in GOterms linked to “gene regulation”. In other words, the level of interactions for a given gene is directly related to its “individual” effect size on intelligence (ie. chromatin remodelers have a very broad interaction network, low LOEUF values and high effect sizes on intelligence).

## Conclusions

The effect-size of deletions or duplications on intelligence can be accurately estimated with additive models using constraint scores. The same relationship between gene dosage and cognition apply to small benign CNVs as well as extreme CNVs such as Down syndrome. We provide a map of effect-sizes at the individual gene level but to move beyond this rough outline, much larger sample sizes are required. Nonetheless, these results suggest that a large proportion (56%) of the coding genome covering all molecular functions influences cognitive abilities. One may therefore view the genetic contribution to cognitive difference as an emergent property of the entire genome not restricted to a limited number of biological pathways.

## Materials and Methods

### 1. Cohorts

We included five cohorts from the general population, two autism cohorts and one familial cohort with at least one CNV-carrier child recruited for a neurodevelopmental disorder (Table1). Studies for each cohort were reviewed by local institutional review boards. Parents/guardians and adult participants gave written informed consent and minors gave assent.

### 2. Measures of general intelligence

General intelligence was assessed using the neurocognitive tests detailed in table 1. Measures of non-verbal intelligence quotient (NVIQ) were available in five cohorts and general intelligence factor (g-factor)*[29]* was computed in four cohorts, based on cognitive tests, primarily assessing fluid non-verbal reasoning (Table1, Supplementary Fig. 1). Intelligence measures were normalized using z-score transformations to render them comparable. The concordance between z-scored NVIQ and g-factor available for three cohorts ranged from 60 to 77% (Supplementary Table 24).

### 3. Genetic information

#### CNV calling and filtering

For all SNP array data, we called CNVs with PennCNV and QuantiSNP using previously published methods *[17]*. For the MSSNG dataset*[30]*, we used CNVs called on whole genome sequencing by Trost *et al. [31]*.

CNV filtering steps were previously published (Supplemental material). For the mega-analysis, we applied an additional filtering criterion, selecting CNVs encompassing at least 10 probes for all array technologies used across all cohorts.

The Sainte-Justine CNV-family cohort included participants on the basis of one pathogenic CNV identified in the diagnostic cytogenetic laboratory using an Agilent 180K array.

#### Annotation of CNVs

We annotated the CNVs using Gencode V19 (hg19) with ENSEMBL (https://grch37.ensembl.org/index.html). Genes with all transcripts fully encompassed in CNVs were annotated using 12 variables present in previous article*[17]*. Non-coding regions were annotated with the number of expression quantitative trait loci (eQTLs) regulating genes expressed in the brain*[32]*. CNV scores were derived by summing all scores of genes within CNVs*.[17]*. Also, we used a list of 256 ID-genes*[2, 20]*, previously identified with an excess of *de-novo* mutations in NDD cohorts.

### 4. Statistical analyses

#### Modelling the effect of CNVs on intelligence

General intelligence was adjusted within each cohort for age and sex when required (*Z_adj Intell._*; see supplemental material and Supplementary Fig. 9 and 10). To estimate the effect of CNVs on general intelligence, we fit the model developed by Huguet at al. *[17]* where the sum of pLI (or any of the 10 other scores) for all genes encompassed in deletions or duplications, respectively, is the variable used to predict the adjusted Z-score of general intelligence:

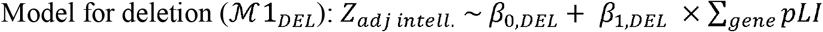

where, *β*_0,*DEL*_, *β*_1,*DEL*_ are the regression coefficients. The same model was applied to duplications. First, models 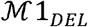 and 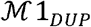 were fitted independently and adjusted for each cohort and results were used in the meta-analyses. Second, in the mega-analysis, 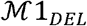 and 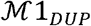 were fitted after pooling all samples and adjusting on the type of cognitive measure and cohort.

To take into account ID-genes that have a greater impact on intelligence, we used a model including 4 predictive variables 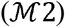:

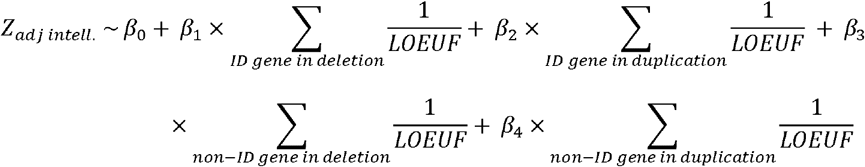

where *β*_0_, *β*_1_, *β*_2_, *β*_3_ and *β*_4_ are the regression coefficients.

The variance explained by deletions and duplications (measured by pLI) was computed using partial R^2^ in the full dataset as well as the subgroup (n=14,874) of unrelated individuals.

#### Sensitivity analyses

We tested non-linearity of the effect of haploinsufficiency scores on general intelligence by using polynomial regression model and by exploring a smooth function of the effect of haploinsufficiency scores using a Gaussian kernel regression method (https://cran.r-project.org/web/packages/KSPM/index.html) flexible enough to account for various types of effects (Supplementary material).

#### Model Validation

To validate our models, we computed the concordance between model predictions and loss of IQ measured for 47 recurrent CNVs obtained in previous publications (supplementary material). The concordance was computed using the intraclass coefficient correlation of type (3,1) (ICC_(3,1)_) *[33]*.

#### Modelling the probability to be de novo

We performed logistic regressions to estimate the probability of a CNV being *de novo* (*P_de novo_*) as a function of the haploinsufficiency scores:

Model for deletions 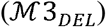:

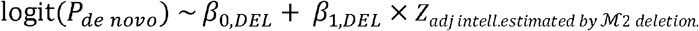

where *β*_0,*DEL*_, *β*_1,*DEL*_ are the regression coefficients. The same model was applied to duplications (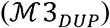)

For these analyses, we added two clinical populations (Decipher, decipher.sanger.ac.uk/) and the cytogenetic database of Sainte-Justine Hospital, where genetic data could be compared between the child and their parents, and applied the same filtering as for the previous CNV selection leading to a total of 26,437 CNVs. (Supplementary Table 16). The binary outcome variable was the type of transmission (1=*de novo*, 0=inherited).

To validate these models, we computed the concordance between model estimates and percentage of *de novo* variants computed with Decipher for 27 recurrent CNVs.

#### Estimating the effect-size of individual genes based on LOEUF values

We used 4 categories of LOEUF values to estimate the effect-size of genes classified as highly intolerant (LOEUF <0.2, n=980), moderately intolerant (0.2≤LOEUF<0.35 n=1,762), tolerant (0.35≤LOEUF<1, n=7,442), and highly tolerant to haploinsufficiency (LOEUF≥1, n=8,267). For deletions, model 4 is as follow:

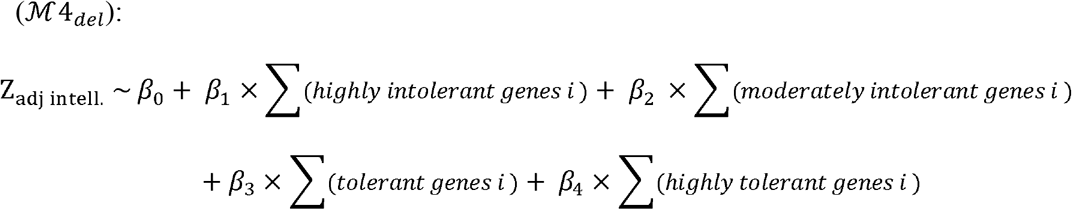

where *β*_0,*CVN type*_, *β*_1,*CVN type*_, *β*_2,*CVN type*_, *β*_3,*CVN type*_ and *β*_4,*CVN type*_ are the regression coefficients. The same model was applied for duplications.

To explore smaller categories of LOEUF values, we slid a window of size 0.15 LOEUF units, in increments of 0.05 units thereby creating 38 categories across the range of LOEUF values. We performed 38 linear models:

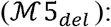

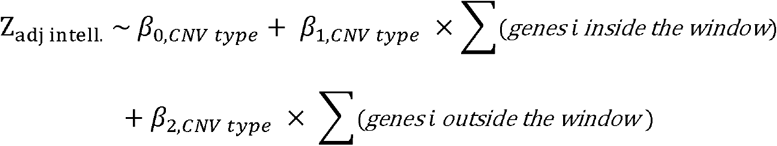

where *β*_0,*CVN type*_, *β*_1,*CVN type*_, and *β*_2,*CVN type*_ are the regression coefficients.

The same models were performed for duplications. Estimates were corrected for multiple testing (38 tests) using FDR.

#### GOterms Enrichment

For the GOterms enrichment for the tolerant and intolerant genes with all a genome and CNVs between unselected, ASD and both populations, we used DAVID release 6.8*[34]* (https://david-d.ncifcrf.gov). We kept the defaults parameters and save only the terms with Bonferroni corrected p-values <0.05. We then passed the list to REVIGO*[35]* (http://revigo.irb.hr/) to summarize and group the redundant GO.

## Conflict of interest

The authors declare that they have no conflict of interest.

## Funding/Support

This research was enabled by support provided by Calcul Quebec (http://www.calculquebec.ca) and Compute Canada (http://www.computecanada.ca). Sebastien Jacquemont is a recipient of a Bursary Professor fellowship of the Swiss National Science Foundation, a Canada Research Chair in neurodevelopmental disorders, and a chair from the Jeanne et Jean Louis Levesque Foundation. Catherine Schramm is supported by an Institute for Data Valorization (IVADO) fellowship. Petra Tamer is supported by a Canadian Institute of Health Research (CIHR) Scholarship Program. Guillaume Huguet is supported by the Sainte-Justine Foundation, the Merit Scholarship Program for foreign students, and the Network of Applied Genetic Medicine fellowships. Thomas Bourgeron is a recipient of a chair of the Bettencourt-Schueler foundation. This work is supported by a grant from the Brain Canada Multi-Investigator initiative and CIHR grant 159734 (Sebastien Jacquemont, Celia Greenwood, Tomas Paus). The Canadian Institutes of Health Research and the Heart and Stroke Foundation of Canada fund the Saguenay Youth Study (SYS). SYS was funded by the Canadian Institutes of Health Research (Tomas Paus, Zdenka Pausova) and the Heart and Stroke Foundation of Canada (Zdenka Pausova). Funding for the project was provided by the Wellcome Trust. This work was also supported by an NIH award U01 MH119690 granted to Laura Almasy, Sebastien Jacquemont and David Glahn and U01 MH119739. The authors wish to acknowledge the resources of MSSNG (www.mss.ng), Autism Speaks and The Centre for Applied Genomics at The Hospital for Sick Children, Toronto, Canada. We also thank the participating families for their time and contributions to this database, as well as the generosity of the donors who supported this program. We are grateful to all the families who participated in the Simons Variation in Individuals Project (VIP) and the Simons VIP Consortium (data from Simons VIP are available through SFARI Base). We thank the coordinators and staff at the Simons VIP and SCC sites. We are grateful to all of the families at the participating SSC sites and the principal investigators (A. Beaudet, M.D., R. Bernier, Ph.D., J. Constantino, M.D., E. Cook, M.D., E. Fombonne, M.D., D. Geschwind, M.D., Ph.D., R. Goin-Kochel, Ph.D., E. Hanson, Ph.D., D. Grice, M.D., A. Klin, Ph.D., D. Ledbetter, Ph.D., C. Lord, Ph.D., C. Martin, Ph.D., D. Martin, M.D., Ph.D., R. Maxim, M.D., J. Miles, M.D., Ph.D., O. Ousley, Ph.D., K. Pelphrey, Ph.D., B. Peterson, M.D., J. Piggot, M.D., C. Saulnier, Ph.D., M. State, M.D., Ph.D., W. Stone, Ph.D., J. Sutcliffe, Ph.D., C. Walsh, M.D., Ph.D., Z. Warren, Ph.D., and E. Wijsman, Ph.D.). We appreciate obtaining access to phenotypic data on SFARI base.

## Additional Contributions

Julien Buratti (Institute Pasteur), and Vincent Frouin, Ph.D. (Neurospin), acquired data for IMAGEN. Manon Bernard, BSc (database architect, The Hospital for Sick Children), and Helene Simard, MA, and her team of research assistants (Cégep de Jonquière) acquired data for the Saguenay Youth Study. Antoine Main, M.Sc. (UHC Sainte-Justine Research Center, HEC Montreal), Lionel Lemogo, M.Sc. (UHC Sainte-Justine Research Center), and Claudine Passo, Pg.D. (UHC Sainte-Justine Research Center), provided bioinformatical support. Maude Auger, Pg.D.; and Kristian Agbogba, B.Sc. (UHC Sainte-Justine Research Center), provided website development. Dr. Paus is the Tanenbaum Chair in Population Neuroscience at the Rotman Research Institute, University of Toronto, and the Dr. John and Consuela Phelan Scholar at Child Mind Institute, New York.

## Role of the Funder/Sponsor

The funder had no role in the design and conduct of the study; collection, management, analysis, or interpretation of the data; preparation, review, or approval of the manuscript; or decision to submit the manuscript for publication.

## Notes

### Competing Interest Statement

The authors have declared no competing interest.

### Summary of Updates

We developed a new section, "Most biological functions affect cognition", based on GOterm analysis with an intolerant gene.

