## Supplementary material for "Genome wide analysis of gene dosage in 24,092 individuals shows that 10,000 genes modulate cognitive ability": Method_and_Supplemental_data

**Abbreviations:**

**Samples:**

- unselected cohorts/populations
- autism cohorts/populations
- G-Scot: generation Scotland
- IMAGEN
- CaG-Omni2.5; CaG-GSA: Cartagene (by technology)
- LBC1936: Lothian birth cohort
- SSC-Omni2.5; SSC-1Mv1; SSC-1Mv3: Simon simplex collection (by technology)
- SYS parents; SYS children: Saguenay youth study
- MSSNG
- Sainte-Justine family genetic cohort

**Statistics:**

- *n*: sample size
- *P*: p-value
- SE: standard error
- SD: standard deviation
- *ρ*: correlation

**Genetics/genetic scores:**

- pLI,
- pLI (gnomAD)
- O/E: observed over expected
- O/E-upper CI
- RVIS
- DEL score; DUP score
- PPI: protein-protein interaction
- DS
- PSD genes
- FMRP genes
- eQTL
- No. of genes

**Other variables:**

- “type of test-cohort” variable
- g-factor: general factor
- NVIQ: non-verbal intelligence quotient
- Age in years
- z-scored measure of general intelligence

**List of Extended Data Tables**

**Supplementary Table 1.** Estimates associated with pLI within each sample included in the meta-analyse.	21

**Supplementary Table 2.** Comparison of estimates associated with pLI when measuring the effect-size of CNVs on IQ and g-factor.	21

**Supplementary Table 3.** Estimates obtained for models performed in Ste-Justine neurodevelopmental cohort.	22

**Supplementary Table 4.** Comparison of model fit for general intelligence according to annotation score.	22

**Supplementary Table 5.** Linear regression models including pLI x age interaction as predictor of general intelligence.	23

**Supplementary Table 6.** Linear regression models including 1/LOEUF x age interaction as predictor of general intelligence.	23

**Supplementary Table 7.** Linear regression models including pLI x sex interaction as predictor of general intelligence.	24

**Supplementary Table 8.** Linear regression models including 1/LOEUF x sex interaction as predictor of general intelligence.	24

**Supplementary Table 9.** Linear regression models performed in the mega-analysis to measure the effect of deleted or duplicated units of pLI on general intelligence on general intelligence.	25

**Supplementary Table 10.** Linear regression models performed in the mega-analysis to measure the effect of deleted or duplicated units of 1/LOEUF on general intelligence.	26

**Supplementary Table 11.** Linear regression models performed in the mega-analysis to measure the effect of deleted or duplicated units of pLI or 1/LOEUF on general intelligence for 3 sensitivity analyses based on exclusion of individuals carrying recurrent CNV or CNV containing ID-gene.	27

**Supplementary Table 12.** Linear regression models performed in the mega-analysis to measure the effect of deleted or duplicated units of pLI or 1/LOEUF, by gene category (ID- and non ID-genes) on general intelligence.	27

**Supplementary Table 13.** Distribution of the effect associated with deletion or duplication of genes by gene categories.	28

**Supplementary Table 14.** Description of the effect of 47 recurrent CNVs on general intelligence and probability of being *de novo*, estimated using empirical data from the literature and/or UKBB and using our models.	30

**Supplementary Table 15.** Empirical data on recurrent CNVs.	31

**Supplementary Table 16.** Detailed table of content for the de novo analysis using Decipher, Sainte-Justine UHC, MSSNG, SSC, Imagen, Generation Scotland, and SYS cohorts.	32

**Supplementary Table 17.** Results of the probability of being de novo in function of pLI and 1/LOEUF.	33

**Supplementary Table 18.** Estimated effects of deleted or duplicated individual genes on general intelligence according to 4 categories of tolerance to pLOF defined by LOEUF.	34

**Supplementary Table 19.** Estimated effects of deleted of duplicated individual genes on general intelligence according to several moving categories defined by a sliding window on LOEUF.	36

**Supplementary Table 20.** DAVID enrichment details with all genes within CNV for each population.	37

**Supplementary Table 21.** DAVID enrichment details with intolerant genes in the genome.	37

**Supplementary Table 22.** DAVID enrichment details with tolerant genes in the genome.	37

**Supplementary Table 23.** DAVID enrichment details with intolerant and tolerant genes within CNV for each population.	37

**Supplementary Table 24.** Correlation and concordance between NVIQ and g-factor in 3 unselected populations.	37

**List of Extended Data Figures**

**Supplementary Fig. 1.** Distribution of z-scored general intelligence measured either by NVIQ or g-factor according to the age of individuals and colored by cohort.	38

**Supplementary Fig. 2.** Sensitivity analyses for model based on pLI score.	39

**Supplementary Fig. 3.** Effect size associated with pLI divided by category of intellectual disability (ID) genes on general intelligence.	39

**Supplementary Fig. 4.** Concordance between observation from literature and UKBB for CNV effects on general intelligence.	40

**Supplementary Fig. 5.** Concordance between model predictions and published observations for CNV effects on general intelligence.	41

**Supplementary Fig. 6.** Estimated probability of *de novo*, based on model including pLI for ID and non ID genes, and its concordance with *de novo* frequency observed in literature.	42

**Supplementary Fig. 7.** Estimated effects of individual genes on general intelligence according to categories based on LOEUF for duplications.	43

**Supplementary Fig. 8.** Prediction of gene coverage according to the sample size.	44

**Supplementary Fig. 9.** Distribution of z-scored measure of general intelligence before (left panels) and after (right panels) adjustment for age (g-factor in general populations).	45

**Supplementary Fig. 10.** Distribution of z-scored measure of general intelligence before (left panels) and after (right panels) adjustment for sex (NVIQ in autism populations).	46

**Supplementary Fig. 11.** TreeMap view of GO-term clusters for intolerant genes in the genome.	47

**Supplementary Fig. 12.** TreeMap view of GO-term clusters for tolerant genes in the genome.	48

### Supplementary Information

#### 1. Cohorts

We included five cohorts from the general population, two autism cohorts and one familial cohort with at least one CNV-carrier child recruited for neurodevelopmental disorder (Table1). All cohorts were reviewed by local institutional review boards. Parents/guardians and adult participants gave written informed consent and minors gave assent.

##### 1.1. General population cohorts

We included 1744 unrelated adolescents from Imagen*[1]* and 967 children from the Saguenay Youth Study (SYS)*[2]*, two cohorts with available intelligence quotient (IQ) measures, previously pooled and studied in Huguet et al 2018*[3]*. Here both cohorts of children were analyzed separately. We also included 602 parents from SYS for which a g-factor was derived. In this study, we added three other general population cohorts.

CartaGene (CaG)*[4]* is a cohort of 6,184 unrelated French Canadians. After quality control, we included in our analysis 2,589 individuals with an available cognitive evaluation to compute the g-factor. When data are not pooled, CaG cohort was divided into two samples: Cag-GSA (N=2,074) and CAG-Omni2.5 (N=515) according to genotyping technology.

The Generation Scotland (G-Scot)*[5]* is a cohort including 16,884 individuals from the UK (6,020 families). After quality control, we keep for our analysis 13,745 individuals from 5,622 families for which cognitive evaluation allowed us to compute the g-factor*[6]*.

The Lothian Birth Cohort*[7]* 1936 (LBC1936) includes 1,091 individuals with IQ measured longitudinally in 1947 and in 2004-2007. After quality control, we included in our analysis 504 individuals having an IQ measure obtained at ~70 years old.

Of notes, cognitive measurements other than IQ are available for Imagen, SYS-children and LBC1936 such as a g-factor could be derived to assess the correlation between IQ and g-factor in these three samples.

##### 1.2. Disease cohorts

Simon Simplex Collection (SSC)*[8]* cohort includes 2,590 simplex families with one child affected by autism (proband), its parents and for some, its unaffected sibling. Only probands, assessed with IQ test, were kept in our study leading to 2,562 individuals after quality control. We analyzed separately three SSC samples: SSC-1Mv1 (N=332), SSC-1Mv3 (N=1,182) and SSC-Omni2.5 (N=1,048) according to genotyping technology.

MSSNG*[9]* is an international consortium (N=3,441) with 2,129 simplex and 617 multiplex autism families including at least one to five affected children. After quality control, we included in our analysis 1379 probands with available IQ measure.

We also included the Sainte-Justine family genetic cohort. Cognitive and behavioral measures were collected in families who carry a CNVs classified as pathogenic or as a variant of unknown significance (VUS). CNV carriers and their first-degree relatives were ascertained from the pediatric developmental disorder clinic. Data includes 202 families with at least one child carrying a large or a recurrent CNV. All 551 participants (130 fathers, 180 mothers, 132 probands, 87 siblings, 21 grand-parents and one oncle) were assessed with IQ test. The characteristics of this cohort reflects the criteria for chromosomal microarray (CMA) testing in the developmental pediatric clinic which include: intellectual disabilities (ID), learning disabilities, autism spectrum disorder as well as children with several comorbidities including attention deficit hyperactivity disorder (ADHD), speech and language disorders and developmental coordination disorders (DCD). The same assessments were performed in first degree relatives (carriers and non-carriers) making it possible to adjust for the effect of additional genetic and environmental background.

#### 2. Definition of phenotypes: IQ and g-factor

In this study, we focus on the non-verbal IQ (NVIQ) when available and on the g-factor otherwise. Both intelligence measures were normalized using z-score transformations to render them comparable. The NVIQ z-score has a mean of 100 and a standard deviation (SD) of 15. Since cognitive measures used in the computation of the g-factor are not the same between cohorts, the g-factor was computed and normalized separately within each cohort using the mean and SD computed on all available individuals. This was feasible since the g-factor was computed in general population cohorts only. Of note, g-factor was computed before excluding individuals due to array quality control, leading to g-factors with means and SDs slightly different from 0 and 1 for the final subset of individuals included in our analyses.

##### 2.1. Evaluation of IQ

IQ is an age-standardized general cognitive ability metric that provides an estimate of how one ranks relative to age-matched peers. In all analyses, IQ here refers to NVIQ, which is also called performance IQ. In the general populations examined here (Table 1), IQ has been measured using the Wechsler Intelligence Scale for Children (Imagen: WISC-IV*[10]*; SYS: WISC-III*[11]*) and the Moray house test*[12]**[13]* (LBC1936). Of note, in LBC1936, the Moray house test is based on cognitive measures obtained at 70 years old and converted into an IQ-type scale with a mean of 100 and SD of 15, as it was previously done in Gow *et al*.*[14]*.

In disease cohorts, adapted tests have been used, such that IQ measures were obtained using Mullen scales of early learning*[15]*, the Leiter international performance scale – original and revised*[16, 17]*, the Raven progressive matrices*[18]*, Stanford-Binet intelligence scale*[19]*, WISC-IV*[20]*, WISC-V*[21]*, Wechsler Abbreviated Scale of Intelligence (WASI-I; WASI-II)[22–24], Wechsler Preschool and Primary Scale of Intelligence (WPPSI-IV) *[25]*, Leiter-reviewed and the Differential Ability Scales (DAS-II *[26]*) (Table 1).

In the Sainte-Justine family genetic cohort, five adults were removed due to a large difference between the performance IQ and the verbal IQ which invalidates their full-scale IQ. Also, 4 probands were removed due to either floor effect or ceiling effect in the tests.

##### 2.2. G-factor computation

The g-factor is an indirect measure of general intelligence, obtained by extracting the first unrotated principal component from principal component analysis (PCA) of different standardized cognitive measures. We computed a g-factor in samples for which IQ was not available (SYS-parents, CaG and G-Scot) as well as in samples for which IQ and cognitive measures allowing g-factor computation were available, only in order to compare both intelligence measures (Imagen, SYS children, LBC1936).

For the LBC1936 cohort, we used the g-factor*[27]* that was already calculated by the LBC consortium. It is based on 6 non-verbal subtests of the Wechsler Adult Intelligence Scale-III*[28]*: matrix reasoning, letter number sequencing, block design, symbol search, digit symbol, and digit span backward. We transformed this z-score t using the mean of 0.02 and the SD of 0.987 that was calculated for the available LBC subjects.

The cognitive measures available for parents and children in the SYS cohort are different, thus the g-factor was not based on the same measures for the two subgroups.

In SYS parents sample, the g-factor was computed using 12 cognitive performances^2^ assessed using the Cambridge brain sciences platform*[29]*: color-word remapping, spatial planning, self-ordered search, paired associates learning, digit span, spatial span, visuospatial working memory, interlocking polygons, feature match, odd one out, grammatical reasoning and spatial rotation. The obtained g-factor represents 31.6% of the observed variance. Then, the g-factor was transformed to a z-score using the mean of -6.221×10^-12^ and the SD of 1.948.

In SYS children, the g-factor was computed using 63 cognitive measures*[2]* : Dot Location (Visual/non-verbal memory), Newman’s Card Sorting Task (Perseveration), Self-ordered Pointing Task (Working memory), Grooved Pegboard Test (Fine motor skills), Children’s Memory Scale (CMS) Stories subtasks (Auditory/verbal memory), Wechsler Intelligence Scale for Children III (Intelligence), Woodcock-Johnson III (Academic achievement), Stroop Color-Word Test (Interference), Ruff 2-&-7 Selective Attention test (Selective attention), Verbal fluency (Cognitive flexibility), Tapping. The obtained g-factor represents 23.6% of the observed variance. The g-factor was then transformed to a z-score using the mean of 0.051 and the SD of 3.799.

In CaG cohort, the g-factor was computed using three cognitive tests: verbal and numeric reasoning (fluid intelligence), paired associates learning (episodic memory) and reaction time based on two-choice items. The obtained g-factor represents 43.2% of the variance observed in the cohort. The g-factor was then transformed to a z-score using the mean of -8.681×10^-16^ and the SD of 1.084.

In G-Scot cohort, the g-factor is based on four cognitive tests measuring processing speed, verbal declarative memory, executive functions and vocabulary. The g-factor represents 42.3% of the observed variance. The g-factor was then transformed to a z-score using the mean of -3.649×10^-16^ and the SD of 1.3.

In Imagen, the g-factor was computed using 4 cognitive measures: similarities score, vocabulary score, block design score, matrix reasoning score (WISC-IV). The obtained g-factor represents 57.3% of the observed variance. The g-factor was then transformed to a z-score using the mean of 9.626×10^-17^ and the SD of 1.514.

Note that the g-factor is a robust measure of general cognitive ability that is not very sensitive to the exact subtest used to calculate it as long as these measure a wide range of cognitive abilities. Moreover, the mean of the cognitive ability levels for the different cohorts were likely different, but our results from the meta-analysis (Fig. 1) suggest that the standard deviations obtained across cohorts were very similar as we observed similar effect sizes of CNV impact.

#### 3. Genetic information

##### 3.1. CNV calling

No CNV calling was performed in Sainte-Justine CNV-familial cohort since this cohort focuses only on the CNV investigated in the clinic for which the family was recruited.

In the MSSNG database*[30]*, 10,032 individuals were sequenced at multiple sites using Illumina sequencing HiSeq, HiSeq 2,500 or HiSeqX. Next generation sequencing data were analysed using Broad institute Genome Analysis ToolKit (GATK) best practices*[31]*. For MSSNG, read alignment data were used to compute CNV calling following the workflow presented in Trost *et al.* *[32]*. We filtered CNVs, by selecting only the CNVs overlapping at least 10 probes of each of the array technologies used by the different cohorts.

For all other datasets, we used the same methodology as in Huguet *et al.**[3]*. For all cohorts, we used stringent quality-control criteria: call rate ≥95%; log R ratio-standard deviation <0.35; B allele frequency-standard deviation <0.08 and |wave factor|<0.05. The probes coordinates were updated from hg18 to hg19 using Illumina information and the liftover tool from the genome browser.

CNVs detected by PennCNV*[33]* and QuantiSNP*[34]* algorithms were combined (CNVision)*[35]* to minimize the number of potential false discoveries. A manual visualisation of some CNVs showed a high number of false positive CNVs in G-Scot, LBC1936 and CaG cohorts; thus, only the CNVs detected by the both algorithms were considered in these three cohorts. After this merging step, the CNV inheritance analysis algorithm (in-house algorithm), was applied to concatenate adjacent CNVs (same type) into one, according to the following criteria: a) gap between CNVs ≤150 kb; b) size of the CNVs ≥ 1000 bp; and c) number of probes ≥ 3. After these steps, we remove from the analyses, all arrays for which a suspiciously high number of CNVs has been detected (≥ 50 for low resolution arrays [<1 million probes] and ≥ 200 for high resolution arrays [≥1 million probes]).

##### 3.2. CNV filtering

After filtering array according to their quality, we applied filtering for autosomal CNVs (X-linked CNVs were not investigated in this study). The CNVs with the following criteria were selected for analysis: confidence score ≥ 30 (with at least one of both detection algorithms), size ≥ 50 kb, unambiguous type (deletions or duplications) and overlap with segmental duplicates, HLA regions or centromeric regions < 50%. Moreover, we applied an in-house algorithm based on the random forest method to detect additional artefactual CNVs. This algorithm, based on several strategies (bagging, boosting) and on 9 CNV characteristics (Array criteria: log R ratio-standard deviation, B allele frequency-standard deviation, wave frequency; Localization CNV criteria: % of CNV overlap with centromeric regions and with segmental duplications; CNV criteria: density SNPs (size of CNV / numbers of SNPs), confidence score, % algorithms overlapping, type of CNV) was trained and tested respectively on 70% and 30% of a total of 22,154 CNVs (19,647 true CNVs and 2,507 artefacts from SSC and G-Scot cohort) and manually visualized by at least two individuals representing a high confidence reference training set for the random forest model. Its application on the test set showed a sensitivity of >0.82 and a specificity >0.88 whatever the strategy. The best strategy was then chosen and tested again on an additional naive dataset of 3,181 CNVs (2808 true CNVs and 373 artefacts from Imagen and SYS cohorts) and showed a sensitivity of 0.89 and a specificity of 0.8.

For the mega-analysis, we applied an additional filtering, by selecting only the CNVs overlapping at least 10 probes of each of the array technologies used by the different cohorts, before pooling the data.

##### 3.3. Genetic analysis of pairwise relatedness

For each cohort, we checked sex genotype and Mendelian error to validate basic information such as sex and parenthood. Also the classical multidimensional scaling (MDS) was used to identify relatedness based on the identity by state (IBS) matrices of genetic distances. These analyses were done using PLINK*[36]* (pngu.mgh.harvard.edu/purcell/plink/). These calculations are based on SNPs that were filtered to keep only autosomal SNPs with minor allele frequency (MAF) > 5% and with good quality, significance threshold for a test of Hardy-Weinberg equilibrium < 1×10^-6^ and missing genotype rates < 10%.

##### 3.4. Criteria to remove outlier individual with particular CNV

We excluded from the analyses all individuals carrying a large structural variant ≥10Mb, a mosaic CNV or an anomaly on the X or Y chromosomes. Thus, we removed 2 individuals from Imagen, 1 from SYS, 2 from CaG, 39 from G-Scot and 3 from MSSNG.

##### 3.5. Definition of recurrent CNVs

We compiled a list of 121 CNVs proposed to be pathogenic in widely accepted sources[37–41], presented with the region coordinates in Huguet *et al*.*[3]* We defined a CNV as recurrent if it overlaps ≥ 40% with one of these 121 CNVs and includes the key genes of this region (if known).

##### 3.6. Annotation of CNVs

We annotated the CNVs using Gencode V19 annotation (the reference release for hg19 Human genome release) with ENSEMBL gene name (<https://grch37.ensembl.org/index.html>). Briefly, we used bedtools suite (<https://bedtools.readthedocs.io/en/latest/>) to compare the different elements of the genes that overlap CNVs (UTRs, start and stop codons, exons and introns). Thus, for each CNV, we obtained its proportion of overlap with each gene as well as with each gene component (UTR, Coding exons, start and stop codons, introns).

Genes were annotated using different scores: the probability to be loss-of–function (LoF) intolerant (pLI), the residual variation intolerance score (RVIS), the CNV intolerance score (DEL/DUP score)[42–44], the upper 95*^th^* CI boundary of the ratio of observed over expected number of LoF mutations (LOEUF), a score developed by gnomAD*[45]* (<https://gnomad.broadinstitute.org/>), the number of protein-protein interactions (PPI)*[46]*, the differential stability (DS) score*[47]* as well as two lists defining genes of likely interest for IQ, including postsynaptic density (PSD) of the human cortex*[48]* and genes regulated by FMRP*[49]*. Also, we used a list of 256 ID-genes*[50, 51]*, previously identified with an excess of *de-novo* mutations in NDD cohorts. Non-coding regions were annotated with the number of expression quantitative trait loci (eQTLs) regulating genes expressed in the brain*[52]*. CNV scores were derived by summing all scores within CNVs. More details about this methodology may be found in Huguet *et al.[3]*.

#### 4. Statistical analyses

##### 4.1. Coverage

The number of different genes totally included in at least one CNV was first calculated in our total cohort of 24,092 subjects (Table 1) and then estimated in relationship with sample size. We randomly ordered the 24,092 subjects of our total cohort and counted, for each step of X new additional individuals (X=4000 for whole cohort or general population and X=1000 for ASD population), the number of different genes totally included in at least one CNV. The final curve was estimated by averaging counts obtained from 500 iterations of this procedure and 95% confidence intervals were estimated as the 2.5*^th^* and 97.5*^th^* quantiles of values resulting from the 500 iterations. This procedure was applied to all CNVs as well as to deletions and duplications separately. We also split the information according to the pLI (genes with pLI >0.9 versus genes with pLI ≤0.9) or according to LOEUF (genes with LOEUF <0.35 versus genes with LOEUF ≥ 0.35).

Then, we predicted how the number of different genes totally included in at least one CNV should evolve when including new individuals using a regression model on the results obtained above. We modeled the number of different genes as a function of number of individuals included in the study as follows:

Number of different genes ~ β_0_ + β_1_ × log(N) + β_2_ × [log(N)]^2^

where N is the number of individuals included in the study. The regression parameters β_0_, β_1_ and β_2_ where obtained from the whole cohort or from subsets divided into general population and autism population (Supplementary Fig. 8).

##### 4.2. Assessing age and sex biases in the cohorts

Since each cohort has specific biases based on their respective ascertainment methods, we first adjusted general intelligence for age and sex within each cohort before performing meta- and mega-analyses.

###### Adjusting for age within samples of unselected populations assessed with a g-factor.

We adjusted g-factor for age in CaG, G-Scot and SYS parents cohorts separately by following these steps:

- We performed a linear regression model to fit g-factor according to the age. Age was incorporated alone (linear effect) or with its square (quadratic effect) depending on the cohort.
- The final model could be written as: g-factor = *a* + *f*(age) + *ε* where *f*(age) = *b*×age (linear effect) or *f*(age) = *b_1_* × age + *b_2_* ×age^2^ (quadratic effect) with *a*, *b*, *b_1_*, *b_2_* the regression coefficients and *ε* the normally distributed error term.
- We computed the age-adjusted g-factor as the g-factor that each individual would have at the mean age of the cohort: age-adjusted g-factor = *a* + *f*(mean age of the cohort) + *ε*

(In other words, this corresponds to the g-factor predicted by the model for the mean age of the cohort + the residual for each individual).

The retained models for adjusting g-factor on age in CaG, G-Scot and SYS parents cohorts were obtained using polynomial regression of order 2 (quadratic effect of the age) in CaG and G-Scot and a simple linear regression in SYS parents. The non-adjusted g-factor and the adjusted g-factor are both displayed in Supplementary Fig. 9.

###### Adjusting for biological sex within autism cohorts

Autism cohorts are biased for sex with a higher number of males than females and a lower IQ for the latter. Thus, in SSC and MSSNG dataset, we adjusted IQ for sex as follows for each cohort:

- We performed an ANOVA analysis to fit IQ according to sex: IQ = *a* + *b*×**1**_(sex=female)_ + *ε* with *a* and *b* the regression coefficients, *ε* the normally distributed error term and **1**_(sex=female)_ the indicator variable equaling 1 if female and 0 if male.
- The sex-adjusted IQ corresponds to the IQ of each participant considered as a male: sex-adjusted IQ = *a* + *e*.

Results of models are displayed in Supplementary Fig. 10.

All the models presented after are based on this adjusted z-score.

In autism cohorts, we did not adjust for age as we adjusted for the type of test used for assessing IQ that already takes into account the age (as well as language level) of the individual. In other cohorts z-scores of general intelligence were not adjusted for age or sex as IQ scores are already adjusted for these two parameters. However, with the exceptions of Imagen and LBC1936 cohorts (there is no variability of age in these two cohorts), in all other cohorts the z-scores have been adjusted for age.

Moreover, models were not adjusted for ancestry components since they did not impact the effect of genetic scores on general intelligence in our previous studies (Huguet *et al.[3]*, Douard *et al.[53]*).

However other adjustments were made specifically inside each sample in meta-analysis or over all samples in mega-analysis.

##### 4.3. Meta-analyses assessing the effect of pLI on general intelligence

Firstly, models were computed independently for each sample (Supplementary Table 2). Outcome of interest was either the adjusted z-score for IQ or the adjusted g-factor (when IQ was not available). Predictor of interest was the pLI score measured at individual level for deletions and duplications separately.

Additional adjustment by sample:

- Imagen: no more adjustment.
- SYS child: this cohort includes siblings, thus we added a random effect in the model taking into account the correlation between individuals from the same family.
- SYS parents: no more adjustment.
- CaG: no more adjustment.
- G-Scot: this cohort includes individuals from the same family and we added a random effect in the model taking into account the correlation between individuals from the same family.
- LBC1936: no more adjustment.
- SSC: this cohort includes individuals from a large range of young age and heterogeneous language levels such that IQ was assessed with 5 different tests that are linked to age and IQ measure leading to an artificial age effect on IQ. To minimize differences associated with age, we adjusted models for which IQ test was used.
- MSSNG: as for SSC, we adjusted for the IQ test used (for 6 different tests). Moreover, this cohort includes individuals from the same family and we added a random effect in the model taking into account the correlation between individuals from the same family.

Secondly, meta-analyses were performed, separately for deletions and duplications, on summary results from each model, specifically on the estimate of regression coefficient associated with the predictor of interest (pLI) (Fig. 1).

##### 4.4. Mega-analysis assessing the effect of pLI or 1/LOEUF on general intelligence

We performed linear regression analyses to measure the impact of CNVs (pLI or 1/LOEUF individual’s scores) on general intelligence on the pooled dataset. The model was adjusted for the combined variable “cohort - type of intelligence measure” (as fixed effect) and family identifier as random effect. Estimates were obtained using the restricted maximum likelihood maximization.

We computed the partial R^2^ associated with pLI in a model that does not include the random effect and confirmed it in the subset of unrelated individuals.

Different sensitivity analyses were performed, including a modification of the manner to take into account random effect, the subset of populations (all, unrelated individuals, autism population, general population) and effect of deletions and duplications taking into account separately or in the same model (Supplementary Tables 9 and 10).

###### Comparing with other annotation scores as a predictor of the general intelligence in the pooled dataset.

We explored if the pLI was the best predictor of impact of deletions or duplications on general intelligence by comparing models based on the pLI with models based on other annotation scores. Models were compared on Akaike’s information criteria (AIC) obtained after fitting each model using likelihood maximization. The better fit is obtained for lower AIC (Supplementary Table 4).

###### Sensitivity analysis to assess the effect of pLI (resp. 1/LOEUF) after removing individuals with higher measure of pLI (resp. 1/LOEUF)

We performed sensitivity analyses by removing incrementally individuals with total pLI > 10, > 5, >3, >2, >1.5 and >1 point. We did the same for 1/LOEUF with the following threshold: total 1/LOEUF >60, 40, 20, 10, 4, 2.85.

###### Linear versus non–linear effect of haploinsufficiency scores

We tested non-linearity of the effect of haploinsufficiency scores on general intelligence by fitting different models on the pooled dataset. First, we introduced the pLI score as well as its square in the same model. Then, we proposed to explore a smooth function of the effect of pLI (resp. 1/LOEUF) using a kernel regression method. We used the Gaussian kernel function as it is flexible enough to account for various types of effects. We defined the Gaussian kernel function such as similarity between two subjects i and j is:

Gaussian(Z_i_, Z_j_, ρ) = exp(-||Z_i_ – Z_j_||^2^)/ρ)

where Z_i_, Z_j_ are scores (total pLI or total 1/LOEUF) for subjects i and j, ||.|| represents the Euclidean distance and ρ is a tuning parameter defining the smoothness of the function. Several values of the ρ parameter were compared (Supplementary Table 9 and 10).

###### Model including information on known ID-genes

CNVs haploinsufficiency scores were divided into sum of haploinsuffiency scores for ID-genes versus non ID-genes; models were then recomputed with the two explanatory variables (sum of scores for ID-genes and sum of scores of non ID-genes included in the CNV) for deletions and duplications separately (Supplementary Table 12).

##### 4.5. Concordance analysis between prediction of our models and literature observations

We examined if our prediction of the effect of haploinsufficiency on cognition was improved using the results of our meta-analysis. To do so, we calculated the concordance between model predictions and empirically measured loss of IQ for 47 known recurrent CNVs obtained from previous publications (Fig. 4, Supplementary Fig. 5, Supplementary Tables 14 and 15) or from UKbiobank study*[54]*. The concordance was computed using the intraclass coefficient correlation of type (3,1) (ICC_(3,1)_)*[55]*.

##### 4.6. Estimation of the probability of a *de novo* CNV using haploinsufficiency scores

We performed logistic regressions to estimate the probability of a CNV being *de novo* using the haploinsufficiency scores (pLI or 1/LOEUF scores, accounting for the effect of ID-genes) or estimated IQ loss as explanatory variables. We trained these *de novo* models on datasets for which mode of transmission of the CNVs was available (inherited or *de novo*). For these analyses, we added two clinical populations (Decipher (https://decipher.sanger.ac.uk/) and the cytogenetic database of Sainte-Justine Hospital and applied the same filtering as for the previous CNV selection. We included a total of 26,437 CNVs from two general populations: 810 individuals from G-Scot and 723 individuals from SYS; and four clinical populations: 3,919 individuals from the SSC, 956 individuals from MSSNG, 10,126 individuals from Decipher and 1,560 individuals from the cytogenetic database of Sainte-Justine Hospital (Supplementary Table 16). The binary outcome variable was the type of transmission (1=*de novo*, 0=inherited). Models for deletions and duplications were tested independently.

To investigate if the associations between *de novo* status and the haploinsufficiency scores were different for deletions and duplications, we also performed a logistic regression model combining the deletions and duplications scores in one explanatory variable (e.g. the IQ loss estimated with 1/LOEUF) and added an interaction between the type of CNV (deletion or duplication) and the haploinsufficiency score.

Then, we validated these models by comparing the model estimates with the percentage of *de novo* computed with Decipher (<https://decipher.sanger.ac.uk/>) for 27 recurrent CNVs (Supplementary Table 14). We used the estimates from models trained on a dataset that excluded these recurrent CNVs.

##### 4.7. Estimating the effect size of genes categories by LOEUF

We coded the information in the LOEUF score using 4 categories of tolerance for the genes: highly intolerant genes (LOEUF <0.2; n=980), moderately intolerant genes (0.2≤LOEUF<0.35 n=1,762), tolerant genes (0.35≤LOEUF<1; n=7,442) and highly tolerant genes (LOEUF≥1; n=8,267). The numbers of genes within each category were used in a linear model as four explanatory variables of the general intelligence.

Then, we explored whether particular score ranges within LOEUF were associated with unique effect sizes by using a sliding window across the range of scores spanned by LOEUF. We defined 2 variables: number of genes inside and outside the window, and considered them as predictors of the general intelligence in a linear model. We slid a window of size 0.15 LOEUF units, in increments of 0.05 units, thereby creating 38 different analyses.

For both methods, the adjusted z-score measure of general intelligence was the dependent variable and annotation scores, with the new codings, were the independent variables. All models were adjusted for “type of test-cohort” as fixed effects and for familial relationship as random effect; and analyses were performed for deletions and duplications separately.

##### 4.8. R packages used for statistical analyses

Statistical analyses were performed using R version 3.1.1. (R Development Core Team (2005). R: A language and environment for statistical computing. R Foundation for Statistical Computing, Vienna, Austria. ISBN 3-900051-07-0, URL: http://www.R-project.org.) Statistical analyses were performed using “nlme”*[56]*, “bootStepAIC”*[57]*, “KSPM” and “psych”*[58]* packages for mixed effect models, bootstrap for variable selection procedure, kernel semi-parametric models and ICC_(3,1)_ respectively.

### Extended Data Tables

| **Cohorts** | **Deletions** | | | **Duplications** | | |
| --- | --- | --- | --- | --- | --- | --- |
|  | **Est.** | **SE** | ***P*** | **Est.** | **SE** | ***P*** |
| **Imagen** | -0.234 | 0.091 | 1.02x10^-02^ | 0.083 | 0.043 | 5.56x10^-02^ |
| **SYS-child** | -0.169 | 0.054 | 1.94x10^-03^ | -0.079 | 0.040 | 4.74x10^-02^ |
| **SYS-parent** | -0.334 | 0.160 | 3.78x10^-02^ | 0.019 | 0.057 | 7.33x10^-01^ |
| **LBC1936** | -0.277 | 0.127 | 2.99x10^-02^ | 0.026 | 0.083 | 7.51x10^-01^ |
| **CaG-GSA** | -0.024 | 0.135 | 8.61x10^-01^ | -0.105 | 0.037 | 4.53x10^-03^ |
| **CaG-Omni2.5** | -0.321 | 0.250 | 1.99x10^-01^ | -0.074 | 0.056 | 1.83x10^-01^ |
| **G-Scot** | -0.179 | 0.031 | 6.03x10^-09^ | -0.066 | 0.014 | 2.24x10^-06^ |
| **SSC-1MV1** | -0.155 | 0.068 | 2.30x10^-02^ | -0.022 | 0.091 | 8.06x10^-01^ |
| **SSC-1MV3** | -0.176 | 0.044 | 6.64x10^-05^ | -0.062 | 0.023 | 7.12x10^-03^ |
| **SSC-Omni2.5** | -0.160 | 0.043 | 1.94x10^-04^ | -0.055 | 0.038 | 1.46x10^-01^ |
| **MSSNG** | -0.200 | 0.065 | 2.15x10^-03^ | -0.013 | 0.036 | 7.21x10^-01^ |

**Supplementary Table 1.** Estimates associated with pLI within each sample included in the meta-analyse.

Models were computed separately for deletions and duplications. The adjusted z-scored measure of general intelligence was the dependent variable and pLI was the independent one. All models were adjusted differently as detailed in supplementary material. SYS: Saguenay youth study; SSC: Simon’s simplex collection; CaG: Cartagene; LBC: Lothian birth cohort; pLI: probability of loss-of-function intolerance; Est.: estimate; SE: standard error; *P*: p-value.

| **Cohort** | **Phenotype** | **Deletions** | | | **Duplications** | | |
| --- | --- | --- | --- | --- | --- | --- | --- |
|  |  | **Est.** | **SE** | ***P*** | **Est.** | **SE** | ***P*** |
| **Imagen (n=1,720)** | **z-scored NVIQ** | -0.233 | 0.091 | 0.011 | 0.086 | 0.044 | 0.048 |
|  | **z-scored g-factor** | -0.222 | 0.092 | 0.017 | 0.004 | 0.044 | 0.930 |
| **SYS-child (n=857)** | **z-scored NVIQ** | -0.146 | 0.055 | 0.009 | -0.066 | 0.042 | 0.114 |
|  | **z-scored g-factor** | -0.150 | 0.054 | 0.006 | -0.063 | 0.041 | 0.130 |
| **LBC1936 (n=504)** | **z-scored IQ Moray** | -0.277 | 0.127 | 0.030 | 0.026 | 0.083 | 0.751 |
|  | **z-scored g-factor** | -0.263 | 0.131 | 0.045 | -0.011 | 0.086 | 0.901 |

**Supplementary Table 2.** Comparison of estimates associated with pLI when measuring the effect-size of CNVs on IQ and g-factor.

Models were computed separately for deletion or duplication. The z-scored measure of general intelligence (unadjusted) was the dependent variable and annotation score was the independent one. In SYS, the model was adjusted for age as fixed effect and on familial relationship as random effect. SYS: Saguenay youth study; LBC: Lothian birth cohort; pLI: probability of loss-of-function intolerance; Est.: estimate; SE: standard error; *P*: p-value.

| **Population (n=)** | **Category of genes** | **Deletion** | | **Duplication** | |
| --- | --- | --- | --- | --- | --- |
|  |  | **Est. (SE)** | ***P*** | **Est. (SE)** | ***P*** |
| ***Models based on pLI*** | | | | | |
| Probands only (132) | All Genes | -0.090 (0.020) | 2.73×10^-5^ | -0.043 (0.019) | 0.023 |
| All (551) | All Genes | -0.147 (0.018) | 1.1×10^-15^ | -0.069 (0.016) | 8.7 ×10^-6^ |
| Probands only (132) | ID genes | -0.477 (0.193) | 0.01 | -0.399 (0.386) | 0.30 |
|  | Non-ID genes | -0.072 (0.021) | 1.01×10^-3^ | -0.036 (0.027) | 0.18 |
| All (551) | ID genes | -0.688 (0.175) | 8.8×10^-5^ | -0.445 (0.390) | 0.25 |
|  | Non-ID genes | -0.117 (0.021) | 1.7×10^-8^ | -0.065 (0.021) | 2.1×10^-3^ |
| ***Models based on 1/LOEUF*** | | | | | |
| Probands only (132) | All Genes | -0.014 (0.003) | 3.08×10^-5^ | -0.007 (0.003) | 0.03 |
| All (551) | All Genes | -0.024 (0.003) | 1.1×10^-15^ | -0.011 (0.002) | 1.6×10^-5^ |
| Probands only (132) | ID genes | -0.061 (0.027) | 0.03 | -0.079 (0.050) | 0.11 |
|  | No ID genes | -0.012 (0.003) | 1.11×10^-3^ | -0.005 (0.004) | 0.21 |
| All (551) | ID genes | -0.077 (0.026) | 3.0×10^-3^ | -0.088 (0.055) | 0.11 |
|  | No ID genes | -0.021 (0.003) | 7.0×10^-10^ | -0.009 (0.003) | 2.3×10^-3^ |

**Supplementary Table 3.** Estimates obtained for models performed in Ste-Justine neurodevelopmental cohort.

The model was computed simultaneously for deletions and duplications for each annotation pLI or 1/LOEUF score. The z-scored measure of general intelligence (unadjusted) was the dependent variable and annotation score was the independent one. All models were adjusted for type of test and sex as fixed effects and for familial kinship matrix as random effect. Est.: estimates; SE: standard error; *P*: p-value; pLI: probability of loss-of-function intolerance; LOEUF: the upper 95th CI boundary of the ratio of observed over expected number of loss-of-function mutations.

| **Variables** | **Deletions** | | | **Duplications** | | |
| --- | --- | --- | --- | --- | --- | --- |
|  | **Est. (SE)** | ***P*** | **AIC** | **Est. (SE)** | ***P*** | **AIC** |
| **No. of genes** | -0.037 (0.004) | 4.43×10^-21^ | 70,348.86 | -0.011 (0.002) | 9.10×10^-8^ | 70,409.34 |
| **pLI *** | -0.176 (0.016) | 1.25×10^-28^ | 70,314.15 | -0.054 (0.009) | 1.90×10^-9^ | 70,401.81 |
| **pLI 2019 †** | -0.194 (0.018) | 3.04×10^-26^ | 70,325.13 | -0.062 (0.010) | 3.09×10^-9^ | 70,402.76‡ |
| **1/(O/E) †** | -0.015 (0.001) | 1.48×10^-28^ | 70,314.49‡ | -0.005 (0.001) | 3.45×10^-9^ | 70,402.97‡ |
| **1/LOEUF †** | -0.029 (0.003) | 7.14×10^-29^ | 70,313.03‡ | -0.009 (0.002) | 8.65×10^-9^ | 70,404.76‡ |
| **RVIS** | -0.001 (8.4×10^-5^) | 1.70×10^-14^ | 70,378.94 | -2.0×10^-4^ (4.6×10^-5^) | 9.12×10^-6^ | 70,418.24 |
| **DEL/DUP score** | -0.023 (0.003) | 1.58×10^-16^ | 70,369.69 | -0.008 (0.002) | 3.25×10^-6^ | 70,416.25 |
| **PPI** | -1.3×10^-4^ (5.0×10^-5^) | 0.007 | 70,430.68 | -7.7×10^-5^ (3.2×10^-5^) | 0.015 | 70,432.07 |
| **DS** | -0.094 (0.009) | 1.62×10^-26^ | 70,323.88 | -0.030 (0.005) | 4.5×10^-9^ | 70,403.49 |
| **PSD genes** | -0.330 (0.040) | 4.40×10^-18^ | 70,362.57 | -0.114 (0.022) | 1.46×10^-7^ | 70,410.26 |
| **FMRP genes** | -0.376 (0.050) | 6.34×10^-14^ | 70,381.54 | -0.162 (0.031) | 1.73×10^-7^ | 70,410.59 |
| **eQTL** | -0.001 (1.1×10^-4^) | 1.68×10^-17^ | 70,365.23 | -2.6×10^-4^ (5.6×10^-5^) | 2.57×10^-6^ | 70,415.80 |

**Supplementary Table 4.** Comparison of model fit for general intelligence according to annotation score.

The model was computed separately for each annotation score. The adjusted z-scored measure of general intelligence was the dependent variable and annotation score was the independent one. All models were adjusted on “type of test-cohort” variable as fixed effect and on familial relationship as random effect. Estimates, SE and *P* were computed using REML approach whereas AIC was computed using ML approach. A lower AIC means a better fit. ML: maximum likelihood; REML: restricted maximum likelihood; Est.: estimate; SE: standard error; *P*: p-value; AIC: Akaike’s information criteria; No.: number; pLI: probability of being loss-of-function intolerant; O/E: observed over expected; LOEUF: the upper 95th confidence interval boundary of the ratio of observed over expected number of loss-of-function mutation score; RVIS: residual variation intolerance score; DEL score: the CNV intolerance score for deletion; DUP score: the CNV intolerance score for duplication; PPI: protein-protein interaction; DS: differential stability; PSD: number of genes in postsynaptic density function category; FMRP: number of genes regulated by FMRP; eQTL: the number of SNP regulating genes expressed in the brain (expression quantitative trait loci (eQTLs)). Details about annotation scores are available at section 3.6 (p.); * pLI variable used as reference; † other GNOMAD scores measuring similar concept than pLI variable; ‡ indicates AICs lower than or closed to the AIC obtained with pLI variable.

| **Model** | **Variables** | **Deletions** | | | **Duplications** | | |
| --- | --- | --- | --- | --- | --- | --- | --- |
|  |  | **Est.** | **SE** | ***P*** | **Est.** | **SE** | ***P*** |
| **All cohorts (n=24,092)** | **(Intercept)** | -1.244 | 0.034 | 5.94x10^-273^ | -1.256 | 0.035 | 6.28x10^-275^ |
|  | **pLI** | -0.169 | 0.024 | 4.89x10^-12^ | -0.048 | 0.014 | 7.98x10^-4^ |
|  | **pLI** x **Age** | -6.76x10^-5^ | 8.19x10^-5^ | 4.09x10^-1^ | -2.77x10^-5^ | 3.33x10^-5^ | 4.06x10^-1^ |
|  | **Sex (female)** | 0.100 | 0.014 | 4.05x10^-13^ | 0.100 | 0.014 | 4.63x10^-13^ |
|  | **Age** | -7.0x10^-4^ | 4.55x10^-5^ | 1.01x10^-52^ | -6.92x10^-4^ | 4.58x10^-5^ | 6.16x10^-51^ |
| **Unselected population (n=20,151)** | **(Intercept)** | 0.266 | 0.035 | 1.67x10^-14^ | 0.263 | 0.035 | 5.07x10^-14^ |
|  | **pLI** | -0.167 | 0.059 | 4.65x10^-3^ | -0.034 | 0.030 | 2.60x10^-1^ |
|  | **pLI** x **Age** | -4.09x10^-5^ | 1.22x10^-4^ | 7.37x10^-1^ | -4.64x10^-5^ | 5.13x10^-5^ | 3.66x10^-1^ |
|  | **Sex (female)** | 0.133 | 0.013 | 3.66x10^-23^ | 0.134 | 0.013 | 1.85x10^-23^ |
|  | **Age** | -5.66x10^-4^ | 4.29x10^-5^ | 2.68x10^-39^ | -5.53x10^-4^ | 4.35x10^-5^ | 8.21x10^-37^ |
| **ASD population (n=3,941)** | **(Intercept)** | -0.177 | 0.063 | 5.19x10^-3^ | -0.171 | 0.064 | 7.56x10^-3^ |
|  | **pLI** | -0.138 | 0.072 | 5.53x10^-2^ | -0.060 | 0.046 | 1.96x10^-1^ |
|  | **pLI** x **Age** | -1.36x10^-4^ | 5.54x10^-4^ | 8.07x10^-1^ | 1.59x10^-4^ | 3.77x10^-4^ | 6.74x10^-1^ |
|  | **Sex (female)** | -0.242 | 0.061 | 8.76x10^-5^ | -0.254 | 0.061 | 4.10x10^-5^ |
|  | **Age** | -0.013 | 0.001 | 1.33x10^-62^ | -0.013 | 0.001 | 1.24x10^-62^ |

**Supplementary Table 5.** Linear regression models including pLI x age interaction as predictor of general intelligence.

The model was computed separately for for deletions and duplications. The z-scored measure of general intelligence (unadjusted) was the dependent variable and pLI (total sum by individual), age (in months) and sex (male as reference level) as well as the pLI x age interaction were the independent ones. All models were adjusted on “type of test-cohort”, as fixed effect and on familial relationship as random effect. Est.: estimates; SE: standard error; *P*: p-value; SYS: Saguenay youth study; LBC: Lothian birth cohort; CaG: Cartagene; G-Scot: generation Scotland; SSC: Simon simplex collection; pLI: probability of loss-of-function intolerance.

| **Model** | **Variables** | **Deletions** | | | **Duplications** | | |
| --- | --- | --- | --- | --- | --- | --- | --- |
|  |  | **Est.** | **SE** | ***P*** | **Est.** | **SE** | ***P*** |
| **All cohorts (n=24,092)** | **(Intercept)** | -1.240 | 0.035 | 6.04x10^-271^ | -1.257 | 0.035 | 6.74x10^-275^ |
|  | **1/LOEUF** | -0.031 | 0.004 | 4.16x10^-14^ | -0.008 | 0.003 | 2.26x10^-3^ |
|  | **1/LOEUF** x **Age** | 3.51x10^-7^ | 1.36x10^-5^ | 9.79x10^-1^ | -4.40x10^-6^ | 5.56x10^-6^ | 4.29x10^-1^ |
|  | **Sex (female)** | 0.100 | 0.014 | 2.95x10^-13^ | 0.100 | 0.014 | 5.07x10^-13^ |
|  | **Age** | -7.01x10^-4^ | 4.55x10^-5^ | 8.89x10^-53^ | -6.92x10^-4^ | 4.59x10^-5^ | 1.03x10^-50^ |
| **Unselected population (n=20,151)** | **(Intercept)** | 0.266 | 0.035 | 1.73x10^-14^ | 0.265 | 0.035 | 3.58x10^-14^ |
|  | **1/LOEUF** | -0.027 | 0.010 | 6.10x10^-3^ | -0.005 | 0.005 | 2.95x10^-1^ |
|  | **1/LOEUF** x **Age** | -9.07x10^-7^ | 2.03x10^-5^ | 9.64x10^-1^ | -7.98x10^-6^ | 8.25x10^-6^ | 3.33x10^-1^ |
|  | **Sex (female)** | 0.133 | 0.013 | 2.96x10^-23^ | 0.134 | 0.013 | 1.95x10^-23^ |
|  | **Age** | -5.65x10^-4^ | 4.30x10^-5^ | 3.77x10^-39^ | -5.52x10^-4^ | 4.36x10^-5^ | 2.05x10^-36^ |
| **ASD population (n=3,941)** | **(Intercept)** | -0.172 | 0.063 | 6.63x10^-3^ | -0.170 | 0.065 | 8.41x10^-3^ |
|  | **1/LOEUF** | -0.029 | 0.012 | 1.60x10^-2^ | -0.010 | 0.008 | 2.05x10^-1^ |
|  | **1/LOEUF** x **Age** | 1.72x10^-5^ | 9.01x10^-5^ | 8.49x10^-1^ | 2.99x10^-5^ | 6.66x10^-5^ | 6.53x10^-1^ |
|  | **Sex (female)** | -0.239 | 0.061 | 1.06x10^-4^ | -0.255 | 0.061 | 3.93x10^-5^ |
|  | **Age** | -0.013 | 0.001 | 1.21x10^-62^ | -0.013 | 0.001 | 3.17x10^-62^ |

**Supplementary Table 6.** Linear regression models including 1/LOEUF x age interaction as predictor of general intelligence.

The model was computed separately =for deletion or duplication. The z-scored measure of general intelligence (unadjusted) was the dependent variable and 1/LOEUF (total sum by individual), age (in months) and sex (male as reference level) as well as the 1/LOEUF x age interaction were the independent ones. All models were adjusted on “type of test-cohort”, as fixed effect and on familial relationship as random effect. Est.: estimates; SE: standard error; *P*: p-value; SYS: Saguenay youth study; LBC: Lothian birth cohort; CaG: Cartagene; G-Scot: generation Scotland; SSC: Simon simplex collection; LOEUF: the upper 95^th^CI boundary of the ratio of observed over expected number of loss-of-function mutations.

| **Model** | **Variables** | **Deletions** | | | **Duplications** | | |
| --- | --- | --- | --- | --- | --- | --- | --- |
|  |  | **Est.** | **SE** | ***P*** | **Est.** | **SE** | ***P*** |
| **All cohorts (n=24,092)** | **(Intercept)** | -1.243 | 0.034 | 3.60x10^-273^ | -1.253 | 0.035 | 4.32x10^-275^ |
|  | **pLI** | -0.165 | 0.020 | 5.43x10^-16^ | -0.061 | 0.012 | 2.71x10^-7^ |
|  | **pLI** x **Sex (female)** | -0.049 | 0.032 | 1.30x10^-1^ | 0.010 | 0.018 | 5.90x10^-1^ |
|  | **Sex (female)** | 0.102 | 0.014 | 1.55x10^-13^ | 0.098 | 0.014 | 3.17x10^-12^ |
|  | **Age** | -7.02x10^-4^ | 4.55x10^-5^ | 4.87x10^-53^ | -6.96x10^-4^ | 4.56x10^-5^ | 4.98x10^-52^ |
| **Unselected population (n=20,151)** | **(Intercept)** | 0.265 | 0.035 | 1.83x10^-14^ | 0.268 | 0.035 | 1.00x10^-14^ |
|  | **pLI** | -0.147 | 0.039 | 1.67x10^-4^ | -0.074 | 0.018 | 6.31x10^-5^ |
|  | **pLI** x **Sex (female)** | -0.065 | 0.051 | 2.02x10^-1^ | 0.024 | 0.023 | 3.06x10^-1^ |
|  | **Sex (female)** | 0.135 | 0.013 | 1.57x10^-23^ | 0.130 | 0.014 | 4.17x10^-21^ |
|  | **Age months** | -5.67x10^-4^ | 4.28x10^-5^ | 1.06x10^-39^ | -5.60x10^-4^ | 4.28x10^-5^ | 8.59x10^-39^ |
| **ASD population (n=3,941)** | **(Intercept)** | -0.175 | 0.063 | 5.46x10^-3^ | -0.176 | 0.063 | 5.39x10^-3^ |
|  | **pLI** | -0.161 | 0.029 | 8.12x10^-8^ | -0.044 | 0.019 | 1.95x10^-2^ |
|  | **pLI** x **Sex (female)** | 0.023 | 0.053 | 6.68x10^-1^ | 0.007 | 0.037 | 8.39x10^-1^ |
|  | **Sex (female)** | -0.246 | 0.061 | 8.35x10^-5^ | -0.258 | 0.062 | 4.63x10^-5^ |
|  | **Age** | -0.013 | 0.001 | 2.91x10^-63^ | -0.013 | 0.001 | 1.83x10^-63^ |

**Supplementary Table 7.** Linear regression models including pLI x sex interaction as predictor of general intelligence.

The model was computed separately for deletions and duplications. The z-scored measure of general intelligence (unadjusted) was the dependent variable and pLI (total sum by individual), age (in months) and sex (male as reference level) as well as the pLI x sex interaction were the independent ones. All models were adjusted on “type of test-cohort” as fixed effect and on familial relationship as random effect. Est.: estimates; SE: standard error; *P*: p-value; SYS: Saguenay youth study; LBC: Lothian birth cohort; CaG: Cartagene; G-Scot: generation Scotland; SSC: Simon simplex collection; pLI: probability of loss-of-function intolerance.

| **Model** | **Variables** | **Deletions** | | | **Duplications** | | |
| --- | --- | --- | --- | --- | --- | --- | --- |
|  |  | **Est.** | **SE** | ***P*** | **Est.** | **SE** | ***P*** |
| **All cohorts (n=24,092)** | **(Intercept)** | -1.240 | 0.034 | 6.41x10^-272^ | -1.253 | 0.035 | 3.12x10^-275^ |
|  | **1/LOEUF** | -0.029 | 0.003 | 2.27x10^-17^ | -0.010 | 0.002 | 5.97x10^-7^ |
|  | **1/LOEUF** x **Sex (female)** | -0.003 | 0.005 | 5.85x10^-1^ | 0.002 | 0.003 | 5.03x10^-1^ |
|  | **Sex (female)** | 0.101 | 0.014 | 2.67x10^-13^ | 0.097 | 0.014 | 6.69x10^-12^ |
|  | **Age** | -7.01x10^-4^ | 4.55x10^-5^ | 6.78x10^-53^ | -6.96x10^-4^ | 4.56x10^-5^ | 4.91x10^-52^ |
| **Unselected population (n=20,151)** | **(Intercept)** | 0.266 | 0.035 | 1.68x10^-14^ | 0.271 | 0.035 | 5.72x10^-15^ |
|  | **1/LOEUF** | -0.027 | 0.007 | 1.12x10^-4^ | -0.012 | 0.003 | 7.58x10^-5^ |
|  | **1/LOEUF** x **Sex (female)** | -0.002 | 0.009 | 8.42x10^-1^ | 0.004 | 0.004 | 3.22x10^-1^ |
|  | **Sex (female)** | 0.133 | 0.013 | 6.70x10^-23^ | 0.130 | 0.014 | 9.82x10^-21^ |
|  | **Age** | -5.66x10^-4^ | 4.28x10^-5^ | 1.62x10^-39^ | -5.60x10^-4^ | 4.28x10^-5^ | 8.75x10^-39^ |
| **ASD population (n=3,941)** | **(Intercept)** | -0.173 | 0.063 | 5.95x10^-3^ | -0.176 | 0.063 | 5.47x10^-3^ |
|  | **1/LOEUF** | -0.028 | 0.005 | 1.87x10^-8^ | -0.007 | 0.003 | 2.64x10^-2^ |
|  | **1/LOEUF** x **Sex (female)** | 0.005 | 0.008 | 5.32x10^-1^ | 0.002 | 0.007 | 7.50x10^-1^ |
|  | **Sex (female)** | -0.246 | 0.061 | 8.67x10^-5^ | -0.261 | 0.063 | 4.35x10^-5^ |
|  | **Age** | -0.013 | 0.001 | 3.29x10^-63^ | -0.013 | 0.001 | 1.80x10^-63^ |

**Supplementary Table 8.** Linear regression models including 1/LOEUF x sex interaction as predictor of general intelligence.

The model was computed separately for deletions and duplications. The z-scored measure of general intelligence (unadjusted) was the dependent variable and 1/LOEUF (total sum by individual), age (in months) and sex (male as reference level) as well as the 1/LOEUF x sex interaction were the independent ones. All models were adjusted on “type of test-cohort” as fixed effect and on familial relationship as random effect. Est.: estimates; SE: standard error; *P*: p-value; SYS: Saguenay youth study; LBC: Lothian birth cohort; CaG: Cartagene; G-Scot: generation Scotland; SSC: Simon simplex collection; LOEUF: the upper 95th CI boundary of the ratio of observed over expected number of loss_of-function mutations.

| **Models** |  |  | **Deletions** | | **Duplications** | | |
| --- | --- | --- | --- | --- | --- | --- | --- |
| **Linear effect of pLI** | | | | | | | |
|  |  | **n=** | **Est. (SE)** | ***P*** | | **Est. (SE)** | ***P*** |
| Main models | Mixed effect models  (random effect on family identifier) | 24,092 | -0.176 (0.016) | 1.25 x 10^-28^ | | -0.054 (0.009) | 1.90 x 10^-9^ |
| Deletions and duplications effect in a unique model* | Mixed effect models  (random effect on family identifier) | 24,092 | -0.175 (0.016) | 1.46 x 10^-28^ | | -0.053 (0.009) | 2.22 x 10^-9^ |
| General population subset | Mixed effect models  (random effect on family identifier) | 20,151 | -0.181 (0.025) | 4.77 x 10^-13^ | | -0.058 (0.011) | 5.17 x 10^-7^ |
| Autism population subset | Mixed effect models  (random effect on family identifier) | 3,941 | -0.173 (0.026) | 2.61 x 10^-10^ | | -0.048 (0.017) | 5.60 x 10^-3^ |
| Familial effect was accounted for using kinship matrix | Mixed effect models  (kinship matrix) | 24,092 | -0.176 (0.016) | 9.77 x 10^-29^ | | -0.054 (0.009) | 1.85 x 10^-9^ |
| Unrelatives subset | Linear regression model in unrelated individuals | 14,874 | -0.174 (0.019) | 2.54 x 10^-20^ | | -0.043 (0.011) | 7.68 x 10^-5^ |
|  |  |  | **Derivative Mean/SD**  **(pLI Del.)** | ***P*** | | **Derivative Mean/SD**  **(pLI Dup.)** | ***P*** |
| Linear kernel function of pLI | Kernel semi-parametric model in unrelated individuals | 14,874 | -0.085/0.495  = -0.172 | 1.04 x 10^-12^ | | -0.035/0.861  = -0.041 | 8.41 x 10^-5^ |
| **Non linear effect of pLI** | | | | | | | |
|  |  | **n=** | **Est. for quadratic effect (SE)** | ***P*** | | **Est. for quadratic effect (SE)** | ***P*** |
| Quadratic effect of pLI | Mixed effect models  (random effect on family identifier) | 24,092 | -0.001 (0.004) | 0.85 | | -0.001 (0.001) | 0.46 |
|  |  |  | **Derivative mean/SD**  **(pLI Del.)** | ***P*** | | **Derivative mean/SD**  **(pLI Dup.)** | ***P*** |
| Gaussian kernel function of pLI (ρ=10) | Kernel semi-parametric model in unrelated individuals | 14,874 | -0.039/0.495  =-0.079 | 5.93 x 10^-11^ | | 0.010/0.861  =0.012 | 0.039 |

**Supplementary Table 9.** Linear regression models performed in the mega-analysis to measure the effect of deleted or duplicated units of pLI on general intelligence on general intelligence.

Models were performed separately for deletions and duplications except for the one marked with *. The dependent variable was the adjusted z-scored measure of general intelligence and pLI was the independent one. All models were adjusted on “type of test-cohort” variable as fixed effect and on familial relationship as random effect when relatives are included in the models. Est.: estimate; SE: standard error; *P*: p-value; SD: standard deviation; pLI: probability of being loss-of-function intolerant. Gaussian kernel was performed using ρ=0.1, ρ=1 and ρ=10. Only the Gaussian kernel model with the better fit according to AIC was displayed in the table. Of note, according to AIC to compare Gaussian and linear kernel, the linear kernel is the best choice for fitting these data. Mean derivatives divided by standard deviation of the covariate of interest may be interpreted as a beta regression coefficient when assessing the model with linear kernel. When Gaussian kernel is performed, this is just indicative.

| **Models** |  |  | **Deletions** | | **Duplications** | | |
| --- | --- | --- | --- | --- | --- | --- | --- |
| **Linear effect of 1/LOEUF** | | | | | | | |
|  |  | **n=** | **Est. (SE)** | ***P*** | | **Est. (SE)** | ***P*** |
| Main models | Mixed effect models  (random effect on family identifier) | 24,092 | -2.92×10^-2^ (2.61×10^-3^) | 7.14×10^-29^ | | -8.73×10^-3^ (1.52×10^-3^) | 8.65×10^-9^ |
| Deletions and duplications effect in a unique model | Mixed effect models  (random effect on family identifier) | 24,092 | -2.90×10^-2^ (2.60×10^-3^) | 1.14×10^-28^ | | -8.58×10^-3^ (1.51×10^-3^) | 1.40×10^-8^ |
| General population subset | Mixed effect models  (random effect on family identifier) | 20,151 | -2.73×10^-2^ (4.21×10^-3^) | 9.27×10^-11^ | | -9.13×10^-3^ (1.83×10^-3^) | 5.85×10^-7^ |
| Autism population subset | Mixed effect models  (random effect on family identifier) | 3,941 | -3.01×10^-2^ (4.29×10^-3^) | 2.56×10^-11^ | | -4.85×10^-2^ (1.73×10^-2^) | 5.60×10^-3^ |
| Familial effect was accounted for using kinship matrix | Mixed effect models  (kinship matrix) | 24,092 | -2.90×10^-2^ (3.00×10^-3^) | 5.54×10^-29^ | | -9.00×10^-3^ (2.00×10^-3^) | 8.48×10^-9^ |
| Unrelatives subset | Linear regression model in unrelated individuals | 14,874 | -2.95×10^-2^ (3.08×10^-3^) | 1.08×10^-21^ | | -7.21×10^-3^ (1.86×10^-3^) | 1.02×10^-4^ |
|  |  |  | **Derivative mean/SD**  **(1/LOEUF Del.)** | ***P*** | | **Derivative mean/SD**  **(1/LOEUF Dup.)** | ***P*** |
| Linear kernel function of 1/LOEUF | Kernel semi-parametric model  in unrelated individuals | 14,874 | -0.088/3.03  = -0.029 | 3.99×10^-13^ | | -0.034/5.02  = -0.007 | 1.17×10^-4^ |
| **Non linear effect of 1/LOEUF** | | | | | | | |
|  |  |  | **Est. for quadratic effect (SE)** | ***P*** | | **Est. for quadratic effect (SE)** | ***P*** |
| Quadratic effect of 1/LOEUF | Mixed effect models  (random effect on family identifier) | 24,092 | -2.50×10^-5^ (1.03×10^-4^) | 0.81 | | -1.58×10^-5^ (3.60×10^-5^) | 0.66 |
|  |  |  | **Derivative mean/SD**  **(1/LOEUF Del.)** | ***P*** | | **Derivative mean/SD**  **(1/LOEUF Dup.)** | ***P*** |
| Gaussian kernel function of 1/LOEUF (DEL: ρ=10; DUP: ρ=1) | Kernel semi-parametric model  in unrelated individuals | 14,874 | -0.073/3.03  = -0.024 | 1.29×10^-11^ | | -0.032/5.02  = -0.006 | 0.13 |

**Supplementary Table 10.** Linear regression models performed in the mega-analysis to measure the effect of deleted or duplicated units of 1/LOEUF on general intelligence.

Models were performed separately for deletions and duplications except for the one marked with *. The dependent variable was the adjusted z-scored measure of general intelligence and pLI was the independent one. All models were adjusted on “type of test-cohort” variable as fixed effect and on familial relationship as random effect when relatives are included in the models. SE: standard error; *P*: p-value; SD: standard deviation; LOEUF: the upper 95th CI boundary of the ratio of observed over expected number of loss-of-function mutations. Gaussian kernel was performed using ρ=0.1, ρ=1 and ρ=10. Only the Gaussian kernel model with the better fit according to AIC was displayed in the table. Of note, according to AIC to compare Gaussian and linear kernel, the linear kernel is the best choice for fitting these data. Mean derivatives divided by standard deviation of the covariate of interest may be interpreted as a beta regression coefficient when assessing the model with linear kernel. When Gaussian kernel is performed, this is just indicative.

| **Annotation score** | **Type of CNVs** | **Models excluding carriers of recurrent CNVs (n=23,484)** | | | **Models excluding carriers of ID-gene CNVs (n=23,967)** | | | **Models excluding carriers of recurrent or ID-gene CNVs (n=23,416)** | | |
| --- | --- | --- | --- | --- | --- | --- | --- | --- | --- | --- |
|  |  | **Est.** | **SE** | ***P*** | **Est.** | **SE** | ***P*** | **Est.** | **SE** | ***P*** |
| **pLI** | **Del.** | -0.173 | 0.023 | 1.98x10^-14^ | -0.157 | 0.020 | 1.13 x10^-14^ | -0.113 | 0.027 | 3.48 x10^-05^ |
|  | **Dup.** | -0.061 | 0.012 | 1.11 x10^-7^ | -0.037 | 0.011 | 5.16 x10^-4^ | -0.047 | 0.013 | 3.32 x10^-04^ |
| **1/LOEUF** | **Del.** | -0.028 | 0.004 | 2.18 x10^-14^ | -0.026 | 0.003 | 4.02 x10^-14^ | -0.018 | 0.005 | 6.70 x10^-05^ |
|  | **Dup.** | -0.010 | 0.002 | 1.13 x10^-7^ | -0.006 | 0.002 | 5.81 x10^-4^ | -0.008 | 0.002 | 2.43 x10^-04^ |

**Supplementary Table 11.** Linear regression models performed in the mega-analysis to measure the effect of deleted or duplicated units of pLI or 1/LOEUF on general intelligence for 3 sensitivity analyses based on exclusion of individuals carrying recurrent CNV or CNV containing ID-gene.

For each sensitivity analysis, the z-scored measure of general intelligence (adjusted) was the dependent variable and annotation score (pLI or 1/LOEUF) for deletions and duplications were the independent ones. All models were adjusted on “type of test-cohort” variable as fixed effect and on familial relationship as random effect. Est.: estimates; SE: standard error; *P*: p-value; pLI: probability of loss-of-function intolerance; LOEUF: the upper 95th CI boundary of the ratio of observed over expected number of loss-of-function mutations.

| **Gene categories** | **Model based on pLI** | | | **Model based on 1/LOEUF** | | |
| --- | --- | --- | --- | --- | --- | --- |
|  | **Est.** | **SE** | ***P*** | **Est.** | **SE** | ***P*** |
| **Deletions Non ID-gene** | -0.142 | 0.018 | 1.27x10^-14^ | -0.023 | 0.003 | 4.33 x10^-15^ |
| **Deletions ID-gene** | -1.025 | 0.243 | 2.56x10^-5^ | -0.174 | 0.035 | 9.24 x10^-7^ |
| **Duplications Non ID-gene** | -0.043 | 0.009 | 4.93x10^-6^ | -0.007 | 0.002 | 2.00 x10^-6^ |
| **Duplications ID-gene** | -0.650 | 0.172 | 1.58x10^-4^ | -0.076 | 0.026 | 3.69 x10^-3^ |

**Supplementary Table 12.** Linear regression models performed in the mega-analysis to measure the effect of deleted or duplicated units of pLI or 1/LOEUF, by gene category (ID- and non ID-genes) on general intelligence.

For both models, the z-scored measure of general intelligence (adjusted) was the dependent variable and annotation score (pLI or 1/LOEUF) computed by category of gene (ID-genes and non ID-genes) and by type of CNV (deletions and duplications) were the independent ones. Models were adjusted on “type of test-cohort” variable as fixed effect and on familial relationship as random effect. Est.: estimates, SE: standard error; P: p-value; pLI: probability of loss-of-function intolerance. LOEUF: the upper 95th CI boundary of the ratio of observed over expected number of loss-of-function mutations

| **Score** | **Tolerance to pLoF** | **ID-gene** | **gene (n=)** | **Score value** | | | **Point of IQ** | | | | | |
| --- | --- | --- | --- | --- | --- | --- | --- | --- | --- | --- | --- | --- |
|  |  |  |  |  |  |  | **Effect size for Del.** | | | **Effect size for Dup.** | | |
|  |  |  |  | **Mean** | **Min.** | **Max.** | **Mean** | **Min.** | **Max.** | **Mean** | **Min.** | **Max.** |
| **1/LOEUF** | **Tolerances score LOEUF ≥0.35** | **No** | 15644 | 1.14 | 0.50 | 2.86 | -0.39 | -0.17 | -0.99 | -0.12 | -0.05 | -0.30 |
|  |  | **Yes** | 65 | 1.66 | 0.53 | 2.86 | -4.33 | -1.38 | -7.46 | -1.89 | -0.60 | -3.26 |
|  | **Intolerance**  **Score LOEUF <0.35** | **No** | 2575 | 5.07 | 2.86 | 33.33 | -1.75 | -0.99 | -11.50 | -0.53 | -0.30 | -3.50 |
|  |  | **Yes** | 167 | 7.73 | 2.93 | 27.03 | -20.18 | -7.65 | -70.54 | -8.81 | -3.34 | -30.81 |
| **pLI** | **Tolerance**  **Score pLI ≤0.9** | **No** | 14432 | 0.15 | 5.4x10^-91^ | 0.90 | -0.32 | -1.1x10^-90^ | -1.92 | -0.10 | -3.5x10^-91^ | -0.58 |
|  |  | **Yes** | 51 | 0.30 | 7.7x10^-45^ | 0.90 | -4.63 | -1.2x10^-43^ | -13.83 | -2.93 | -7.5x10^-44^ | -8.77 |
|  | **Intolerance**  **Score pLI >0.9** | **No** | 2833 | 0.98 | 0.90 | 1 | -2.09 | -1.92 | -2.13 | -0.63 | -0.58 | -0.65 |
|  |  | **Yes** | 167 | 0.99 | 0.90 | 1 | -15.24 | -13.90 | -15.38 | -9.66 | -8.82 | -9.75 |

**Supplementary Table 13.** Distribution of the effect associated with deletion or duplication of genes by gene categories.

Estimates are based on models including effect of annotation score (pLI or 1/LOEUF) by gene category (ID- or non ID-genes) and type of CNV (Table S11). pLI: probability of being loss-of-function intolerant; LOEUF: the upper 95th CI boundary of the ratio of observed over expected number of loss-of-function mutations; Min. Minimum value; Max. maximum value

| **Name** | **Name detail** | **Chr** | **START** | **STOP** | **TYPE** | **n _Genes_** | **ID gene** | **Effect size of z-score for general intelligence** | | | | **Prediction of effect size** | | **Freq. to be**  ***de novo*** | **Prediction to be *de novo*** | |
| --- | --- | --- | --- | --- | --- | --- | --- | --- | --- | --- | --- | --- | --- | --- | --- | --- |
|  |  |  |  |  |  |  |  | **In UKBB**^56^ | **In Lit.** | **based on** | **Mean** | **pLI and ID-gene** | **1/LOEUF and ID-gene** |  | **pLI and ID-gene** | **LOEUF and ID-gene** |
| 1 | TAR | chr1 | 145390000 | 145810000 | Del. | 15 |  | -0.11 | -0.73 | OR*[3]* | -0.42 | -0.48 | -0.5 | 0.2*[3]* | 0.39 | 0.39 |
| 2 | 1q21.1 | chr1 | 146530000 | 147390000 | Del. | 7 |  | -0.27 | -1.01 | NVIQ*[3]* | -0.64 | -0.24 | -0.21 | 0.231*[3]* | 0.22 | 0.2 |
| 3 | 2q11.2 | chr2 | 96740000 | 97680000 | Del. | 18 |  | -0.13 |  |  | -0.13 | -0.83 | -1.03 |  |  |  |
| 4 | 2q13 | chr2 | 111390000 | 112010000 | Del. | 3 |  | -0.16 |  |  | -0.16 | -0.13 | -0.12 |  |  |  |
| 5 | NRXN1 | chr2 | 50140000 | 51260000 | Del. | 1 |  | -0.19 |  |  | -0.19 | -0.14 | -0.09 |  |  |  |
| 6 | 2q13 (*NPHP1*) | chr2 | 110860000 | 110980000 | Del. | 1 |  | -0.01 |  |  | -0.01 | 0 | -0.02 |  |  |  |
| 7 | 3q29 (*DLG1*) | chr3 | 195734000 | 197340000 | Del. | 21 |  |  | -2.1 | FSIQ*[3]* | -2.1 | -0.93 | -0.97 | 0.84*[3]* | 0.74 | 0.74 |
| 8 | 7q11.23 (William-Beuren) | chr7 | 72722981 | 74155278 | Del. | 23 |  |  | -2.09 | NVIQ*[3]* | -2.09 | -1.31 | -1.11 | 0.97*[3]* | 0.91 | 0.82 |
| 9 | 8p23.1 | chr8 | 8100055 | 11764629 | Del. | 26 |  |  | -2.39 | FSIQ*[3]* | -2.39 | -0.46 | -0.61 | 1*[3]* | 0.37 | 0.47 |
| 10 | 10q11.21q11.23 | chr10 | 49390000 | 51060000 | Del. | 16 | *WDFY4* | -0.18 |  |  | -0.18 | -0.23 | -1.03 |  |  |  |
| 11 | 13q12.12 | chr13 | 23560000 | 24880000 | Del. | 5 |  | -0.16 |  |  | -0.16 | -0.01 | -0.14 |  |  |  |
| 12 | 13q12 (*CRYL1*) | chr13 | 20980000 | 21100000 | Del. | 0^a^ |  | 0.01 |  |  | 0.01 | 0 | 0 |  |  |  |
| 13 | 15q13.3 (BP4-BP5) | chr15 | 30918248 | 32515000 | Del. | 7 |  |  | -1.46 | OR*[3]* | -1.46 | -0.24 | -0.22 | 0.282*[3]* | 0.22 | 0.2 |
| 14 | 15q11.2 | chr15 | 22810000 | 23090000 | Del. | 4 |  | -0.15 | -0.6 | NVIQ*[3]* | -0.38 | -0.24 | -0.2 | 0.091*[3]* | 0.22 | 0.19 |
| 15 | 16p11.2-p12.2 | chr16 | 21605180 | 29315879 | Del. | 67 |  |  | -3.33 | FSIQ | -3.33 | -2.9 | -3.38 | 0.92 | 1 | 1 |
| 16 | 16p13.3 ATR-16 syndrome | chr16 | 60001 | 834372 | Del. | 43 | *CAPN15* |  | -2.25 | FSIQ | -2.25 | -1.81 | -1.59 | 0.94 | 0.98 | 0.96 |
| 17 | 16p11.2 | chr16 | 29650000 | 30200000 | Del. | 27 | *MAPK3* | -0.34 | -1.61 | NVIQ*[3]* | -0.97 | -1.37 | -1.18 | 0.57*[3]* | 0.92 | 0.85 |
| 18 | 16p11.2 distal | chr16 | 28820000 | 29050000 | Del. | 9 |  | -0.41 | -0.81 | OR*[3]* | -0.61 | -0.54 | -0.5 | 0.364*[3]* | 0.43 | 0.38 |
| 19 | 16p13.11 | chr16 | 15510000 | 16290000 | Del. | 6 |  | -0.26 | -0.73 | OR*[3]* | -0.49 | -0.32 | -0.41 | 0.256*[3]* | 0.27 | 0.32 |
| 20 | 16p12.1 | chr16 | 21950000 | 22430000 | Del. | 7 |  | -0.1 | -0.5 | OR*[3]* | -0.3 | -0.08 | -0.2 | 0.039*[3]* | 0.14 | 0.19 |
| 21 | 17p11.2 (Smith-Magenis) | chr17 | 17000000 | 21450000 | Del. | 58 | *RAI1* |  | -2.93 | NVIQ*[3]* | -2.93 | -3.03 | -3.26 | 0.956*[3]* | 1 | 1 |
| 22 | 17q12 | chr17 | 34815551 | 36249430 | Del. | 15 |  |  | -0.77 | FSIQ*[3]* | -0.77 | -0.72 | -0.87 | 0.696*[3]* | 0.58 | 0.68 |
| 23 | 17q21.31 | chr17 | 43700000 | 44340000 | Del. | 5 | *KANSL1* |  | -3.4 | FSIQ*[3]* | -3.4 | -1.17 | -0.83 | 0.978*[3]* | 0.86 | 0.65 |
| 24 | *NF1*-microdeletion syndrome | chr17 | 29107097 | 30263321 | Del. | 13 |  |  | -1.54 | FSIQ | -1.54 | -0.85 | -0.6 | 0.882 | 0.68 | 0.47 |
| 25 | 17p12 (*HNPP*) | chr17 | 14140000 | 15430000 | Del. | 4 |  | 0.01 | 0 | OR*[3]* | 0.01 | -0.16 | -0.12 | 0.083*[3]* | 0.18 | 0.15 |
| 26 | 22q11.2 | chr22 | 18893541 | 21901736 | Del. | 49 |  |  | -1.9 | FSIQ | -1.9 | -1.73 | -1.92 | 0.89 | 0.98 | 0.98 |
| 27 | TAR | chr1 | 145390000 | 145810000 | Dup. | 15 |  | -0.12 |  |  | -0.12 | -0.15 | -0.16 |  |  |  |
| 28 | 1q21.1 | chr1 | 146530000 | 147390000 | Dup. | 7 |  | -0.08 |  |  | -0.08 | -0.07 | -0.07 | 0.227 | 0.08 | 0.08 |
| 29 | 2q21.1 | chr2 | 131480000 | 131930000 | Dup. | 4 |  | 0.07 |  |  | 0.07 | -0.01 | -0.04 |  |  |  |
| 30 | 2q13 | chr2 | 111390000 | 112010000 | Dup. | 3 |  | -0.02 |  |  | -0.02 | -0.04 | -0.04 |  |  |  |
| 31 | 2q13 (*NPHP1*) | chr2 | 110860000 | 110980000 | Dup. | 1 |  | 0.01 |  |  | 0.01 | 0 | -0.01 |  |  |  |
| 32 | 7q11.23 | chr7 | 72700000 | 74100000 | Dup. | 24 |  |  | -0.93 | NVIQ | -0.93 | -0.4 | -0.36 | 0.613 | 0.21 | 0.2 |
| 33 | 10q11.21q11.23 | chr10 | 49390000 | 51060000 | Dup. | 16 | *WDFY4* | 0.03 |  |  | 0.03 | -0.07 | -0.41 |  |  |  |
| 34 | 13q12.12 | chr13 | 23560000 | 24880000 | Dup. | 5 |  | -0.01 |  |  | -0.01 | 0 | -0.04 |  |  |  |
| 35 | 15q11q13 (BP3-BP4) | chr15 | 29160000 | 30380000 | Dup. | 3 |  | -0.18 |  |  | -0.18 | -0.05 | -0.12 |  |  |  |
| 36 | 15q11.2 | chr15 | 22810000 | 23090000 | Dup. | 4 |  | -0.03 | -0.05 | NVIQ | -0.04 | -0.07 | -0.06 | 0.022 | 0.08 | 0.08 |
| 37 | 15q13.3 | chr15 | 31080000 | 32460000 | Dup. | 5 |  | -0.09 |  |  | -0.09 | -0.07 | -0.06 |  |  |  |
| 38 | 15q13.3 (*CHRNA7*) | chr15 | 32020000 | 32460000 | Dup. | 0^b^ |  | 0.01 |  |  | 0.01 | 0 | 0 |  |  |  |
| 39 | 16p11.2 | chr16 | 29650000 | 30200000 | Dup. | 27 | *MAPK3* | -0.41 | -0.81 | NVIQ | -0.61 | -0.57 | -0.42 | 0.232 | 0.32 | 0.23 |
| 40 | 16p11.2 distal | chr16 | 28820000 | 29050000 | Dup. | 9 |  | -0.2 |  |  | -0.2 | -0.16 | -0.16 |  |  |  |
| 41 | 16p13.11 | chr16 | 15510000 | 16290000 | Dup. | 6 |  | -0.06 | -1.09 | NVIQ | -0.58 | -0.1 | -0.13 | 0.049 | 0.09 | 0.1 |
| 42 | 16p12.1 | chr16 | 21950000 | 22430000 | Dup. | 7 |  | -0.09 |  |  | -0.09 | -0.02 | -0.06 |  |  |  |
| 43 | 17p11.2 | chr17 | 17000000 | 21400000 | Dup. | 58 | *RAI1* |  | -3.28 | NVIQ | -3.28 | -1.29 | -1.25 | 0.857 | 0.83 | 0.84 |
| 44 | 17q12 (*HNF1B*) | chr17 | 34810000 | 36220000 | Dup. | 15 |  | -0.18 | -0.704 | NVIQ | -0.44 | -0.22 | -0.28 | 0.1 | 0.13 | 0.16 |
| 45 | 17p12 (*CMT1A*) | chr17 | 14140000 | 15430000 | Dup. | 4 |  | -0.04 |  |  | -0.04 | -0.05 | -0.04 |  |  |  |
| 46 | Trisomic 21 | chr21 | 1 | 48100000 | Dup. | 222 | *DYRK1A, KCNJ6, SON, DSCAM* |  | -3.3 | FSIQ | -3.3 | -4.62 | -3.68 | 1 | 1 | 1 |
| 47 | 22q11.2 | chr22 | 19040000 | 21470000 | Dup. | 42 |  | -0.32 | -1.51 | FSIQ | -0.91 | -0.45 | -0.55 | 0.923 | 0.24 | 0.32 |

**Supplementary Table 14.** Description of the effect of 47 recurrent CNVs on general intelligence and probability of being *de novo*, estimated using empirical data from the literature and/or UKBB and using our models.

Most of these CNV are described in Kendall *et al.* 2019*[54]* and in Huguet *et al.* 2018*[3]*. The others are described in table S15. IQ loss was estimated form OR according to the method presented in Huguet *et al.* 2018*[3]*. The *de novo* frequency is defined by Decipher or extracted from Huguet *et al.* 2018*[3]*. a) partially *CRYL1*; b) partially *OTUD7A* and *CHRNA7*. Chr: chromosome, START: start of CNV, STOP: stop of CNV, TYPE: deletion (Del.) or duplication (Dup.), Lit.: Literature, Freq.: Frequency, OR: odds-ratio, FSIQ: Full Scale Intelligence Quotient.

| **Name** | **Name detail** | **Loci** | **IQ References** | **IQ test used** | **n= cases** | **FSIQ loss mean** | **VIQ loss mean** | **NVIQ loss mean** | **Final**  **z-score value** |
| --- | --- | --- | --- | --- | --- | --- | --- | --- | --- |
| 15 | 16p11.2-p12.2 | chr16:21605180-29315879 (Del.) | Hempel et al 2009*[59]* | - | 6 | 50 | - | - | -3.33 ^a^ |
| 16 | 16p13.3 ATR-16 syndrome | chr16:60001-834372 (Del.) | Milone et al 2016*[60]* | Leiter-R | 1 | 32 | - | - | -2.25 ^a^ |
|  |  |  | Wilkie et al 1990*[61]* | - | 8 | 34 | - | - |  |
|  |  |  | Tam et al 2013*[62]* | WISC IV | 1 | 16 | - | - |  |
|  |  |  | Gibson et al 2008*[63]* | - | 4 | 38.25 | - | - |  |
|  |  |  | **Weighted mean** | | 14 | 33.79 | - | - |  |
| 24 | *NF1*-microdeletion syndrome | chr17:29107097-30263321 (Del.) | Mautner et al 2010*[64]* | - | 21 | 23.1 | - | - | -1.54 ^a^ |
| 32 | 7q11.23 | chr7:72700000-74100000 (Dup.) | Mervis et al 2015*[65]* | DAS-II | 63 | 17.95 | 15.86 | 13.36 | -0.93 ^b^ |
|  |  |  | Sanders et al 2011*[66]* | - | 4 | 16 | - | - |  |
|  |  |  | Berg et al 2007*[67]* | WISC-III, WPPSI-III, BSID-III, DAS | 4 | - | 35.25 | 22.25 |  |
|  |  |  | Castiglia et al 2018*[68]* | WISC-III, K-ABC, CFT-20 | 10 | 41.5 | - | - |  |
|  |  |  | **Weighted mean** | | 81 | 20.91 | 17.02 | 13.89 |  |
| 36 | 15q11.2 | chr15:22810000-23090000 (Dup.) | Stefansson et al 2014*[40]* | WASI-I | 136 | - | 0.9 | 0.75 | -0.05 ^b^ |
| 39 | 16p11.2 | chr16:29650000-30200000 (Dup.) | D'Angelo et al 2016*[69]*  (USA adult carrier cohort) | WASI | 39 to 40 | 6.8 | 5.1 | 8.2 | -0.55 ^b^ |
| 41 | 16p13.11 | chr16:15510000-16290000 (Dup.) | Stefansson et al 2014^1^ | WASI-I | 22 | - | 10.95 | 16.35 | -1.09 ^b^ |
|  |  |  | Siu et al 2016*[70]* | WAIS-III | 3 | 10.67 | - | - |  |
|  |  |  | Ullman et al 2007*[71]* | - | 2 | 37 | - | - |  |
|  |  |  | **Weighted mean** | | 27 | 21.2 | 10.95 | 16.35 |  |
| 43 | 17p11.2 | chr17:17000000-21400000 (Dup.) | Treadwell-Deering et al 2010*[72]* | MSEL, SB-IV, SB-V, WISC-III | 14 | 47.64 | - | - | -3.28 ^b^ |
|  |  |  | Potocki et al 2007*[73]* | SB-IV, MSEL | 7 | 51 | 48 | 49 |  |
|  |  |  | Greco et al 2008*[74]* | WISC-III, Leiter-R | 2 to 3 | 52 | 47 | 50 |  |
|  |  |  | **Weighted mean** | | 10 to 14 | 49.17 | 47.78 | 49.22 |  |
| 44 | 17q12 (*HNF1B*) | chr17:34810000-36220000 (Dup.) | Stefansson et al 2014*[40]* | WASI-I | 7 | 12 | 6 | 6.5 | -0.70 ^b^ |
|  |  |  | Verhoeven et al 2017*[75]* | WAIS-III | 1 | 49 | 52 | 39 |  |
|  |  |  | **Weighted mean** | | 8 | 16.63 | 11.75 | 10.56 |  |
| 46 | Trisomic 21 | chr21:1-48100000 (Dup.) | Devenny et al 2000*[76]* | WISC-R | 44 | 46,89 | - | - | -3.30 ^b^ |
|  |  |  | Nicham et al 2003*[77]* | HAWIK-III, HAWIE-R | 20 | 53.15 | 45.33 | 51 |  |
|  |  |  | Capone et al 2005*[78]* | SB-IV | 33 | 59.4 | - | - |  |
|  |  |  | Breia et al 2014*[79]* | WAIS-III | 26 | 50.35 | 47.73 | 49.23 |  |
|  |  |  | Breslin et al 2014*[80]* | KBIT-II | 12 | 51.08 | 45.58 | 47.33 |  |
|  |  |  | **Weighted mean** | | 135 | 51.91 | 46.46 | 49.45 |  |
| 47 | 22q11.2 | chr22:19040000-21470000 (Dup.) | Courtens et al 2008*[81]* | - | 2 | 17.5 | 16.5 | 13.5 | -1.51 ^a^ |
|  |  |  | Rochebrochard et al 2006*[82]* | WAIS-III | 1 | 28 | 29 | 24 |  |
|  |  |  | Van Copenhaut et al 2012*[83]* | BSID, WISC-III | 8 | 23.19 | - | - |  |
|  |  |  | **Weighted mean** | | 11 | 22.6 | 20.67 | 17 |  |

**Supplementary Table 15.** Empirical data on recurrent CNVs.

a) value based on FSIQ; b) value based on NVIQ. FSIQ: Full Scale Intelligence Quotient; VIQ: Verbal Intelligence Quotient, NVIQ: Non Verbal Intelligence Quotient.

| **Cohort** | **n _ind._** | **N CNVs** | | **N *de novo* CNVs** | | **N non-exonic CNVs** | | **N ID-gene CNVs** | | **N CNVs with a 1/LOEUF ≥ 1/0.35** | | **N CNVs with a 1/LOEUF = 0** | | **N CNVs *de novo* with a**  **1/LOEUF = 0** | |
| --- | --- | --- | --- | --- | --- | --- | --- | --- | --- | --- | --- | --- | --- | --- | --- |
|  |  | **Del.** | **Dup.** | **Del.** | **Dup.** | **Del.** | **Dup.** | **Del.** | **Dup.** | **Del.** | **Dup.** | **Del.** | **Dup.** | **Del.** | **Dup.** |
| **DECIPHER** | 10,126 | 6,500 | 5,670 | 3,451 | 1,418 | 1,580 | 1,273 | 1,341 | 784 | 4,246 | 3,258 | 1,592 | 1,300 | 284 | 106 |
| **G-Scot** | 810 | 610 | 740 | 5 | 15 | 422 | 221 | 2 | 9 | 24 | 104 | 514 | 386 | 3 | 6 |
| **MSSNG** | 956 | 797 | 1,074 | 17 | 19 | 533 | 360 | 7 | 22 | 26 | 89 | 665 | 629 | 5 | 5 |
| **SSC probands** | 2,053 | 1,966 | 2,182 | 51 | 101 | 1,311 | 711 | 28 | 58 | 109 | 275 | 1,630 | 1,254 | 24 | 17 |
| **SSC siblings** | 1,866 | 1,723 | 1,949 | 53 | 6 | 1,187 | 655 | 3 | 47 | 51 | 201 | 1,457 | 1,143 | 15 | 13 |
| **Ste-Justine UHC** | 1,560 | 924 | 852 | 403 | 140 | 205 | 189 | 156 | 130 | 645 | 524 | 205 | 194 | 34 | 7 |
| **SYS** | 723 | 594 | 856 | 6 | 8 | 385 | 307 | 4 | 2 | 32 | 121 | 513 | 451 | 3 | 4 |
| **Total** | 18,094 | 13,114 | 13,323 | 3,986 | 1,707 | 5,623 | 3,716 | 1,541 | 1,052 | 5,133 | 4,572 | 6,576 | 5,357 | 368 | 158 |

**Supplementary Table 16.** Detailed table of content for the de novo analysis using Decipher, Sainte-Justine UHC, MSSNG, SSC, Imagen, Generation Scotland, and SYS cohorts.

N: Number; CNVs: Copy number variants; DEL: Deletions; DUP: Duplications; G-Scot: generation Scotland; SSC: Simon simplex collection; Ste Justine: Sainte-Justine UHC cytogenetic database; SYS: Saguenay youth study; LOEUF : the upper 95th CI boundary of the ratio of observed over expected number of LoF mutations.

| **Type of Y value** | **Type of X value** | | **Del.** | | | **Dup.** | | |
| --- | --- | --- | --- | --- | --- | --- | --- | --- |
|  |  |  | **proba (%)** | **95%CI** | ***P*** | **proba (%)** | **95%CI** | ***P*** |
| **Per point of intolerance score** | pLI | **for all genes** | 18.24 | [17.46-19.05] | <1.00x10^-314^ | 8.10 | [7.62-8.61] | 1.78x10^-270^ |
|  |  | **for ID-genes** | 49.14 | [43.22-55.09] | <1.00x10^-314^ | 17.32 | [14.36-20.73] | 1.51x10^-25^ |
|  |  | **for non-ID-genes** | 17.22 | [16.46-18.01] | <1.00x10^-314^ | 7.84 | [7.26-8.35] | 6.30x10^-169^ |
| **Per point of NVIQ computed**  **with intolerance score** |  | **for all genes** | 13.86 | [13.18-14.56] | <1.00x10^-314^ | 8.58 | [8.08-9.11] | 1.78x10^-270^ |
|  |  | **adjusted for ID-genes** | 14.12 | [13.44-14.83] | <1.00x10^-314^ | 8.56 | [8.06-9.08] | 5.77x10-^263^ |
|  |  | **adjusted for ID-genes without recurrent CNVs** | 13.75 | [13.05-14.49] | <1.00x10^-314^ | 8.33 | [7.82-8.86] | 2.015x10^-228^ |
| **Per point of intolerance score** | 1/LOEUF | **for all genes** | 12.09 | [11.45-12.75] | <1.00x10^-314^ | 6.62 | [6.19-7.09] | 9.02x10^-260^ |
|  |  | **for ID-genes** | 14.67 | [13.80-15.58] | 1.94x10^-65^ | 7.21 | [6.71-7.74] | 4.63x10^-24^ |
|  |  | **for non-ID-genes** | 11.7 | [11.07-12.36] | <1.00x10^-314^ | 6.59 | [6.15-7.05] | 2.59x10^-182^ |
| **Per point of NVIQ computed**  **with intolerance score** |  | **for all genes** | 13.34 | [12.67-14.04] | <1.00x10^-314^ | 8.57 | [8.07-9.10] | 9.02x10^-260^ |
|  |  | **adjusted for ID-genes** | 13.49 | [12.82-14.19] | <1.00x10^-314^ | 8.56 | [8.07-9.09] | 1.66x10^-238^ |
|  |  | **adjusted for ID-genes without recurrent CNVs** | 13.12 | [12.43-13.84] | <1.00x10^-314^ | 8.35 | [7.85-8.88] | 2.01x10^-204^ |

**Supplementary Table 17.** Results of the probability of being de novo in function of pLI and 1/LOEUF.

Models were computed independently for deletions and duplications with pLI or 1/LOEUF score as explanatory variables. The status of *de novo* or inherited variant was the dependent variable. Estimates of the probability of being *de novo* (proba), 95% of confidence interval (95%CI) and p-values were computed using a logistic regression. DEL: Deletions; DUP: Duplications; pLI: probability of loss-of-function intolerance; LOEUF : the upper 95th CI boundary of the ratio of observed over expected number of LoF mutations.

| **Variable** | **Model with deletion** | | | **Model with duplication** | | |
| --- | --- | --- | --- | --- | --- | --- |
|  | **Est.** | **SE** | ***P*** | **Est.** | **SE** | ***P*** |
| **nb of highly intolerant genes (**LOEUF <0.2) | -0.378 | 0.102 | 2.20x10^-04^ | 0.015 | 0.054 | 0.779 |
| **nb of moderately intolerant genes (**0.2≤LOEUF<0.35) | -0.25 | 0.066 | 1.48x10^-04^ | -0.098 | 0.031 | 0.002 |
| **nb of tolerant genes**  **(**0.35≤LOEUF<1) | -0.039 | 0.013 | 0.002 | -0.009 | 0.008 | 0.261 |
| **nb of genes highly tolerant**  **(**LOEUF≥1) | 0.011 | 0.011 | 0.316 | -0.006 | 0.005 | 0.275 |

**Supplementary Table 18.** Estimated effects of deleted or duplicated individual genes on general intelligence according to 4 categories of tolerance to pLOF defined by LOEUF.

The model was computed separately for deletion or duplication. The z-scored measure of general intelligence (adjusted) was the dependent variable and number of genes within each category were the 4 independent ones. All models were adjusted on “type of test-cohort” variable as fixed effect and on familial relationship as random effect. Est.: estimate; SE: standard error; *P*: p-value. LOEUF : the upper 95th CI boundary of the ratio of observed over expected number of LoF mutations.

| **Window Interval** | **Selection** | **Deletions** | | | | **Duplications** | | | |
| --- | --- | --- | --- | --- | --- | --- | --- | --- | --- |
|  |  | **Est. (SE)** | ***P*** | ***P* adj** | **Sum of genes** | **Est.** | ***P*** | ***P* adj** | **Sum of genes** |
| [0;0.15] | in the window | -0.739 (0.13) | 1.35x10^-08^ | 3.78x10^-07^ | 68 | -0.042 (0.074) | 5.71x10^-01^ | 8.68x10^-01^ | 233 |
|  | out of the window | -0.027 (0.005) | 1.07x10^-08^ | 1.70x10^-08^ | 6317 | -0.012 (0.002) | 1.21x10^-06^ | 5.10x10^-06^ | 22010 |
| [0.05;0.2] | in the window | -0.488 (0.097) | 4.96x10^-07^ | 6.28x10^-06^ | 100 | -0.022 (0.052) | 6.69x10^-01^ | 8.70x10^-01^ | 387 |
|  | out of the window | -0.026 (0.005) | 9.91x10^-08^ | 1.34x10^-07^ | 6285 | -0.012 (0.003) | 2.79x10^-06^ | 8.83x10^-06^ | 21856 |
| [0.1;0.25] | in the window | -0.38 (0.083) | 4.32x10^-06^ | 2.73x10^-05^ | 165 | -0.09 (0.036) | 1.32x10^-02^ | 1.25x10^-01^ | 670 |
|  | out of the window | -0.022 (0.006) | 1.77x10^-04^ | 1.82x10^-04^ | 6220 | -0.008 (0.003) | 3.60x10^-03^ | 3.90x10^-03^ | 21573 |
| [0.15;0.3] | in the window | -0.372 (0.078) | 2.16x10^-06^ | 2.05x10^-05^ | 158 | -0.112 (0.034) | 1.12x10^-03^ | 2.13x10^-02^ | 695 |
|  | out of the window | -0.023 (0.006) | 4.01x10^-05^ | 4.23x10^-05^ | 6227 | -0.007 (0.003) | 8.80x10^-03^ | 9.29x10^-03^ | 21548 |
| [0.2;0.35] | in the window | -0.347 (0.062) | 1.99x10^-08^ | 3.78x10^-07^ | 324 | -0.097 (0.029) | 9.25x10^-04^ | 2.13x10^-02^ | 1028 |
|  | out of the window | -0.017 (0.006) | 5.27x10^-03^ | 5.27x10^-03^ | 6061 | -0.007 (0.003) | 2.46x10^-02^ | 2.46x10^-02^ | 21215 |
| [0.25;0.4] | in the window | -0.204 (0.044) | 4.12x10^-06^ | 2.73x10^-05^ | 454 | -0.031 (0.027) | 2.55x10^-01^ | 6.06x10^-01^ | 1182 |
|  | out of the window | -0.025 (0.005) | 4.29x10^-06^ | 4.94x10^-06^ | 5931 | -0.011 (0.003) | 1.60x10^-04^ | 2.34x10^-04^ | 21061 |
| [0.3;0.45] | in the window | -0.153 (0.04) | 1.32x10^-04^ | 7.15x10^-04^ | 547 | -0.015 (0.025) | 5.51x10^-01^ | 8.68x10^-01^ | 1331 |
|  | out of the window | -0.027 (0.006) | 3.15x10^-06^ | 3.74x10^-06^ | 5838 | -0.012 (0.003) | 1.07x10^-04^ | 1.73x10^-04^ | 20912 |
| [0.35;0.5] | in the window | -0.128 (0.049) | 9.16x10^-03^ | 2.49x10^-02^ | 480 | 0.005 (0.027) | 8.63x10^-01^ | 9.37x10^-01^ | 1173 |
|  | out of the window | -0.03 (0.007) | 6.20x10^-06^ | 6.93x10^-06^ | 5905 | -0.014 (0.003) | 2.02x10^-05^ | 4.04x10^-05^ | 21070 |
| [0.4;0.55] | in the window | -0.081 (0.045) | 7.08x10^-02^ | 1.12x10^-01^ | 434 | -0.038 (0.028) | 1.73x10^-01^ | 5.27x10^-01^ | 1170 |
|  | out of the window | -0.034 (0.006) | 3.73x10^-08^ | 5.45x10^-08^ | 5951 | -0.01 (0.003) | 1.60x10^-03^ | 2.03x10^-03^ | 21073 |
| [0.45;0.6] | in the window | -0.02 (0.047) | 6.67x10^-01^ | 7.04x10^-01^ | 406 | -0.041 (0.028) | 1.45x10^-01^ | 5.02x10^-01^ | 1212 |
|  | out of the window | -0.041 (0.006) | 9.36x10^-11^ | 1.87x10^-10^ | 5979 | -0.01 (0.003) | 2.09x10^-03^ | 2.56x10^-03^ | 21031 |
| [0.5;0.65] | in the window | -0.044 (0.041) | 2.79x10^-01^ | 3.31x10^-01^ | 389 | -0.067 (0.025) | 8.75x10^-03^ | 1.11x10^-01^ | 1282 |
|  | out of the window | -0.038 (0.006) | 3.40x10^-11^ | 8.08x10^-11^ | 5996 | -0.008 (0.003) | 1.63x10^-02^ | 1.67x10^-02^ | 20961 |
| [0.55;0.7] | in the window | -0.067 (0.04) | 9.86x10^-02^ | 1.42x10^-01^ | 581 | -0.05 (0.023) | 2.81x10^-02^ | 1.78x10^-01^ | 2038 |
|  | out of the window | -0.036 (0.006) | 8.20x10^-10^ | 1.53x10^-09^ | 5804 | -0.009 (0.003) | 3.03x10^-03^ | 3.39x10^-03^ | 20205 |
| [0.6;0.75] | in the window | -0.104 (0.039) | 8.21x10^-03^ | 2.40x10^-02^ | 596 | -0.045 (0.023) | 4.62x10^-02^ | 2.20x10^-01^ | 2126 |
|  | out of the window | -0.032 (0.006) | 2.75x10^-08^ | 4.17x10^-08^ | 5789 | -0.009 (0.003) | 2.39x10^-03^ | 2.84x10^-03^ | 20117 |
| [0.65;0.8] | in the window | -0.115 (0.04) | 3.86x10^-03^ | 1.47x10^-02^ | 631 | -0.026 (0.02) | 1.95x10^-01^ | 5.29x10^-01^ | 2469 |
|  | out of the window | -0.03 (0.006) | 4.25x10^-07^ | 5.39x10^-07^ | 5754 | -0.011 (0.003) | 3.74x10^-04^ | 5.07x10^-04^ | 19774 |
| [0.7;0.85] | in the window | -0.169 (0.049) | 4.91x10^-04^ | 2.33x10^-03^ | 447 | 0.001 (0.024) | 9.65x10^-01^ | 9.65x10^-01^ | 1814 |
|  | out of the window | -0.026 (0.006) | 3.80x10^-05^ | 4.13x10^-05^ | 5938 | -0.014 (0.003) | 3.08x10^-05^ | 5.86x10^-05^ | 20429 |
| [0.75;0.9] | in the window | -0.117 (0.044) | 8.08x10^-03^ | 2.40x10^-02^ | 536 | -0.001 (0.023) | 9.60x10^-01^ | 9.65x10^-01^ | 1890 |
|  | out of the window | -0.03 (0.006) | 2.65x10^-06^ | 3.25x10^-06^ | 5849 | -0.013 (0.003) | 1.09x10^-04^ | 1.73x10^-04^ | 20353 |
| [0.8;0.95] | in the window | -0.077 (0.042) | 6.48x10^-02^ | 1.12x10^-01^ | 534 | 0.005 (0.024) | 8.34x10^-01^ | 9.32x10^-01^ | 1698 |
|  | out of the window | -0.035 (0.006) | 6.39x10^-09^ | 1.06x10^-08^ | 5851 | -0.014 (0.003) | 3.63x10^-05^ | 6.56x10^-05^ | 20545 |
| [0.85;1] | in the window | -0.052 (0.038) | 1.71x10^-01^ | 2.24x10^-01^ | 635 | -0.008 (0.019) | 6.87x10^-01^ | 8.70x10^-01^ | 2095 |
|  | out of the window | -0.037 (0.006) | 8.47x10^-10^ | 1.53x10^-09^ | 5750 | -0.013 (0.003) | 9.99x10^-05^ | 1.72x10^-04^ | 20148 |
| [0.9;1.05] | in the window | -0.089 (0.041) | 2.80x10^-02^ | 5.99x10^-02^ | 754 | 0.014 (0.022) | 5.34x10^-01^ | 8.68x10^-01^ | 2210 |
|  | out of the window | -0.033 (0.006) | 1.14x10^-07^ | 1.49x10^-07^ | 5631 | -0.015 (0.003) | 6.26x10^-06^ | 1.59x10^-05^ | 20033 |
| [0.95;1.1] | in the window | -0.081 (0.038) | 3.36x10^-02^ | 6.72x10^-02^ | 798 | -0.014 (0.021) | 5.12x10^-01^ | 8.68x10^-01^ | 2315 |
|  | out of the window | -0.034 (0.006) | 3.91x10^-09^ | 6.76x10^-09^ | 5587 | -0.012 (0.003) | 2.69x10^-04^ | 3.79x10^-04^ | 19928 |
| [1;1.15] | in the window | -0.136 (0.043) | 1.70x10^-03^ | 7.20x10^-03^ | 659 | -0.025 (0.022) | 2.53x10^-01^ | 6.06x10^-01^ | 1947 |
|  | out of the window | -0.03 (0.006) | 8.84x10^-08^ | 1.24x10^-07^ | 5726 | -0.011 (0.003) | 4.39x10^-04^ | 5.75x10^-04^ | 20296 |
| [1.05;1.2] | in the window | -0.085 (0.041) | 3.75x10^-02^ | 7.12x10^-02^ | 630 | -0.046 (0.02) | 2.40x10^-02^ | 1.78x10^-01^ | 2541 |
|  | out of the window | -0.035 (0.005) | 8.81x10^-11^ | 1.86x10^-10^ | 5755 | -0.009 (0.003) | 2.94x10^-03^ | 3.39x10^-03^ | 19702 |
| [1.1;1.25] | in the window | -0.087 (0.043) | 4.21x10^-02^ | 7.62x10^-02^ | 597 | 0.003 (0.019) | 8.91x10^-01^ | 9.41x10^-01^ | 3358 |
|  | out of the window | -0.035 (0.005) | 6.13x10^-11^ | 1.37x10^-10^ | 5788 | -0.014 (0.003) | 4.12x10^-06^ | 1.20x10^-05^ | 18885 |
| [1.15;1.3] | in the window | 0.031 (0.043) | 4.65x10^-01^ | 5.19x10^-01^ | 629 | 0.009 (0.014) | 5.05x10^-01^ | 8.68x10^-01^ | 4171 |
|  | out of the window | -0.045 (0.005) | 9.75x10^-17^ | 2.47x10^-16^ | 5756 | -0.014 (0.003) | 4.47x10^-08^ | 5.77x10^-07^ | 18072 |
| [1.2;1.35] | in the window | 0.051 (0.046) | 2.67x10^-01^ | 3.27x10^-01^ | 526 | 0.007 (0.014) | 6.26x10^-01^ | 8.70x10^-01^ | 3325 |
|  | out of the window | -0.045 (0.005) | 1.53x10^-18^ | 5.30x10^-18^ | 5859 | -0.014 (0.003) | 6.07x10^-08^ | 5.77x10^-07^ | 18918 |
| [1.25;1.4] | in the window | 0.053 (0.047) | 2.53x10^-01^ | 3.21x10^-01^ | 470 | 0.014 (0.021) | 5.07x10^-01^ | 8.68x10^-01^ | 2259 |
|  | out of the window | -0.045 (0.005) | 3.41x10^-19^ | 1.44x10^-18^ | 5915 | -0.014 (0.003) | 1.57x10^-07^ | 9.93x10^-07^ | 19984 |
| [1.3;1.45] | in the window | 0.035 (0.048) | 4.57x10^-01^ | 5.19x10^-01^ | 416 | 0.016 (0.026) | 5.38x10^-01^ | 8.68x10^-01^ | 1313 |
|  | out of the window | -0.043 (0.005) | 5.89x10^-19^ | 2.24x10^-18^ | 5969 | -0.014 (0.003) | 6.28x10^-07^ | 3.41x10^-06^ | 20930 |
| [1.35;1.5] | in the window | 0.09 (0.049) | 6.80x10^-02^ | 1.12x10^-01^ | 376 | 0.054 (0.027) | 4.41x10^-02^ | 2.20x10^-01^ | 1332 |
|  | out of the window | -0.048 (0.005) | 1.01x10^-19^ | 4.80x10^-19^ | 6009 | -0.017 (0.003) | 8.56x10^-09^ | 3.25x10^-07^ | 20911 |
| [1.4;1.55] | in the window | 0.128 (0.047) | 7.09x10^-03^ | 2.40x10^-02^ | 441 | 0.043 (0.029) | 1.35x10^-01^ | 5.02x10^-01^ | 1283 |
|  | out of the window | -0.052 (0.006) | 5.03x10^-21^ | 3.19x10^-20^ | 5944 | -0.016 (0.003) | 1.08x10^-07^ | 8.18x10^-07^ | 20960 |
| [1.45;1.6] | in the window | 0.121 (0.049) | 1.39x10^-02^ | 3.51x10^-02^ | 431 | 0.043 (0.027) | 1.15x10^-01^ | 4.88x10^-01^ | 1233 |
|  | out of the window | -0.051 (0.005) | 1.41x10^-20^ | 7.65x10^-20^ | 5954 | -0.016 (0.003) | 5.03x10^-08^ | 5.77x10^-07^ | 21010 |
| [1.5;1.65] | in the window | 0.11 (0.048) | 2.14x10^-02^ | 5.08x10^-02^ | 414 | 0.005 (0.024) | 8.25x10^-01^ | 9.32x10^-01^ | 1349 |
|  | out of the window | -0.049 (0.005) | 1.50x10^-21^ | 1.96x10^-20^ | 5971 | -0.014 (0.003) | 2.57x10^-06^ | 8.83x10^-06^ | 20894 |
| [1.55;1.7] | in the window | 0.065 (0.044) | 1.38x10^-01^ | 1.88x10^-01^ | 411 | -0.01 (0.024) | 6.79x10^-01^ | 8.70x10^-01^ | 1504 |
|  | out of the window | -0.045 (0.005) | 3.47x10^-21^ | 2.64x10^-20^ | 5974 | -0.012 (0.003) | 1.36x10^-05^ | 2.86x10^-05^ | 20739 |
| [1.6;1.75] | in the window | 0.012 (0.04) | 7.64x10^-01^ | 7.85x10^-01^ | 485 | -0.033 (0.024) | 1.80x10^-01^ | 5.27x10^-01^ | 1529 |
|  | out of the window | -0.042 (0.005) | 4.18x10^-18^ | 1.32x10^-17^ | 5900 | -0.011 (0.003) | 1.36x10^-04^ | 2.07x10^-04^ | 20714 |
| [1.65;1.8] | in the window | -0.002 (0.04) | 9.61x10^-01^ | 9.61x10^-01^ | 478 | -0.006 (0.022) | 7.99x10^-01^ | 9.32x10^-01^ | 1438 |
|  | out of the window | -0.041 (0.005) | 1.74x10^-17^ | 5.10x10^-17^ | 5907 | -0.013 (0.003) | 2.03x10^-06^ | 7.73x10^-06^ | 20805 |
| [1.7;1.85] | in the window | -0.018 (0.036) | 6.28x10^-01^ | 6.82x10^-01^ | 612 | -0.009 (0.021) | 6.44x10^-01^ | 8.70x10^-01^ | 1438 |
|  | out of the window | -0.04 (0.005) | 3.11x10^-17^ | 8.43x10^-17^ | 5773 | -0.012 (0.003) | 5.29x10^-06^ | 1.44x10^-05^ | 20805 |
| [1.75;1.9] | in the window | 0.057 (0.034) | 9.19x10^-02^ | 1.40x10^-01^ | 661 | 0.004 (0.018) | 8.14x10^-01^ | 9.32x10^-01^ | 1902 |
|  | out of the window | -0.047 (0.005) | 2.16x10^-21^ | 2.05x10^-20^ | 5724 | -0.014 (0.003) | 7.65x10^-07^ | 3.63x10^-06^ | 20341 |
| [1.8;1.95] | in the window | 0.047 (0.029) | 1.01x10^-01^ | 1.42x10^-01^ | 778 | -0.011 (0.015) | 4.44x10^-01^ | 8.68x10^-01^ | 2175 |
|  | out of the window | -0.048 (0.005) | 1.55x10^-21^ | 1.96x10^-20^ | 5607 | -0.012 (0.003) | 7.45x10^-06^ | 1.76x10^-05^ | 20068 |
| [1.85;2] | in the window | 0.071 (0.032) | 2.84x10^-02^ | 5.99x10^-02^ | 598 | -0.013 (0.015) | 3.75x10^-01^ | 8.37x10^-01^ | 2046 |
|  | out of the window | -0.05 (0.005) | 2.63x10^-22^ | 1.00x10^-20^ | 5787 | -0.012 (0.003) | 7.89x10^-06^ | 1.76x10^-05^ | 20197 |

**Supplementary Table 19.** Estimated effects of deleted of duplicated individual genes on general intelligence according to several moving categories defined by a sliding window on LOEUF.

We used a sliding window to define 2 categories of genes by LOEUF, one in the window and other out the window. We evaluated the effect sizes of genes within 37 windows with a sliding of 0.05 for each step. The model was computed separately for deletion or duplication. The z-scored measure of general intelligence (adjusted) was the dependent variable number of gene in and out of the window were the 2 independent ones. All models were adjusted on “type of test-cohort” variable as fixed effect and on familial relationship as random effect. Est.: estimate; SE: standard error; *P*: p-value. LOEUF : the upper 95th CI boundary of the ratio of observed over expected number of LoF mutations.

**Supplementary Table 20.** DAVID enrichment details with all genes within CNV for each population.

**Supplementary Table 21.** DAVID enrichment details with intolerant genes in the genome.

**Supplementary Table 22.** DAVID enrichment details with tolerant genes in the genome.

**Supplementary Table 23.** DAVID enrichment details with intolerant and tolerant genes within CNV for each population.

| **Cohort** | **Correlation^(a)^ (95%CI)** | **ICC^(b)^ (95%CI)** |
| --- | --- | --- |
| **Imagen (n=1,720)** | 0.77 (0.75;0.78) | 0.77 (0.74;0.78) |
| **SYS-child (n=857)** | 0.61 (0.56;0.65) | 0.60 (0.56;0.64) |
| **LBC1936 (n=504)** | 0.66 (0.61;0.71) | 0.66 (0.62;0.70) |

**Supplementary Table 24.** Correlation and concordance between NVIQ and g-factor in 3 unselected populations.

(a) Pearson’s correlation, (b) Intraclass correlation coefficient, Single fixed raters; CI: confidence interval

### Extended Data Figures

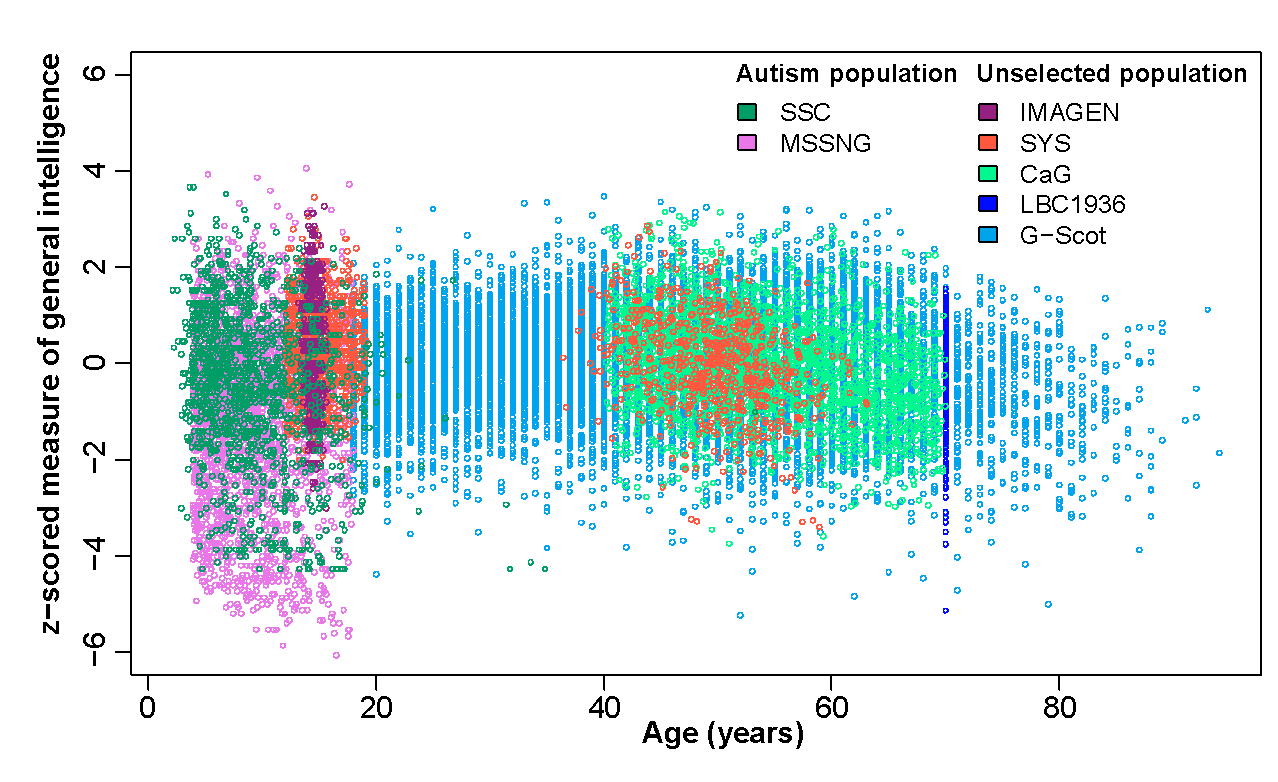

**Supplementary Fig. 1.** Distribution of z-scored general intelligence measured either by NVIQ or g-factor according to the age of individuals and colored by cohort.

Z-scored measures of general intelligence were obtained by z-scoring the NVIQ with a mean of 100 and a standard deviation of 15 and z-scored g-factor was obtained separately for each cohort using mean and standard deviation of the cohort; NVIQ: non-verbal intelligence quotient; g-factor: general factor; SSC: Simon simplex collection; SYS: Saguenay youth study; G-Scot: generation Scotland, CaG: Cartagene; LBC: Lothian birth cohort.

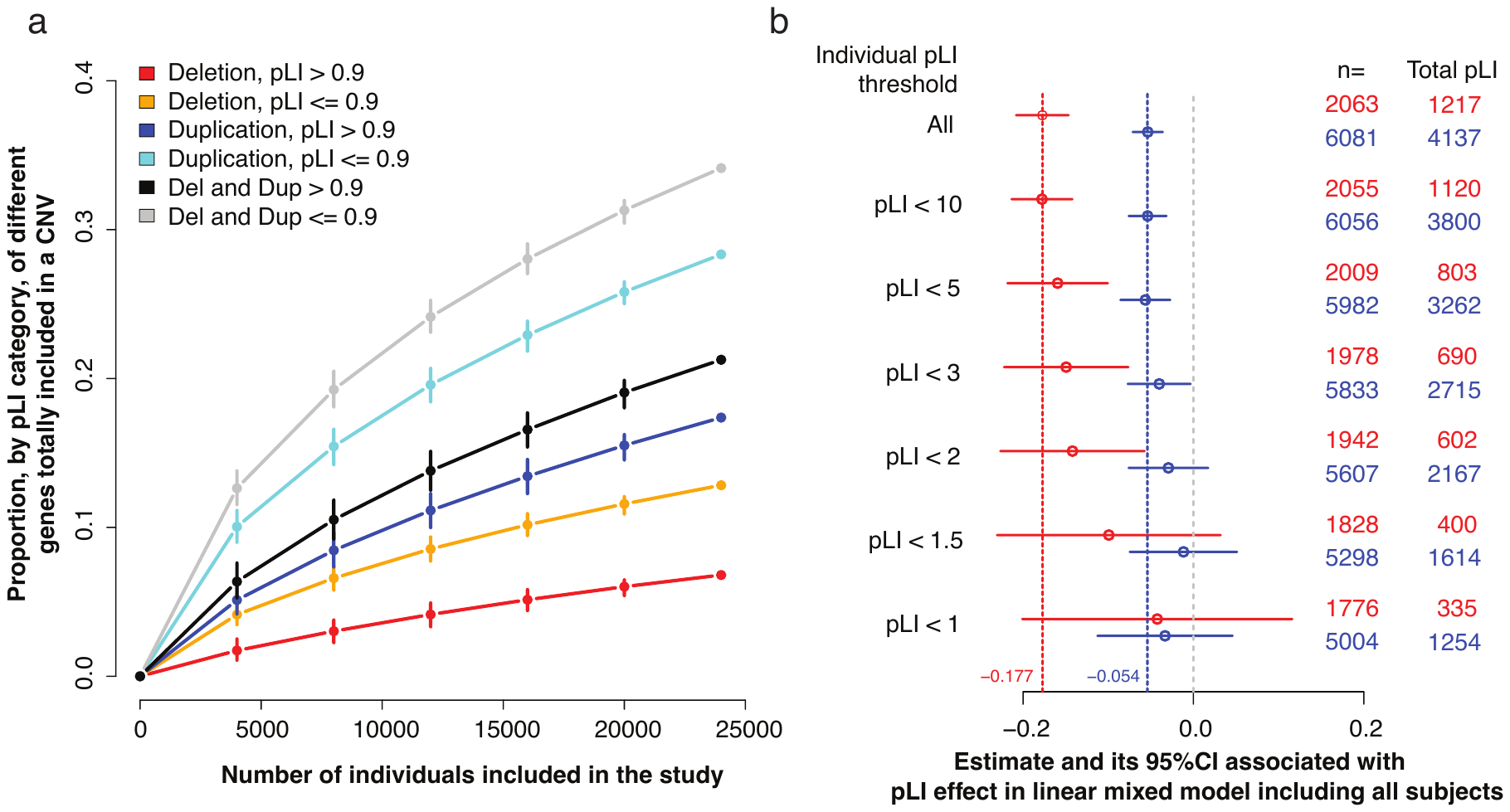

**Supplementary Fig. 2.** Sensitivity analyses for model based on pLI score.

**a.** Estimated proportion of the coding genome, within each category defined by pLI, encompassed in CNVs present in the mega-analysis according to sample size (randomly selected within the mega-analysis). **b.** Estimated effect of pLI on general intelligence after removing individuals with a sum of pLI larger that 10, 5, 3, 2, 1.5 and 1. n: number of individuals with total sum of pLI > 0.

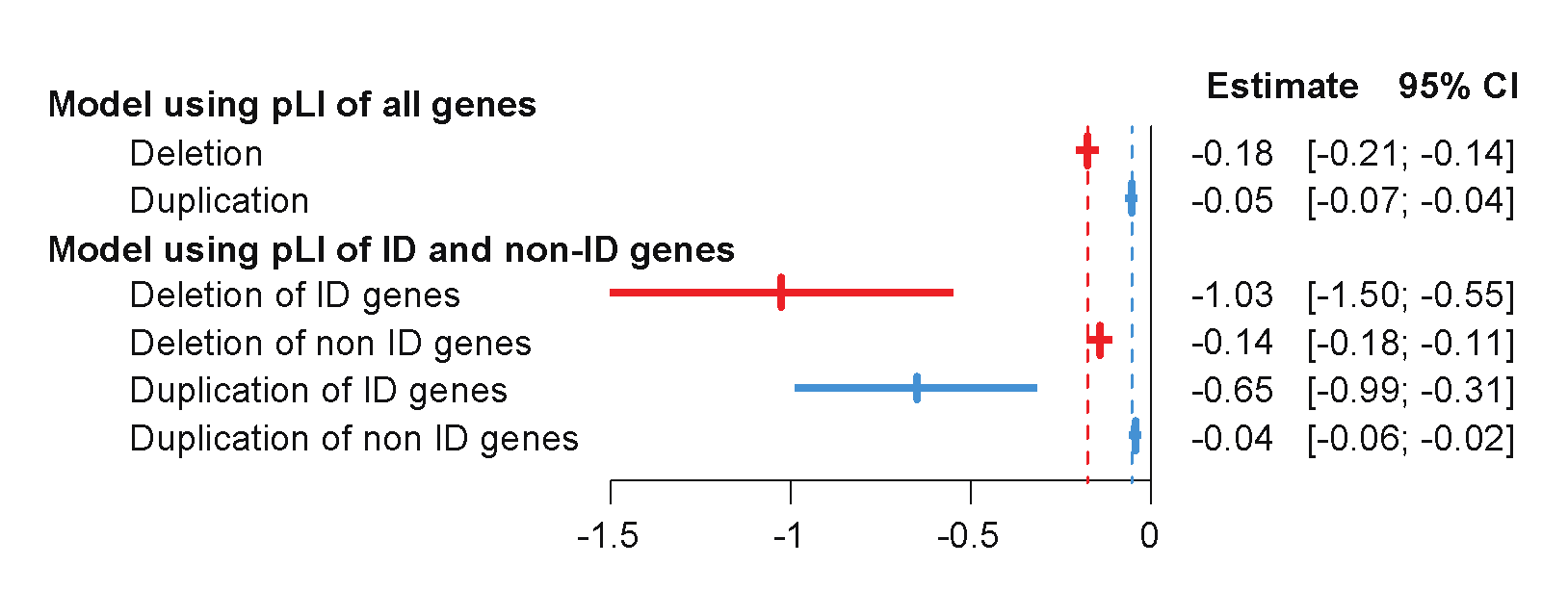

**Supplementary Fig. 3.** Effect size associated with pLI divided by category of intellectual disability (ID) genes on general intelligence.

The z-scored measure of general intelligence (adjusted) was the dependent variable. In the first model, the 2 predictors are the sum of pLI for deletion and the sum of pLI for duplications. In the second model, the 4 predictors are the sum of pLI for ID genes and non-ID genes for deletions and duplication. Both models were adjusted for “type of test-cohort” variable as fixed effect and on familial relationship as random effect. pLI: probability of loss-of-function intolerance; CI: confidence interval.

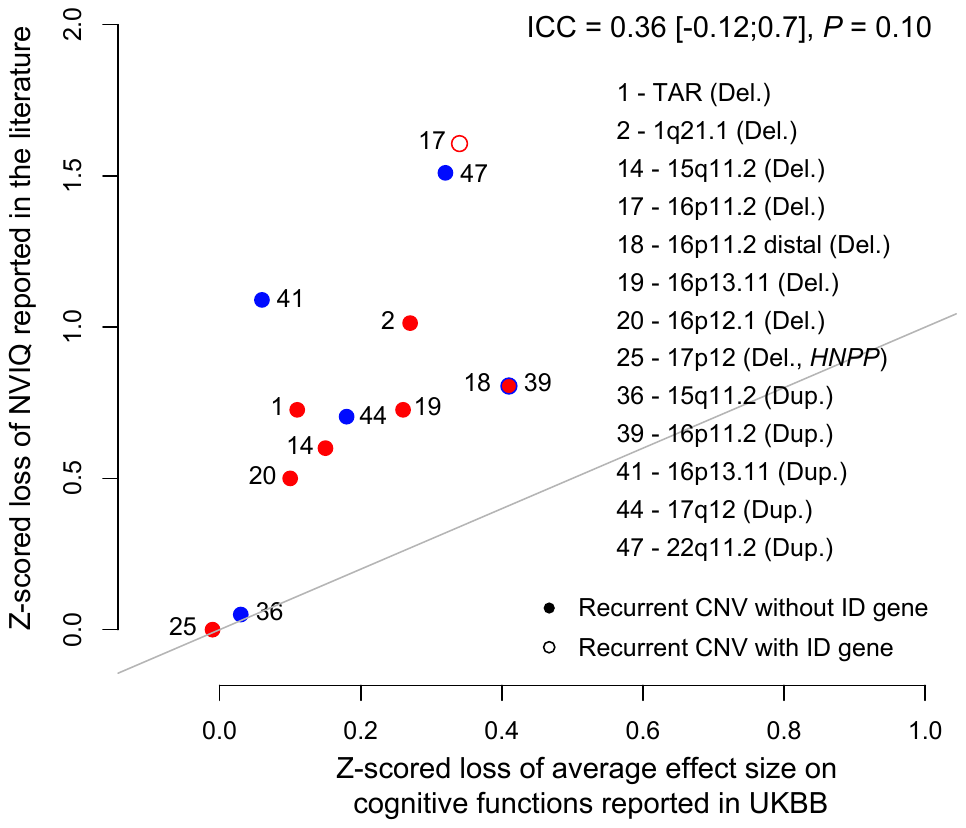

**Supplementary Fig. 4.** Concordance between observation from literature and UKBB for CNV effects on general intelligence.

X and Y values: effect size of CNVs on z-scored general intelligence. Concordance between literature reports for general intelligence loss observed in clinically and UKBB ascertained carriers of 13 recurrent CNVs (Table S14). Each point represents a recurrent CNV, red and blue points are deletions and duplications respectively. Empty circles are CNVs encompassing ID-genes. The model uses 2 explanatory variables (LOEUF of non-ID-genes and ID-genes). ICC indicates intraclass correlation coefficient (3, 1).

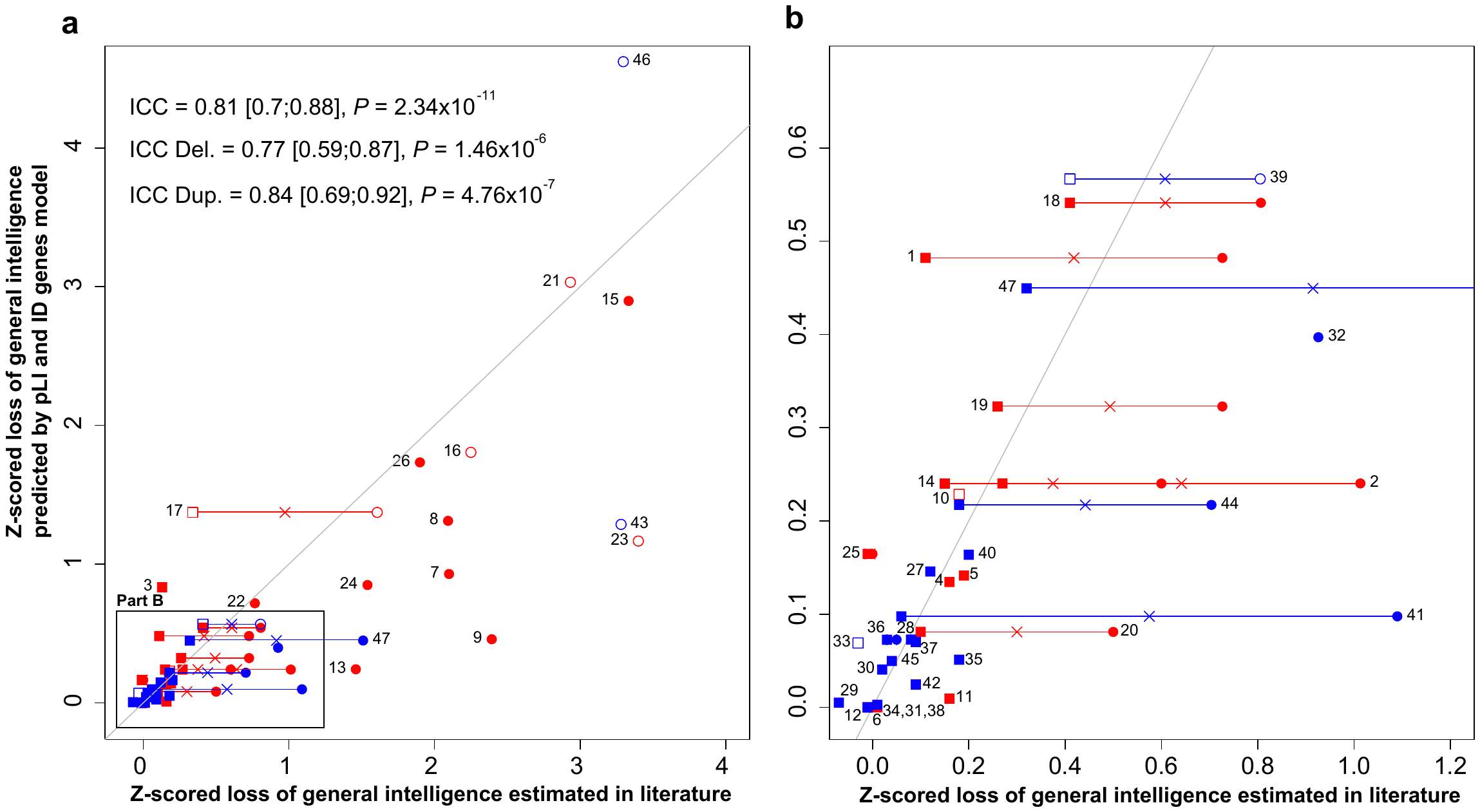

**Supplementary Fig. 5.** Concordance between model predictions and published observations for CNV effects on general intelligence.

**a.** and **b.** Concordance between model estimates (with pLI and ID-genes) and literature of clinical data and UKBB reports for general intelligence loss observed in respectively 27 and 33 recurrent CNVs for a total of ascertained carriers of 47 recurrent CNVs (Table S12). X- and Y-values: effect size of CNVs on z-scored general intelligence. **b.** Zoom of the rectangle drawn in the lower left section of panel **a.** We represented values from clinical data by a circle and those from UKBB data by a square. The cross represents the mean value of z-scored IQ loss for the 13 recurrent CNVs observed both in literature and in UKBB. Deletions are in red and duplications in blue. Empty circles or square are CNVs encompassing ID-genes. The model uses 2 explanatory variables (pLI of non-ID-genes and ID-genes). ICC indicates intraclass correlation coefficient (3, 1). Each point represents a recurrent CNV: (1) TAR Deletion; (2) 1q21.1 Deletion; (3) 2q11.2 Deletion; (4) 2q13 Deletion; (5) *NRXN1* Deletion; (6) 2q13 (*NPHP1*) Deletion; (7) 3q29 (*DLG1*) Deletion; (8) 7q11.23 (William-Beuren) Deletion; (9) 8p23.1 Deletion; (10) 10q11.21q11.23 Deletion; (11) 13q12.12 Deletion; (12) 13q12 (*CRYL1*) Deletion; (13) 15q13.3 (BP4-BP5) Deletion; (14) 15q11.2 Deletion; (15) 16p11.2-p12.2 Deletion; (16) 16p13.3 ATR-16 syndrome Deletion; (17) 16p11.2 Deletion; (18) 16p11.2 distal Deletion; (19) 16p13.11 Deletion; (20) 16p12.1 Deletion; (21) 17p11.2 (Smith-Magenis) Deletion; (22) 17q12 Deletion; (23) 17q21.31 Deletion; (24) NF1-microdeletion syndrome Deletion; (25) 17p12 (*HNPP*) Deletion; (26) 22q11.2 Deletion; (27) TAR Duplication; (28) 1q21.1 Duplication; (29) 2q21.1 Duplication; (30) 2q13 Duplication; (31) 2q13 (*NPHP1*) Duplication; (32) 7q11.23 Duplication; (33) 10q11.21q11.23 Duplication; (34) 13q12.12 Duplication; (35) 15q11q13 (BP3-BP4) Duplication; (36) 15q11.2 Duplication; (37) 15q13.3 Duplication; (38) 15q13.3 (*CHRNA7*) Duplication; (39) 16p11.2 Duplication; (40) 16p11.2 distal Duplication; (41) 16p13.11 Duplication; (42) 16p12.1 Duplication; (43) 17p11.2 Duplication; (44) 17q12 (*HNF1B*) Duplication; (45) 17p12 (*CMT1A*) Duplication; (46) Trisomic 21 Duplication; (47) 22q11.2 Duplication.

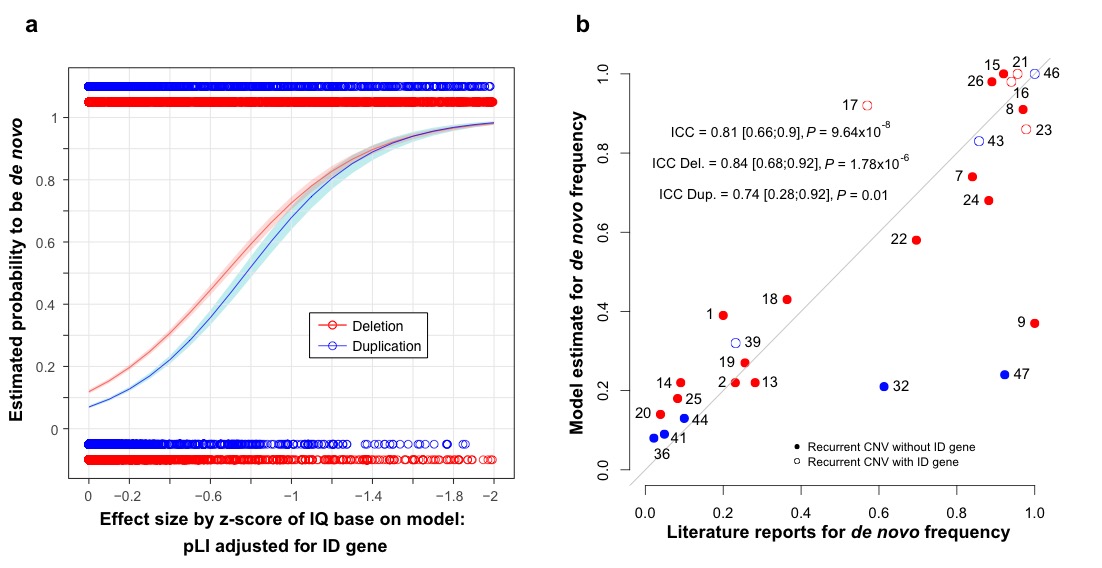

**Supplementary Fig. 6.** Estimated probability of *de novo*, based on model including pLI for ID and non ID genes, and its concordance with *de novo* frequency observed in literature.

**a.** Probability of *de novo* estimated by our *de novo* model (Y-axis) according to the loss of IQ estimated by a model using pLI for ID and non-ID genes as two explanatory variables (X-axis). The *de novo* model was fitted on 13,114 deletions (red) and 13,323 duplications (blue) with available inheritance information observed in DECIPHER, CHU Sainte-Justine, SSC, MSSNG, SYS and G-Scot. **b.** Concordance between *de novo* frequency observed in DECIPHER (X-axis) and the probability of being *de novo* estimated by models when excluding recurrent CNVs of the training dataset (Y-axis) pLI as an explanatory variable for 28 recurrent CNVs. The first bisector represents the perfect concordance. ICC indicates intraclass correlation coefficient (3, 1). Each point corresponds to a known recurrent CNV: (1) TAR Deletion; (2) 1q21.1 Deletion; (7) 3q29 (*DLG1*) Deletion; (8) 7q11.23 (William-Beuren) Deletion; (9) 8p23.1 Deletion; (13) 15q13.3 (BP4-BP5) Deletion; (14) 15q11.2 Deletion; (15) 16p11.2-p12.2 Deletion; (16) 16p13.3 ATR-16 syndrome Deletion; (17) 16p11.2 Deletion; (18) 16p11.2 distal Deletion; (19) 16p13.11 Deletion; (20) 16p12.1 Deletion; (21) 17p11.2 (Smith-Magenis) Deletion; (22) 17q12 Deletion; (23) 17q21.31 Deletion; (24) NF1-microdeletion syndrome Deletion; (25) 17p12 (*HNPP*) Deletion; (26) 22q11.2 Deletion; (32) 7q11.23 Duplication; (36) 15q11.2 Duplication; (39) 16p11.2 Duplication; (41) 16p13.11 Duplication; (43) 17p11.2 Duplication; (44) 17q12 (*HNF1B*) Duplication; (46) Trisomic 21 Duplication; (47) 22q11.2 Duplication.

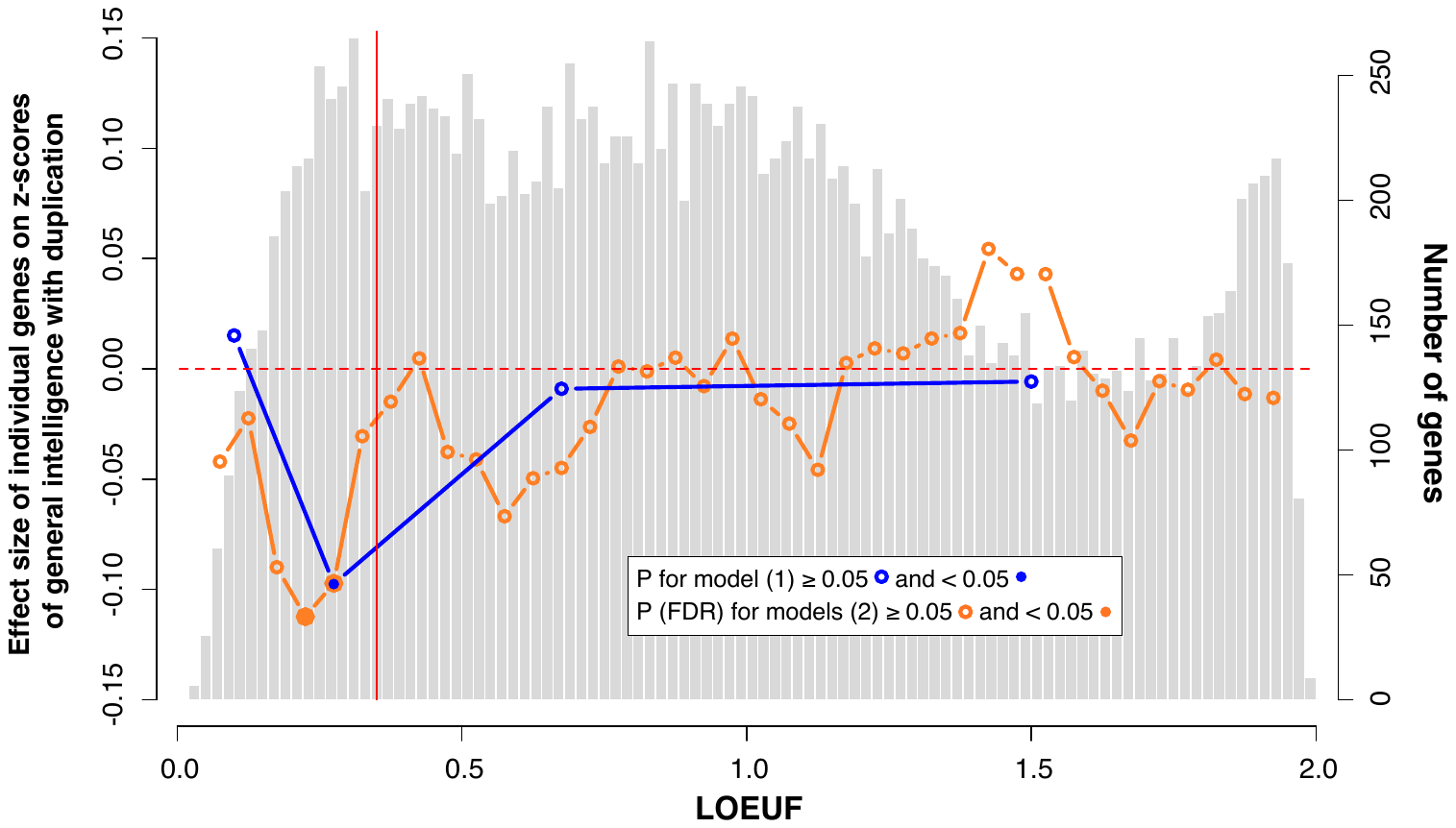

**Supplementary Fig. 7.** Estimated effects of individual genes on general intelligence according to categories based on LOEUF for duplications.

The light grey histogram represents the distribution of LOEUF values for 18,451 autosomal genes. The blue line represents the estimates for a gene in each of the 4 categories of LOEUF included in the model (Supplementary material): highly intolerant genes (LOEUF <0.2, n=980), moderately intolerant genes (0.2≤LOEUF<0.35 n=1,762), tolerant genes (0.35≤LOEUF<1, n=7,442) and genes highly tolerant to pLoF (LOEUF≥1, n=8,267). The orange line represents the estimated effect size of 37 categories of genes based on their LOEUF values (sliding windows=0.15) in the model (Supplementary material). Genes with a LOEUF below 0.35 (vertical red line) are considered to be intolerant to pLoF by gnomAD. Left Y axis values: z-scored general intelligence (1 z-score is equivalent to 15 points of IQ) for duplication. Right Y axis values: number of genes represented in the histogram.

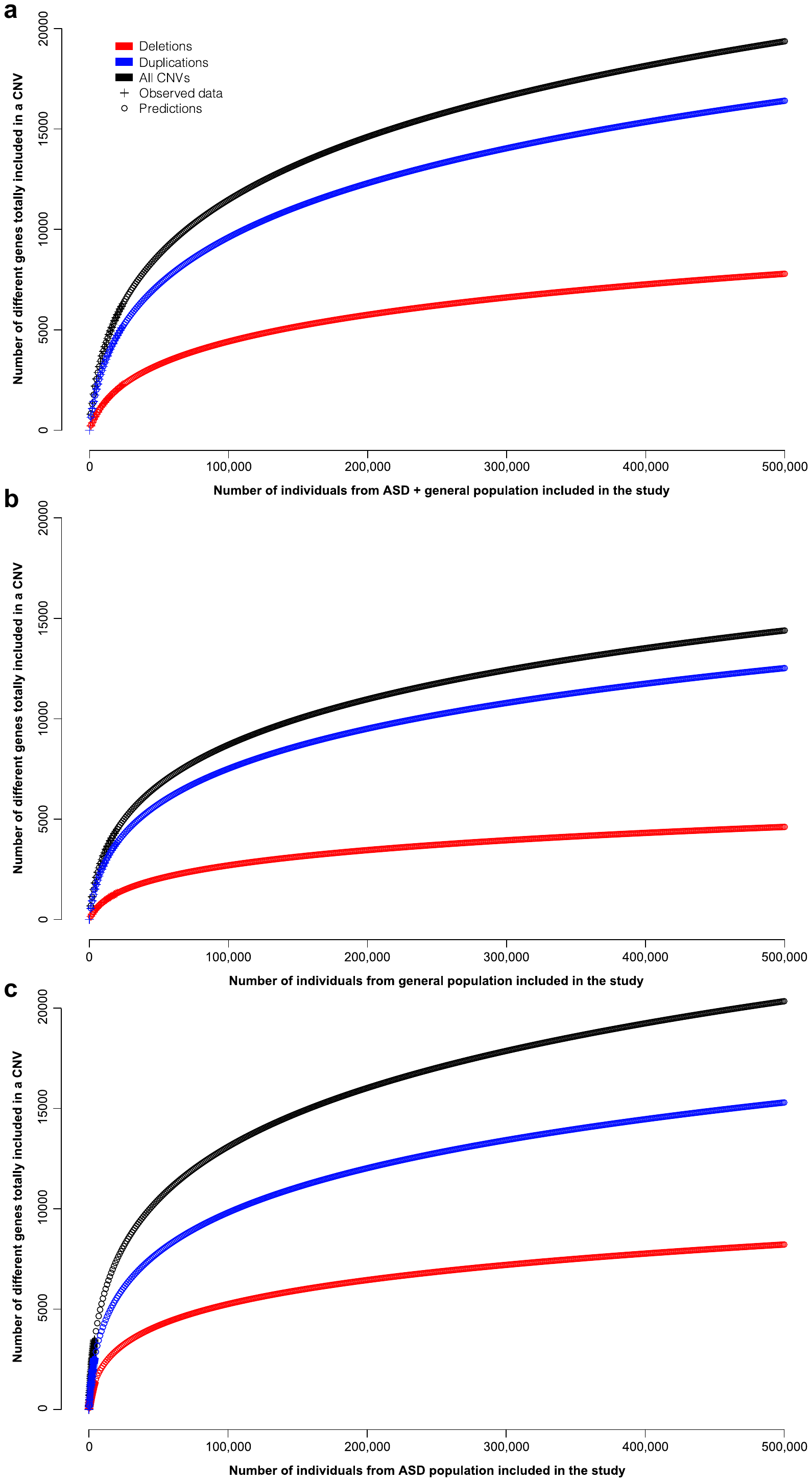

**Supplementary Fig. 8.** Prediction of gene coverage according to the sample size.

**a.** Predictions based on coefficients from regression analysis obtained using gene coverage estimated in the entire cohort (n=24,092).

**b.** Predictions based on coefficients from regression analysis obtained using gene coverage estimated in general population (n=20,151).

**c.** Predictions based on coefficients from regression analysis obtained using gene coverage estimated in ASD population (n=3,941).

CNV: copy number variant; ASD: autism spectrum disorder.

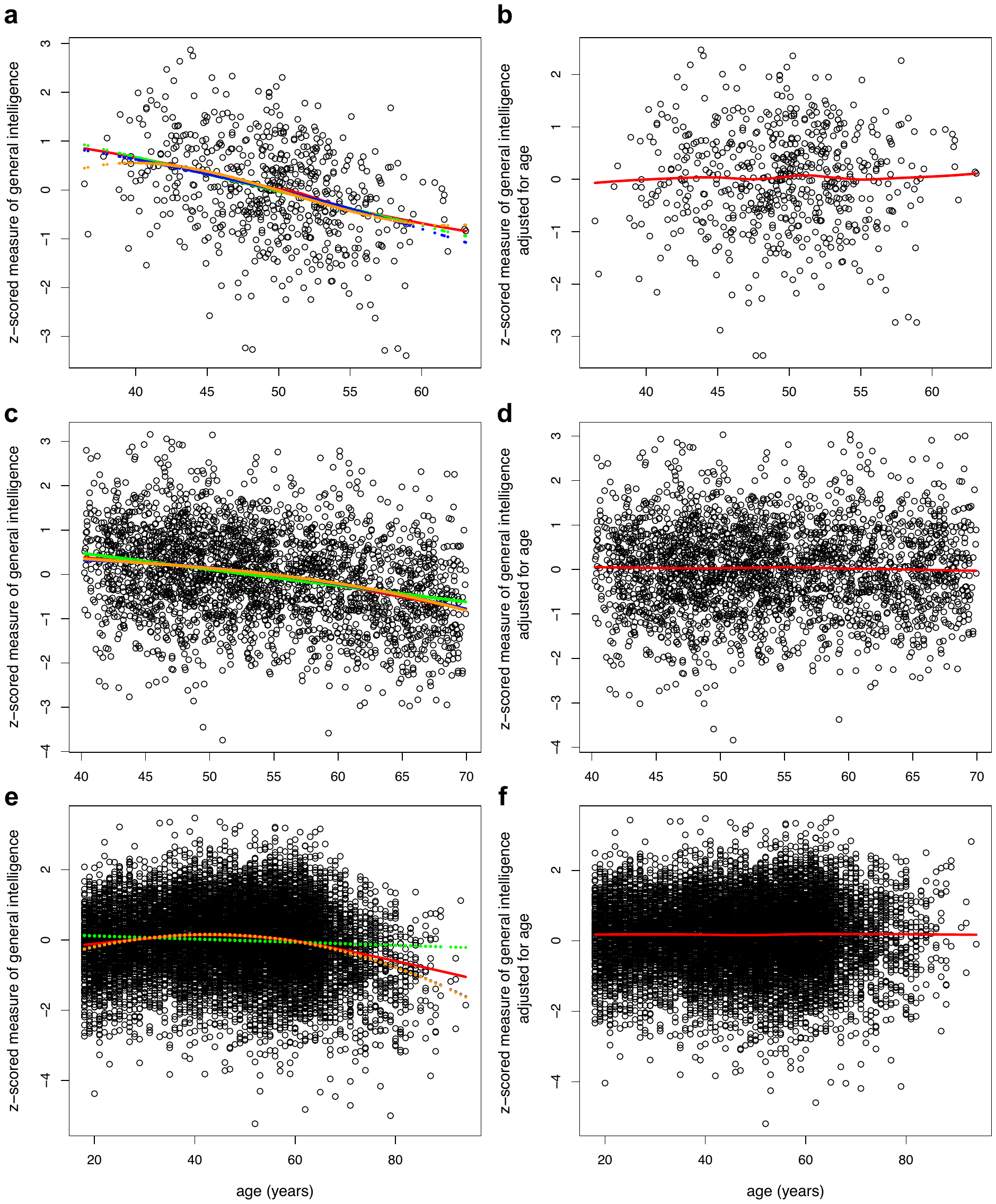

**Supplementary Fig. 9.** Distribution of z-scored measure of general intelligence before (left panels) and after (right panels) adjustment for age (g-factor in general populations).

**a,b**: SYS-parents; **c,d**: CaG; **e,f**: G-Scot. In **a-f** panels, X-axis represents the age of individuals, and Y-axis represents the z-scored measure of general intelligence before (left panels) and after (right panels) age-adjustment. Points were fitted using different models as represented in the legend: “lowess” is the local weighted polynomial regression curve, “linear”: the effect of the age was modeled as linear, “poly. d=2”: the effect of the age was modeled as quadratic; “poly. d=3”: the effect of the age was modeled as cubic. No effect of age shows that the phenotype was correctly adjusted for age. Of note, adjustment was made using quadratic effect of the age in CaG (intercept: Est.=-0.96 [SE=0.98, *P*=0.3260], age: estimate=5.94×10^-3^ [SE=3.00×10^-3^, *P*=0.0480], age²: estimate=-6.82×10^-6^ [SE=2.27×10^-6^, *P*=0.0027]) and G-Scot (intercept: Est.=-1.09 [SE=6.81×10^-2^, *P*=9.85×10^-57^], age: Est.=4.84×10^-3^ [SE=2.56×10^-4^, *P*=8.46×10^-79^], age²: Est.=-4.71×10^-6^ [SE=2.27×10^-7^, *P*=7.18×10^-94^]) and using linear effect of the age in SYS-parents (intercept: Est.=3.48 [SE=0.39, *P*=1.07×10^-17^], age: Est.=-0.01 [SE=6.60×10^-4^, *P*=8.71x10-18]). Est.: estimate; SE: standard error; *P*: p-value.

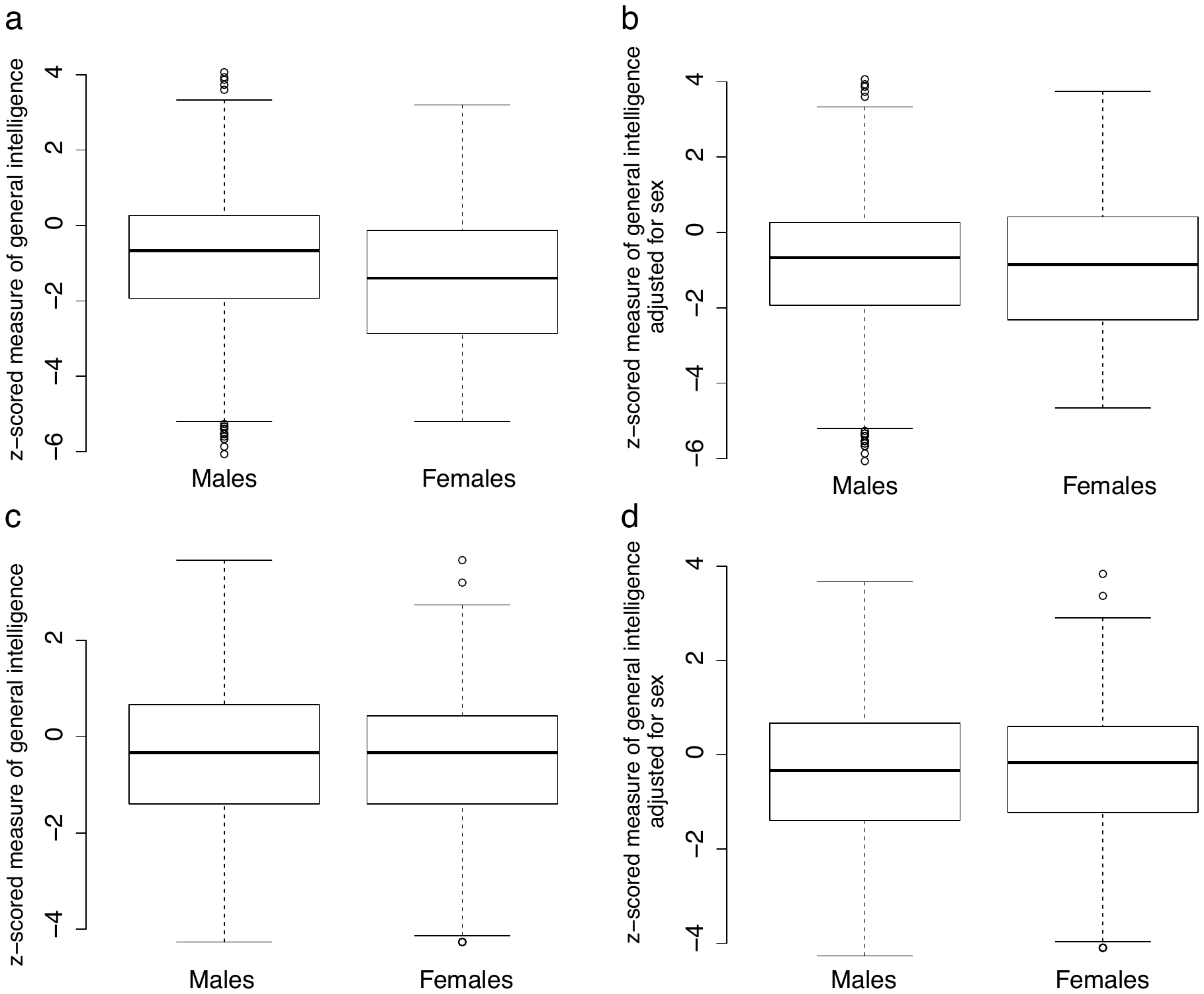

**Supplementary Fig. 10.** Distribution of z-scored measure of general intelligence before (left panels) and after (right panels) adjustment for sex (NVIQ in autism populations).

**a,b:** SSC, **c,d:** MSSNG. In **a-d** panels, boxplots represents the distribution of the z-scored measure of general intelligence before (left panels) and after (right panels) sex-adjustment for males and females in SSC (intercept: Est.=-0.96 [SE=0.04, *P*=1.43×10^-132^], sex_F/M_=-0.55 [SE=0.10, *P*=6.61×10^-8^]) and MSSNG (intercept: Est.=-0.43 [SE=0.05, *P*=3.03×10^-19^], sex_F/M_=-0.17 [SE=0.11, *P*=0.1155]). CaG: Cartagene; G-Scot: generation Scotland; SYS: Saguenay youth study; SSC: Simon simplex collection; NVIQ: non-verbal intelligence quotient; g-factor: general factor. Est.: estimate; SE: standard error; *P*: p-value.

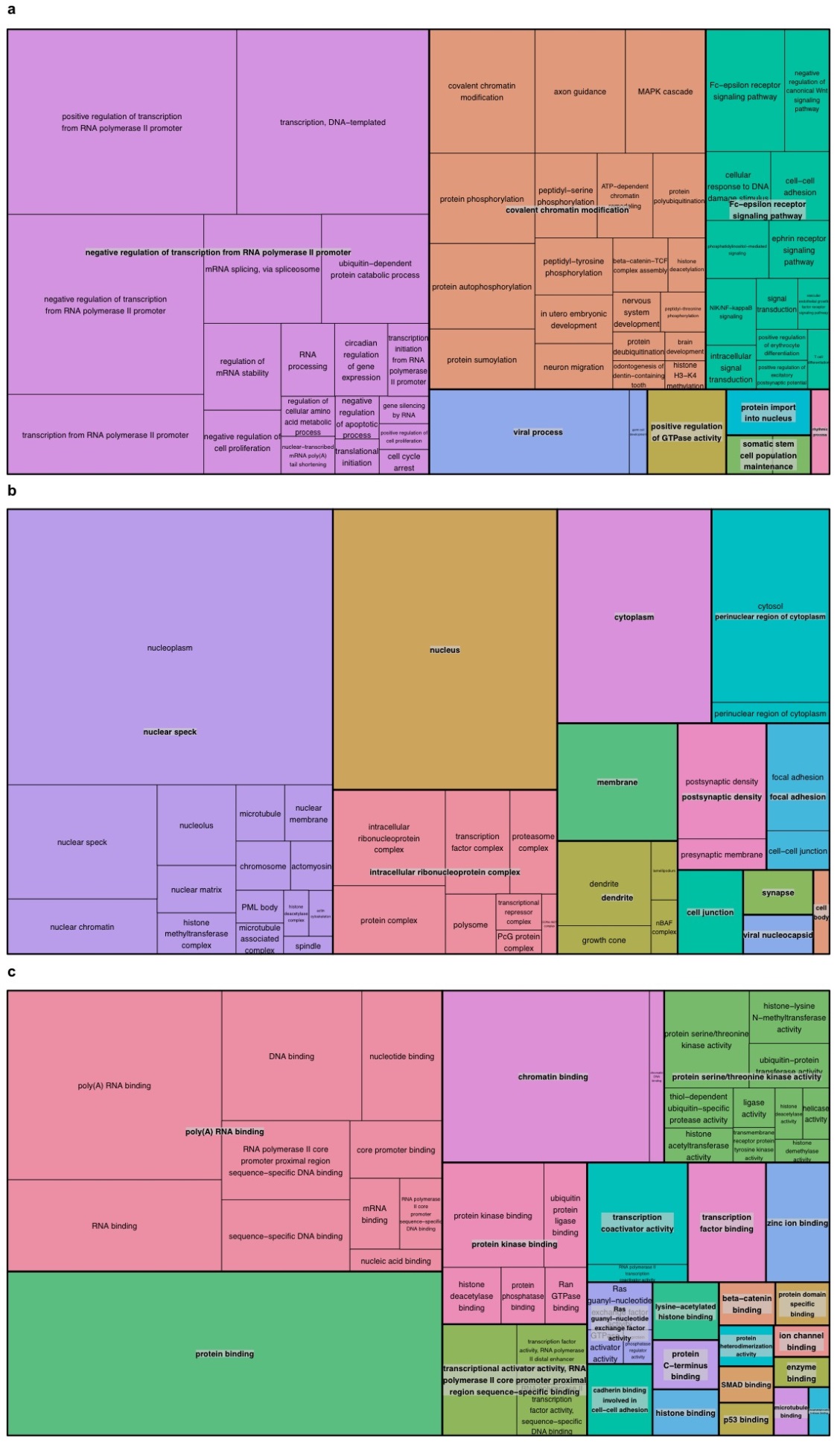

**Supplementary Fig. 11.** TreeMap view of GO-term clusters for intolerant genes in the genome.

The TreeMap was generated using REVIGO with the input of all enriched GOterms for **a** biological process, **b** Cellular component and **c** Molecular Function after Bonferroni correction at 5%. Each rectangle represents a single GO-term cluster. Sizes of the rectangle reflect the -log10(p) distribution. The representatives are joined into ‘superclusters’ of loosely related terms, visualized with different colors.

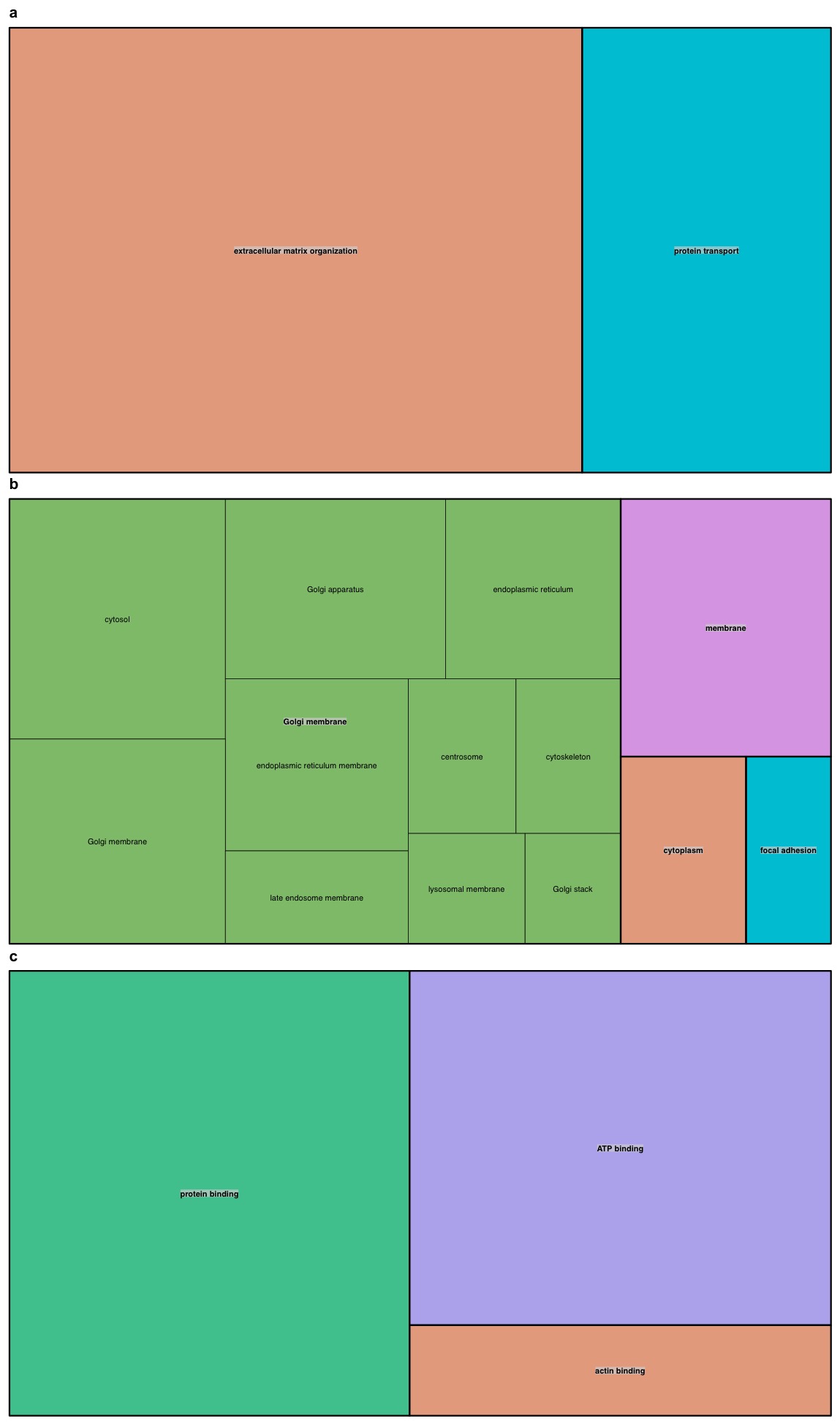

**Supplementary Fig. 12.** TreeMap view of GO-term clusters for tolerant genes in the genome.

The TreeMap was generated using REVIGO with the input of all enriched GOterms for **a** biological process, **b** Cellular component and **c** Molecular Function after Bonferroni correction at 5%. Each rectangle represents a single GO-term cluster. Sizes of the rectangle reflect the -log10(p) distribution. The representatives are joined into ‘superclusters’ of loosely related terms, visualized with different colors.

.
